# Cuticular hydrocarbon divergence in *Drosophila melanogaster* populations evolving under differential operational sex ratios

**DOI:** 10.1101/351718

**Authors:** Rochishnu Dutta, Tejinder Singh Chechi, Ankit Yadav, Nagaraj Guru Prasad

## Abstract

The ability of interlocus sexual conflict to facilitate reproductive isolation is widely anticipated. However, very few experimental evolutionary studies have convincingly demonstrated the evolution of reproductive isolation due to sexual conflict. Recently a study on replicate populations of *Drosophila melanogaster* under differential sexual conflict found that divergent mate preference evolved among replicate populations under high sexual conflict regime. The precopulatory isolating mechanism underlying such divergent mate preference could be sexual signals such as cuticular hydrocarbons since they evolve rapidly and are involved in *D. melanogaster* mate recognition. Using *D. melanogaster* replicates used in the previous study, we investigate whether cuticular hydrocarbon divergence bears signatures of sexually antagonistic coevolution that led to reproductive isolation among replicates of high sexual conflict regime. We found that *D. melanogaster* cuticular hydrocarbon profiles are sexually dimorphic. Although replicate populations under high sexual conflict displayed assortative mating, we found no significant differences in the cuticular hydrocarbon profile between the high and low sexual conflict regimes. Instead we find cuticular hydrocarbon divergence patterns to be suggestive of the Buridan’s Ass regime which is one of the six possible mechanisms to resolve sexual conflict. Sexual selection that co-vary between populations under high and low sexual conflict regimes may also have contributed to the evolution of cuticular hydrocarbons. This study indicates that population differentiation as a result of cuticular hydrocarbon divergence cannot be credited to sexual conflict despite high sexual conflict regime evolving divergent cuticular hydrocarbon profiles.

## Introduction

Mathematical models of interlocus sexual conflict based on mating rate envisage six possible evolutionary outcomes, out of which two lead to speciation: allopatric speciation as a by-product of sexually antagonistic coevolution (SAC) and sympatric speciation as a result of males responding to female strategy under Buridan’s Ass regime (1). SAC occurs when males’ mate harming traits and female traits countering such mate harm continually evolve until a stabilising selection restricts further evolution of traits. Buridan’s Ass regime refers to genetic differentiation of females in response to harm caused by males such that males lose out on fitness due to genetic mismatch. Males may, however, gain advantage by mirroring female genetic diversification that may lead to isolation of divergent genetic groups within a population (2). A comparative study on groups of insect species has shown that the prevalence of high sexual conflict makes a taxonomic group more speciose than those with low or no sexual conflict (3). However, the support for speciation through sexual conflict is not universal and comparative studies on other groups of species have not found a similar trend (4, 5). Experimental studies that directly investigate speciation linked to sexual conflict too are inconsistent in their support for the hypothesis presumably due to the occurrence of four other possible dynamic outcomes predicted in the mathematical models that do not result in speciation (6).

By evolving independent replicates of a dung fly population for 35 generations under monogamous (low) and polygamous (high) sexual conflict regime, Martin and Hosken (7) showed that mating success between individuals of the same population was significantly higher than that of pairs consisting of males and females from different populations under polygamous regime. In the monogamous regime, there was no difference in mating success between individuals from hetero-population and same population. Recently, a study on *Drosophila melanogaster* population using male biased (high sexual conflict) and female biased (low sexual conflict) operational sex ratio regimes showed that evolution of both premating and post mating prezygotic isolation was possible within 105 generations in the high sexual conflict regime but not in the low sexual conflict regime (8). Contrary to these two studies, few other studies were unable to find any support for sexual conflict driven reproductive isolation leading us to question the potential of sexual conflict as an “engine of speciation” (4, 5, 9–11).

Manipulating sex ratio in *D. melanogaster* population in favour of either sex leads to differential sexual conflict. Males of male biased regime evolved higher mate harming ability indicated by higher mortality of ancestral females that they mated with while males from female biased regime were found to be relatively benign to ancestral females (12). Concurrently, females from male biased regime show increased fecundity and longevity when continuously exposed to ancestral males (13). These results indicate that the males from male biased regimes evolved more mate harming abilities and females from the same regimes have evolved resistance to mate harm compared to male and females from female biased population. It is important to note that both male biased and female biased regimes had three independent replicate populations each and as such, these replicate populations could be construed as allopatric populations, each potentially following its own evolutionary trajectory. The allopatric male biased replicates have now shown within replicate assortative mating that indicates reproductive isolation due to SAC (8).

Premating isolation by mate choice in *Drosophila* sp. is often mediated by cuticular hydrocarbon (henceforth CHC) profiles that provide information about species identity, sex, mating status, age, etc. to a receiver (14–16). Specifically, CHCs such as 7-tricosene, 5-tricosene, 9-pentacosene, 7-pentacosene, 7,11-heptacosadiene and 7,11-nonacosadiene play a pivotal role in sexual communication during various heterosexual courtship rituals in *Drosophila melanogaster* (17). Since reproductive isolation in male biased *D*. *melanogaster* replicates is most likely mediated by mate choice rather than any conspicuous female reluctance to mating (8), it is possible that the CHC profiles form the mechanistic basis for mate preference within male biased replicates. The possibility of mate choice forming the basis for premating reproductive isolation under SAC has cropped up recently (8). Until now, it has been suggested that traits undergoing SAC evolve directly due to natural selection on female resistance to mate harm (18). However, this novel idea assumes that the CHC profiles have also differentiated among male biased replicates as a by-product of SAC within each allopatric replicate.

CHC profiles of *Drosophila melanogaster* are dimorphic (17) and have the ability to evolve rapidly (19). Those CHCs that take part in mate preference are known to have independent genetic controls in the two sexes, often linked to sex determination genes, for their biosynthesis (14, 19–21). Many previous studies have implicated differentiated CHC profiles in the evolution of reproductive isolation in *Drosophila* sp. through sexual selection (22–25). It is, therefore, impossible to ignore that the different operational sex ratio may also lead to differential sexual selection in male biased and female biased *D. melanogaster* regime. Male biased population may experience high sexual selection as males face higher mate competition and choosy females. Female biased population may experience lowered sexual selection pressures as males experience less competition for mates and females cannot be as choosy (or may even have to compete for mates). In case, SAC does not affect CHC profiles of male biased replicates (mate attraction being multimodal in *Drosophila* sp. (26)), it is equally possible for us to see the evolutionary outcome of sexual selection in CHC profiles of both male biased and female biased regime. It is important to note that sexual selection and sexual conflict are not mutually exclusive mechanisms through which cuticular hydrocarbon profiles of populations under altered sex ratio could evolve (27). In this study, we investigate the possibility of CHC divergence under selection pressure imposed by different operational sex ratio that may lead to differential sexual conflict or differential sexual selection in *D. melanogaster*.

Our primary objective is to examine the patterns of CHC divergence among the replicates of *D. melanogaster* male biased and female biased regimes, the same population previously used by Syed et al (8). Assuming that CHC profiles (that play a role in mate attraction in *D. melanogaster)* have evolved independently in male biased replicates (isolated by mate choice as indicated by Syed et al (8)) due to SAC, we expect that the CHC profiles of male biased replicates will diverge more than the CHC profiles of the female biased replicates. However, if CHC profiles of male biased regime is not affected by SAC, we may see convergence of CHC profiles of males compared to females due to sexual selection. In male biased regime, males will have to compete for females and can potentially evolve a common CHC profile most preferred by females. The CHC profiles among replicates of female biased regime are not expected to be affected by SAC, unlike male biased regime. We also expect that all *D. melanogaster* populations will show sexual dimorphism in CHC profiles. Our results suggest that reproductive isolation among male biased replicates due to high sexual conflict is not explained by the patterns of CHC divergence. Instead we find sexual selection may predominantly be shaping CHC divergence in *D. melanogaster* populations maintained under different operational sex ratio.

## Methods

### *D. melanogaster* population maintenance

In this study, we used 168-170 generations old *Drosophila melanogaster* MCF population (henceforth MCF) whose adult flies are maintained at three operational sex-ratio. A 3:1::male:female sex ratio (24 males + 8 females per vial) led to male biased regime (henceforth M). Similarly, a 1:3::male:female sex ratio (8 males + 24 females per vial) gave rise to female biased regime (henceforth F). The control regime (henceforth C) had 1:1 sex ratio (16 males + 16 females per vial). Each regime has three replicate populations (M_1,2,3_ in the M regime, F_1,2,3_ in the F regime and C_1,2,3_ in the C regime). All nine populations trace their ancestry to a large, laboratory adapted *D. melanogaster* LHst population (12).

Nandy et al (12, 28) derived three populations, labelled C_1,2,3_, from LHst (a recessive but benign scarlet eyed *D. melanogaster* population named after Lawrence Harshman). The C_1,2,3_ populations were maintained at a 1:1 adult sex ratio for five generations. Subsequently, from C_1,2,3_ populations, we derived M_1,2,3_ and F_1,2,3_ regime with the subscripted numbers indicating common ancestry. For example, M_1_, F_1_ and C_1_ were derived from C_1_. Moreover, during population maintenance, MCF populations with the same subscript are handled on the same day and, therefore, can be treated as statistical blocks.

These populations are maintained on a 14-day discrete generation cycle, at 25°C and 60% relative humidity, 12:12 hours light/dark cycle and standard cornmeal– molasses–yeast food in standard vials (90-mm length × 30-mm diameter). We collect virgin males and females every generation and combine them in appropriate sex ratios for each regime. Males and females interact for around 48 hours, after which they are transferred to fresh vials for oviposition. Eggs laid are trimmed to a density of around 140-160 eggs per vial containing 6-8 ml food. These eggs are used to start the next generation (refer (12) for detailed maintenance protocol).

### Generating flies for cuticular hydrocarbon extraction

Before CHC extraction from virgin MCF flies, we allowed one generation of “standardization” or common-garden rearing at a 1:1 sex ratio (12) to account for any potential non-genetic parental effect (29). In order to generate flies for CHC extraction, we collected eggs from “standardised” flies. On the 10th day after egg collection from the MCF population, we collected newly eclosed flies as virgins within 6 hours of eclosion under light carbon dioxide anesthesia. The flies were isolated by keeping each of them singly in a vial for 2 days. On 12th day (ensuring flies of same age to minimize age-specific variation of CHCs) the CHC extraction assay was conducted.

### Cuticular hydrocarbon extraction

We transferred each virgin individual to a 1.5 ml gas chromatography (GC) vial without anesthetising it. We then poured 200 μl HPLC grade 95% *n*-Hexane solvent (AlfaAesar, 39199) containing 10ng/μl (10μl Pentadecane in 1ml *n*-Hexane) 99% pure Pentadecane (C_15_H_32_, AlfaAesar (A10336), internal standard) into the GC vials. The vials were left undisturbed for 4 minutes and then vortexed for 1 minute (30). We removed the dead individual from each GC vial and subsequently left the *n*-Hexane to dry out. The extracted samples in the GC vials were then stored at 4°C until gas chromatography was performed. We extracted 119 M (M_1_ = 20 males + 20 females, M_2_ = 20 males + 20 females and M_3_ = 20 males + 19 females) flies, 115 F (F_1_ = 19 males + 18 females, F_2_ = 20 males + 19 females and F_3_ = 19 males + 20 females) flies and 118 C (C_1_ = 20 males + 20 females, C_2_ = 19 males + 20 females and C_3_ = 20 males + 19 females) flies in total. The extraction of 352 samples was conducted at two different time periods each having approximately equal sample size. Each extraction time period consisted of three days, a day each for three replicates of MCF population. To be precise, replicates 1, 2 and 3 of MCF was extracted on the days 1, 2 and 3 respectively on two different extraction events. We used this information later in linear mixed effects analysis (see below) under the variable termed “time of extraction” that have six levels.

### Gas chromatography

Prior to GC-FID (gas chromatography fitted with flame ionization detector) analysis, the dried extract adhering to the walls of glass GC vials was re-dissolved in 20 μl *n-* Hexane and vortexed for 30 sec. We injected 2μl of this solution manually into splitless Shimadzu GC-2010 Plus instrument fitted with a Restek Rxi-5Sil MS (30m x 0.25mm x 0.25μm) capillary column. The 24.88 minutes’ temperature program began by holding at 57°C for 1.1 minutes, ramping at 100°C/min to 190°C with a hold time of 1.2 minutes. Further, with a ramp rate of 5°C/min, the temperature was increased to 270° and finally raised to 300°C at 120°C/min with a hold time of 5 minutes at 300°C (30). The carrier gas was Nitrogen with column flow rate fixed at 2ml per minute.

### Cuticular hydrocarbon identification

CHCs were identified with the help of GC-MS (gas chromatography mass spectrometer) using Agilent Technologies 7890B GC system fitted with Agilent J&W GC column (HP-5MS, 30m x 0.25mm x 0.25μm) and mass spectrometer (model no. 5977B GCMSD). The resultant chromatograms were analysed using Agilent MassHunter Qualitative Analysis Navigator B.08.00 by comparing mass spectra of each peak and SUPELCO *n*-alkane mixture standard (C8-C40) with characteristic retention indices. Further, the distinctive diagnostic ions were complemented with PUBCHEM compound summaries and probabilistic match of the mass spectras to the NIST Mass Spectral search program (version 2.3).

### Cuticular hydrocarbon quantification

All peaks that consistently appeared in the samples when compared with a blank chromatographic run (using only *n-*Hexane solvent) were considered relevant to this study. The area under the peak value was calculated by drawing a baseline to each relevant peak using the Shimadzu GCSolutions software. We manually checked the integrated peaks in all the sample chromatograms for any irregularities. The peak areas of the 352 samples was standardized by dividing area under each peak with that of internal standard (*n-*Pentadecane) from respective chromatograms. Each standardized value was then multiplied by 10000 ng/ml (concentration of *n-*Pentadecane) to derive relative concentration of each cuticular hydrocarbon in all samples for use in subsequent analysis.

### Linear mixed effects analysis

Linear mixed effects analysis involved two models that examined how different variables recorded during sampling from the MCF population affect abundance of each CHC peak in all samples. Selection regime (M, C and F), sex (male and female) and their interaction were considered explanatory variables with fixed effects while replicates within each selection regime (blocks 1, 2, and 3) and time of extraction were assigned as variables with random effects (30). We used “lme4” package in R (31) and modelled the fixed and random effects on the response variables according to the following formula:

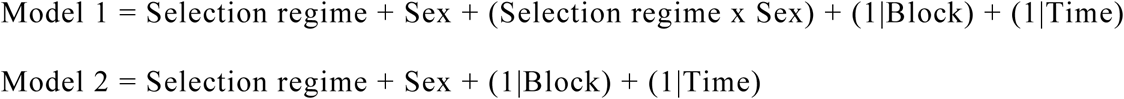

For the sake of simplicity, we present only Model 1 that includes all biologically relevant explanatory variables and their interaction. The simpler Model 2 (detailed analysis presented in S1 Supplementary Information) either had a higher Akike Information Criterion (AIC) or did not significantly differ from the Model 1 in the analysis of deviance, especially, where there was no significant effect of selection and sex interaction term on CHC variance. The distribution of each peak that are affected by the selection was further examined in details and their means were compared across sex and populations using Wilcoxon test to observe trends of differences in peak abundance across populations.

### Random Forest analysis

Closely related populations such as MCF usually show divergence in combinations of many CHCs rather than in a single peak. Therefore, we used Random Forest based multivariate analysis to study the divergence of CHCs. Multivariate analysis like principal component analysis and discriminant function analysis are generally used in studies that investigates group patterns within samples based on cuticular profiles (32). However, these statistical techniques are adversely impacted by small sample-size(n):variables(p), non-independence of CHCs in a sample (since many CHCs share common biochemical pathways (21)) and disproportionate effects of CHCs found in trace amounts (33). To overcome this drawbacks of traditional techniques, we used Random Forest analysis (R package “Party” (34)) that performs well under “small n large p” conditions, unaffected by highly correlated variables and is unbiased towards the nature of variables used (35, 36).

By comparing identities of the samples derived from the Random Forest analysis with their already known group identity, we calculate the percentage correct prediction of individual identities. Based on the probabilities of individual identities, proximity analysis (identical to similarity-dissimilarity matrix/ correlation matrix in traditional multivariate analysis) gives us a measure of samples’ group affinity. Multidimensional scaling (identical to Principal Component Analysis) of the proximity matrix is then illustrated in a scatterplot to show how experimental individuals group with respect to each other (37).

We performed Random Forest analysis using relative amount of all relevant CHC peaks. Seed (a random number) was set to 1220. We checked the stability of the classification scheme by increasing the number of subsampling events gradually from 5000 to 20000 and decided to set the number of trees (ntree) parameter in the analysis at 10000. The number of variables (mtry = 6) to be used in the analysis was fixed to rounded square root of the number of predictor variables (in the present study, the number of peaks analysed in chromatograms) in the data (Strobl et al. 2009b). Multidimensional scaling (MDS) of proximity matrix was used to produce two dimensional scatterplots showing clustering of individuals.

## Results

### Characterisation of MCF cuticular hydrocarbons

We found a total of 38 CHCs peaks that reliably and consistently appeared in MCF chromatograms (see Figure 1 and Table 1). Most of the CHCs that were identified using GCMS had hydrocarbons chain lengths between C_20_ and C_30_. Peak 17 turned out to be a combined peak of the hydrocarbon *n*-Tetracosane, and a non-hydrocarbon Adipate and was left out of further analyses. Only two peaks (Peak 3 and Peak 6) could not be precisely identified into known classes of CHCs. The appearance of different CHCs in *D. melanogaster* MCF flies appeared to follow a fixed pattern (or CHC fingerprint): methyl substituted straight chain *n*-alkanes appears first followed by differently positioned straight chain *n*-alkenes and finally, corresponding straight chain *n*-alkanes (Peak 10-14, Peak 18-23, Peak 26-30 and Peak 32-35).

**Fig. 1.**
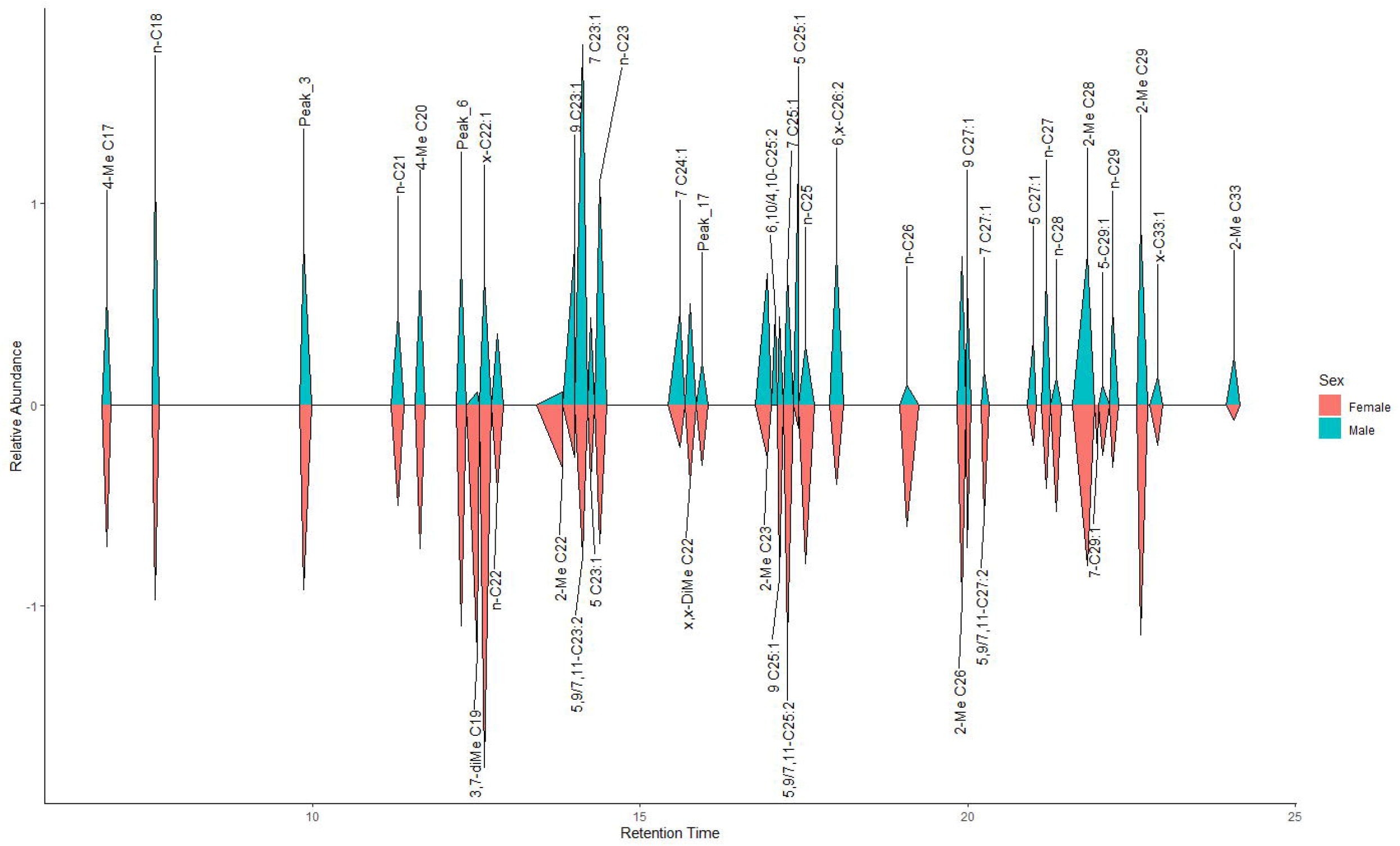
Representative chromatogram showing 38 cuticular hydrocarbon peaks of *Drosophila melanogaster* MCF population.

**Table 1:**
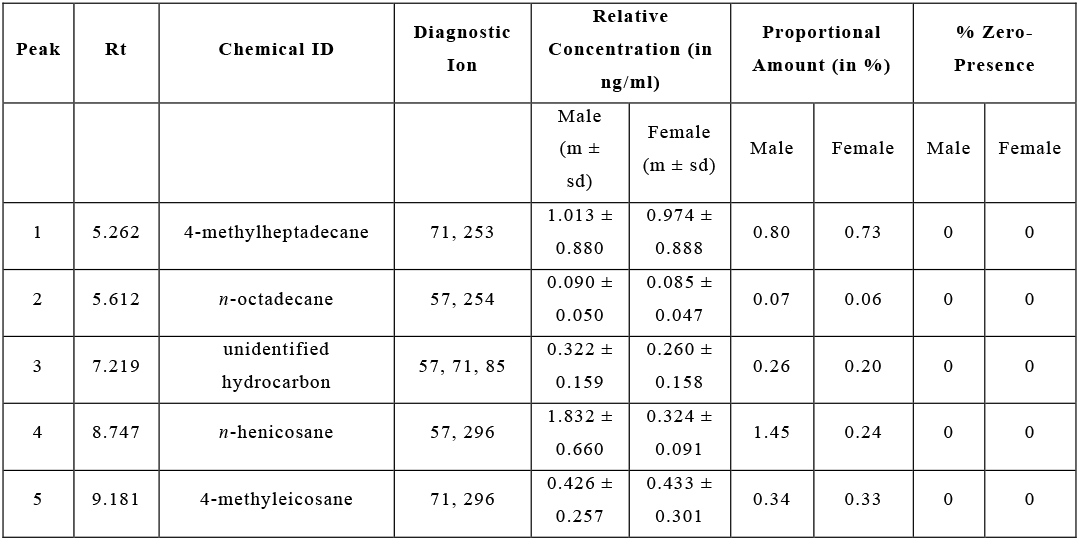
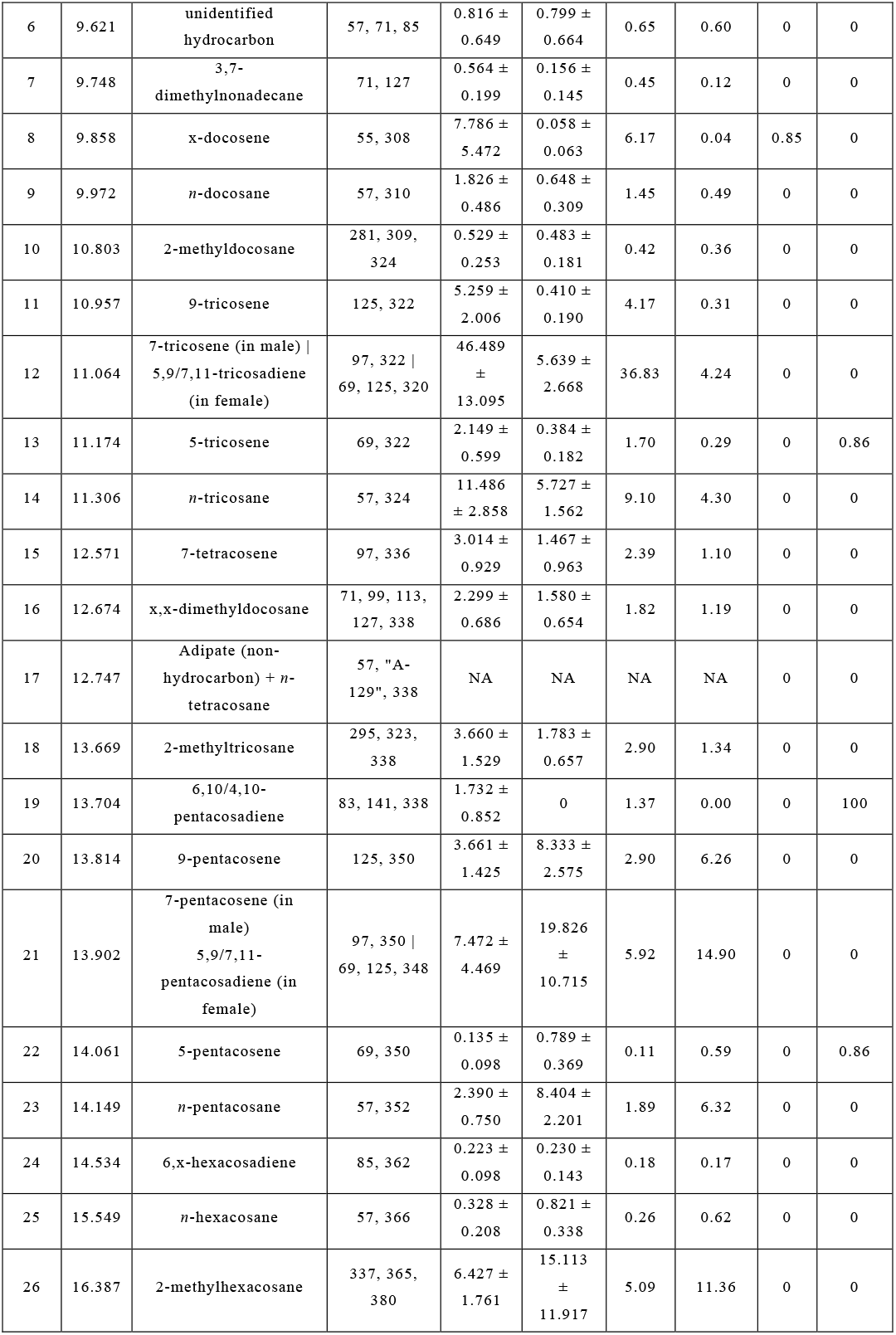
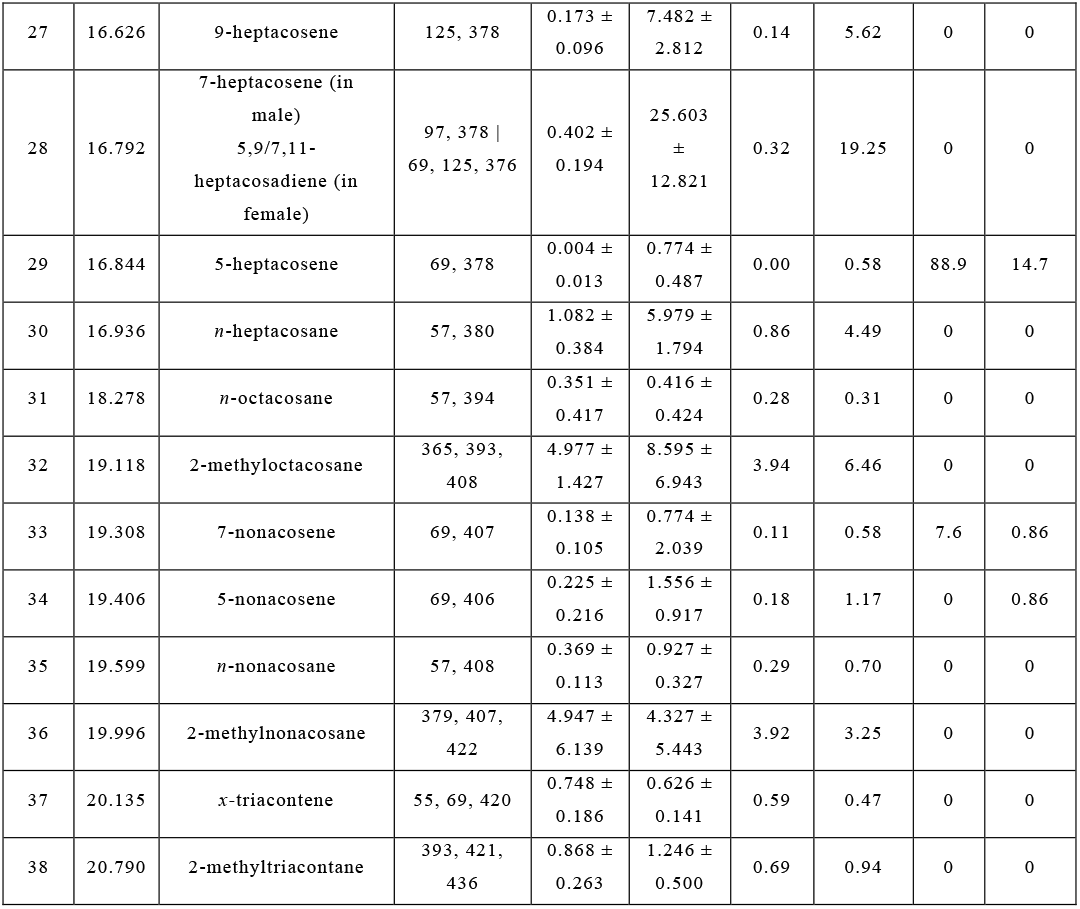
List of 38 cuticular hydrocarbon peaks along with their retention time (Rt), chemical identities (Chemical ID), diagnostic ions from mass spectrum found in *D. melanogaster* MCF population. Relative concentration (mean (m) and standard deviation (sd)), proportional amount of each cuticular hydrocarbon peak (in %) and % zero presence of 38 cuticular hydrocarbon peaks in both sexes of 352 *D. melanogaster* MCF population is also furnished here. % zero presence is the percentage of individuals that have absence (or zero presence) of a particular CHC.

The abundance of CHC peaks was measured using area under the peak relative to that of internal standard, *n-*Pentadecane that had a known concentration of 10 ng/ul in *n-* Hexane. Male and female chromatographic profiles significantly differed in abundance of most peaks. Peak 19 (6, 10/4, 10-pentacosadiene) apart from being the only alkadiene in male was totally absent in females indicating a likely sex specific role (alkadienes usually have pheromonal functions in female *D. melanogaster* (38)) in MCF population. Peak 29 (5-heptacosene) was generally absent from male chromatograms (% zero presence = 88.9) but were found in most females (% zero presence = 14.7). Peak 33 (7-nonacosene) that appeared consistently in female chromatograms were found to be missing in a few males (% zero presence = 7.6). Only eight CHCs occurred in similar quantities in both males and female.

*D. melanogaster* MCF males predominantly produce monoenes (e.g. Tricosene, Pentacosene, Heptacosene, Nonacosene) while the females predominantly produce dienes (e.g. Tricosadiene, Pentacosadiene, Heptacosadiene) (17). Most major and minor CHCs that are known to act as contact pheromones in *Drosophila melanogaster* during reproduction such as 7-tricosene (major), 5-tricosene (minor), 9-pentacosene (minor), 7-pentacosene (minor) and 5,9/7,11-heptacosadiene (major) were found in MCF population (17). However, a major female contact pheromone, 7,11-nonacosadiene was absent from the MCF chromatograms. It is interesting to note that 7-monoalkenes (Peaks 12, 21 and 28) occur in males only but corresponding peaks in females are a mixture of 5,9- and 7,11-alkadienes. The 5,9-alkadiene isomer is predominant over 7,11-alkadiene isomer in MCF females. 5,9-alkadiene was found to be present in 17 *D. melanogaster* Genetic Reference Panel recently due to presence of *Desat2* allele that was initially thought to occur only in African and Caribbean *D. melanogaster* (21).

### Effects of selection regime, sex, replicate and the time of extraction on the MCF cuticular hydrocarbons

We considered 37 MCF CHCs (Peak 17 was not considered since it is a combined peak of Adipate and *n*-tetracosane) for the linear mixed effects analysis (see S1 Supplementary Information). Our model considered time of extraction and replicates of MCF population as variables with random effects and selection line, sex and their interaction as variables with fixed effect on the CHCs. Time of extraction (see Methods section under subsection Cuticular hydrocarbon extraction for details) refers to six different time points when CHC extractions of *D. melanogaster* samples were done. Among variables with random effect, we compared the variance in replicates and time of extraction with that of the individual (residual) variance (see S1 Supplementary Information). We found no peaks where MCF replicates had more variance than the residuals. In fact, most of the CHC peaks had negligible variance for blocks indicating no effect of replicate populations on peak variability. However, six peaks (Peaks 1, 3, 5, 6, 36 and 37) were affected by time of extractions (variance due to time of extraction was more than the residual). These peaks were not considered for Random Forest analysis. The sex of the sampled *D. melanogaster* MCF individuals is the most important variable affecting 30 CHC peaks (see Table 2) as seen previously in Table 1. The variables with fixed effects such as selection regime affected 19 CHCs while its interaction with sex of the MCF individuals affected 10 CHCs (see Table 2). Only three CHCs (Peak 24, 31 and 36) are not affected by any of the explanatory variables.

**Table 2:** Influence of variables with fixed effects (selection, sex and their interactions) and random effects (block and time of extraction) on 37 cuticular hydrocarbons of *D. melanogaster* MCF population assessed by linear mixed effect model analysis. Those fixed effect variables with statistically significant effect have been represented by a p-value based on *Type II Wald chi-square* test. Those variables with no significant effect on cuticular peak abundance has been denoted NS. Effect of replicates of selection lines (Block) and time of extraction (Time) was assessed by comparing their variance with that of the residual (see S1 Supplementary Information). † represents known sex pheromones in *D. melanogaster*.

| Peak | Selection | Sex | Selection x Sex | Time | Block |
| --- | --- | --- | --- | --- | --- |
| 1 | NS | NS | p = 0.005413 ** | Effect present | NS |
| 2 | p = 0.02986 * | NS | NS | NS | NS |
| 3 | p = 0.01683 * | p < 2e-16 *** | NS | Effect present | NS |
| 4 | NS | p < 2e-16 *** | NS | NS | NS |
| 5 | P = 0.02936 * | NS | NS | Effect present | NS |
| 6 | NS | NS | p = 0.01266 * | Effect present | NS |
| 7 | NS | p < 2e-16 *** | NS | NS | NS |
| 8 | NS | p < 2e-16 *** | NS | NS | NS |
| 9 | p = 0.03019 * | p < 2e-16 *** | NS | NS | NS |
| 10 | NS | p = 0.02483 * | NS | NS | NS |
| 11 | NS | p < 2e-16 *** | NS | NS | NS |
| 12† | p = 0.04202 * | p < 2e-16 *** | NS | NS | NS |
| 13† | NS | p < 2e-16 *** | NS | NS | NS |
| 14 | p = 0.004357 ** | p < 2e-16 *** | NS | NS | NS |
| 15 | p = 7.744e-05 *** | p < 2e-16 *** | NS | NS | NS |
| 16 | NS | p < 2e-16 *** | NS | NS | NS |
| 18 | NS | p < 2e-16 *** | NS | NS | NS |
| 19 | p = 0.016364 * | p < 2e-16 *** | p = 0.009102 ** | NS | NS |
| 20† | p = 0.005125 ** | p < 2e-16 *** | NS | NS | NS |
| 21† | p = 0.01224 * | p < 2e-16 *** | NS | NS | NS |
| 22 | p = 0.001317 ** | p < 2e-16 *** | p = 0.009922 ** | NS | NS |
| 23 | p = 0.0001536 *** | p < 2e-16 *** | p = 0.0285458 * | NS | NS |
| 24 | NS | NS | NS | NS | NS |
| 25 | p = 0.002671 ** | p < 2e-16 *** | NS | NS | NS |
| 26 | NS | p < 2e-16 *** | NS | NS | NS |
| 27 | p = 0.001985 ** | p < 2e-16 *** | p = 0.001615 ** | NS | NS |
| 28† | p = 0.0009216 *** | p < 2e-16 *** | p = 0.0007424 *** | NS | NS |
| 29 | p = 0.04211 * | p < 2e-16 *** | p = 0.03855 * | NS | NS |
| 30 | p = 0.003904 ** | p < 2e-16 *** | p = 0.008833 ** | NS | NS |
| 31 | NS | NS | NS | NS | NS |
| 32 | NS | p = 7.81e-13 *** | NS | NS | NS |
| 33 | NS | p = 2.547e-05 *** | NS | NS | NS |
| 34 | p = 0.04416 * | p < 2e-16 *** | NS | NS | NS |
| 35 | NS | p < 2e-16 *** | p = 0.01684 * | NS | NS |
| 36 | NS | NS | NS | Effect present | NS |
| 37 | p = 0.03032 * | p < 2e-16 *** | NS | Effect present | NS |
| 38 | NS | p < 2e-16 *** | NS | NS | NS |

The 19 peaks affected by differential operational sex ratio i.e. selection regime as shown in linear mixed effects analysis are an obvious point of interest in our study. However, 3 (Peak 3, 5 and 37) of those 19 peaks were affected by differences in time of extractions arising out of logistical reasons. Rest of the 16 peaks includes four known sex pheromones (Peak 12, 20, 21 and 28). To examine the trend of these 16 CHC peak distributions that were not affected by the time of extraction but affected by imposed selection, we plotted grouped Box and Whisker graphs (see S2 Supplementary Information). The differences in mean CHC amounts between sexes mirrored linear mixed effects results (see Table 3). We found that, unlike the rest of 15 CHC peaks, sexual dimorphism in Peak 2 did not exist in F and C populations. Peak 2 was sexually dimorphic in M population only. By comparing means of peak distribution of M and F populations in each sex with respect to C population, we found that M and F males and females are not different from C in most of the cases. Both M and F males was different from C males for Peak 19 while only M males was different from C males for Peak 9. In case of females, we found differences between M population and C population in four peaks (Peak 20, 23, 27 and 30) but there was no difference between F population and C in any peak distribution. Qualitatively, in comparison to C population, it appears that the amount of some CHCs have changed only in M population than F population. Comparison of means of CHC peak distribution between M and F population indicates that only Peak 15 is significantly different in males while most of the differences between M and F populations are due to differences in 11 CHC peaks in females (Peak 2, 20, 21, 22, 23, 25, 27, 28, 29, 30 and 34). The analysis indicates that male M and F CHC profiles have not evolved to be different from that of C males and the differences between M and F males themselves are based on only one peak. Any divergence in M and F CHC profiles may have been because of the M females that have significantly diverged from C females (see S2 Supplementary Information). Although F females have not diverged from C females in most peaks, M females were found to be significantly different from F females in majority of the CHC peaks. It is interesting to note that 7 CHCs found on both M and F females that were not significantly different from C turned out to be significantly different between M and F populations themselves.

**Table 3:** Sexual dimorphism in sixteen cuticular hydrocarbons (that was affected by differential operational sex ratios but not time of extraction as shown in linear mixed effects analysis) found on *D. melanogaster* MCF population. M represents male biased population, F represents female biased population and C represents control population of unit sex ratio. Sex-wise comparison using Wilcoxon test of the means of peak distributions between M and F populations when compared to C as well as between themselves are presented. * represents statistically significant difference (* means p ≤ 0.05, ** means p ≤ 0.01 and *** means p ≤ 0.001) and † represents known sex pheromones in *D. melanogaster* (see S2 Supplementary Information). Stars coloured red indicate males have higher amount of the particular CHC whereas golden stars indicate females have higher amount of the particular CHC than males.

| Peaks | Sexual Dimorphism |  |  | Male |  |  | Female |  |  |
| --- | --- | --- | --- | --- | --- | --- | --- | --- | --- |
|  | M | C | F | M vs C | F vs C | M vs F | M vs C | F vs C | M vs F |
| 2 | * | NS | NS | NS | NS | NS | NS | NS | **, F>M |
| 9 | *** | *** | *** | *, C>M | NS | NS | NS | NS | NS |
| 12† | *** | *** | *** | NS | NS | NS | NS | NS | NS |
| 14 | *** | *** | *** | NS | NS | NS | NS | NS | NS |
| 15 | *** | *** | *** | NS | NS | *, F>M | NS | NS | NS |
| 19 | *** | *** | *** | *, C>M | **, C>F | NS | Peak absent in females |  |  |
| 20† | *** | *** | *** | NS | NS | NS | *, C>M | NS | *, F>M |
| 21† | *** | *** | *** | NS | NS | NS | NS | NS | *, F>M |
| 22 | *** | *** | *** | NS | NS | NS | NS | NS | **, F>M |
| 23 | *** | *** | *** | NS | NS | NS | *, C>M | NS | **, F>M |
| 25 | *** | *** | *** | NS | NS | NS | NS | NS | *, F>M |
| 27 | *** | *** | *** | NS | NS | NS | *, C>M | NS | ***, F>M |
| 28† | *** | *** | *** | NS | NS | NS | NS | NS | **, F>M |
| 29 | *** | *** | *** | NS | NS | NS | NS | NS | *, F>M |
| 30 | *** | *** | *** | NS | NS | NS | *, C>M | NS | ***, F>M |
| 34 | *** | *** | *** | NS | NS | NS | NS | NS | **, F>M |

### Divergence patterns of MCF cuticular hydrocarbons

We used 31 out of 37 MCF CHCs, whose variance were not affected by time of extraction as shown in linear mixed effects analysis, for multivariate analysis using Random Forest. Multidimensional scaling of proximity matrix showed that sexes formed distinct groups in *D. melanogaster* MCF population with 100 % correct sex identification in random forest analysis. In C population, Random Forest analysis produced 94.1% correct prediction of individuals belonging to 3 different replicates. It is important to note that the triplicates of C are maintained at unit sex ratio and are derived from LHst while M and F populations are derived from the C replicates (see Methods section for details). Therefore, the divergence pattern of C in Fig. 2 can be considered most close to ancestral pattern and is subsequently used as reference pattern to M and F populations. The CHC divergence pattern in Fig. 3 suggests that the M females are more dispersed than the M males that are tightly grouped (91 % correct identification of M individuals from 3 replicate populations). In comparison to C population, M males appear to have evolved CHC profiles which are more similar to each other while M females have evolved a more variable CHC profile than C females. In F population (Fig. 4), we see an opposite trend where the F males are dispersed more than the F females that form tight group (87 % correct identification of F individuals belonging to 3 replicate populations). Comparison with C females suggests that F females have evolved CHC profiles that are similar to each other while F males have increased the variability of their CHC profile.

**Fig. 2.**
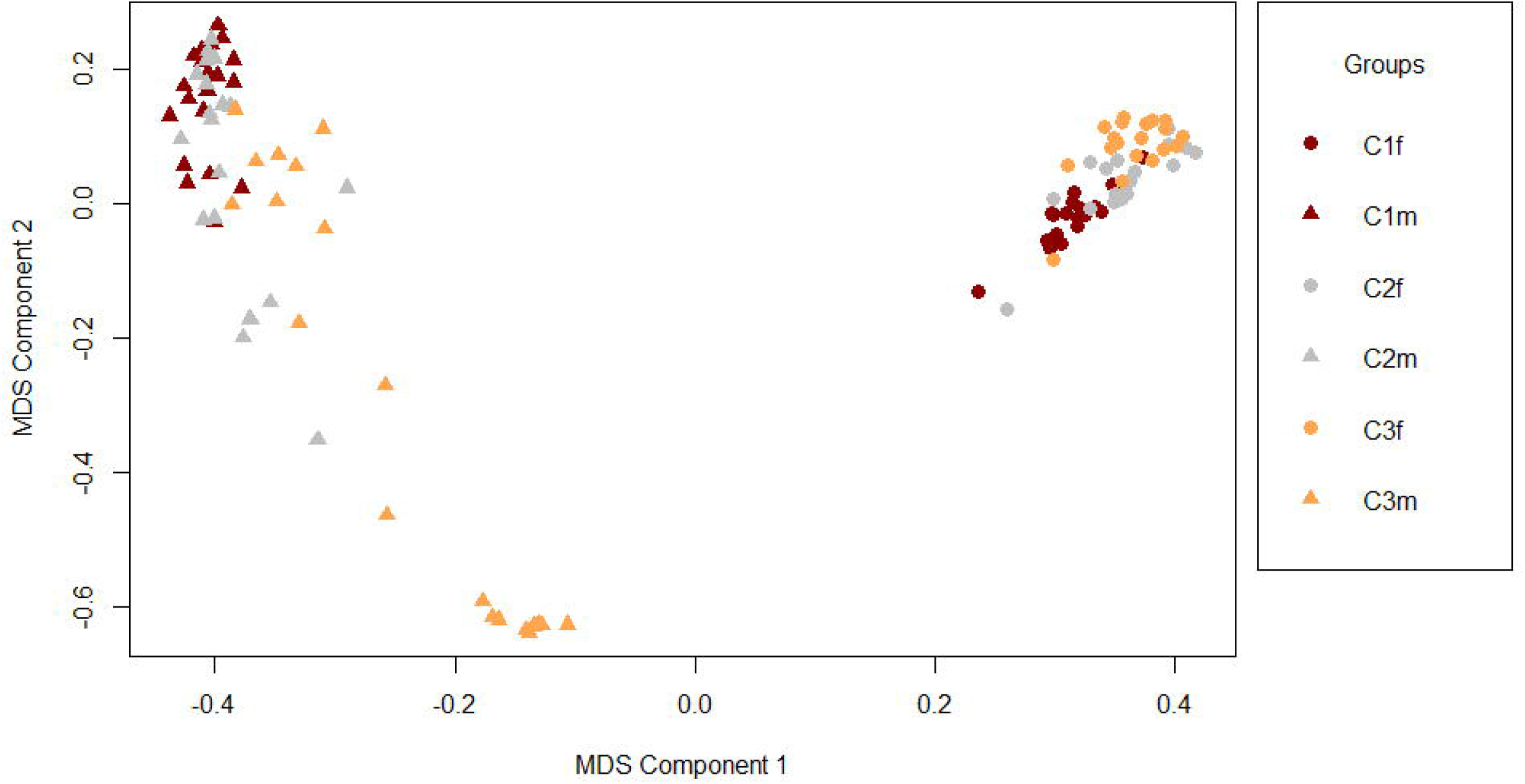
Multi-dimensional scaling (MDS) of the proximity matrix of C individuals (C_1/2/3_ are C blocks, m= males and f= females)

**Fig. 3.**
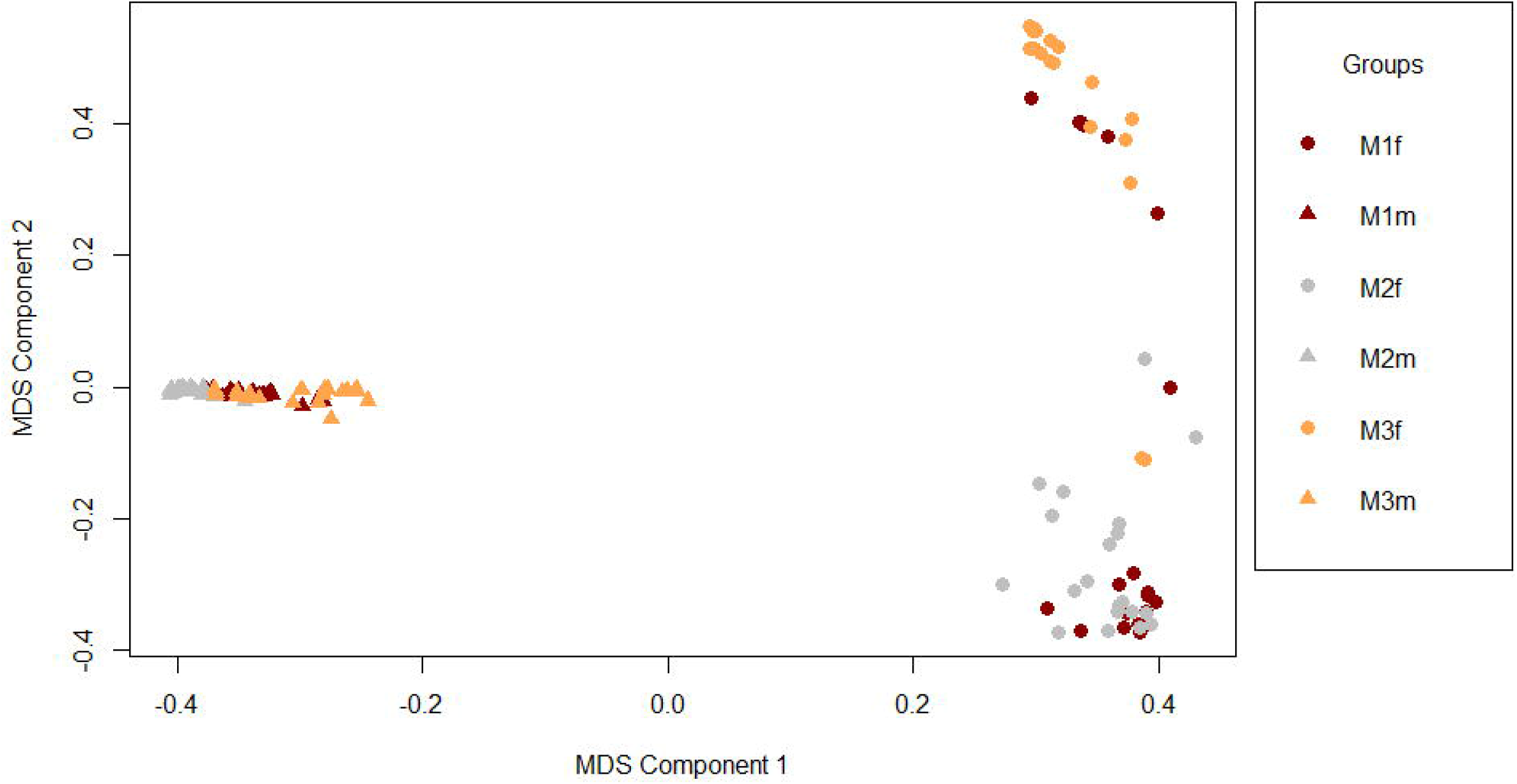
Multi-dimensional scaling (MDS) of the proximity matrix of M individuals (M_1/2/3_ are M blocks, m= males and f= females)

**Fig. 4.**
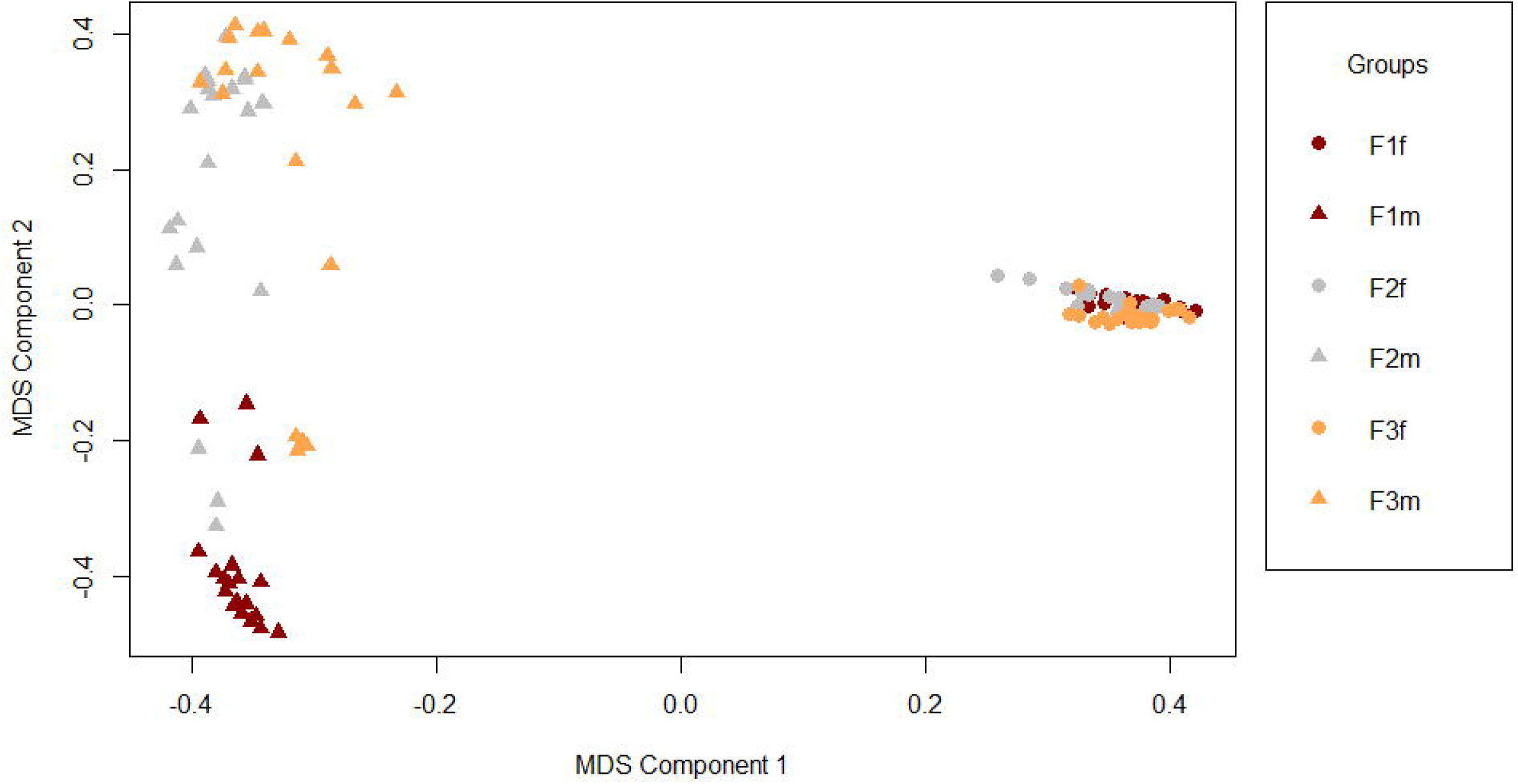
Multi-dimensional scaling (MDS) of the proximity matrix of F individuals (F_1/2/3_ are F blocks, m= males and f= females)

When we compare male CHC profile of M and F replicates, Random Forest analysis predicted 86.4% correct identification of the M_1,2,3_ and F_1,2,3_ males (see Fig. 5). In Fig. 5, males of M replicates formed separate groups with few overlaps among themselves. However, F replicates did not group separately from the M replicates as predicted but overlapped with the corresponding M replicates. In Random Forest analysis involving females of M and F replicates, we could identify M_1,2,3_ and F_1,2,3_ females with 85.3% correct prediction (see Fig. 6). Fig. 6 revealed that M females from different replicates formed separate groups with few overlaps. We again find a considerable overlap of F replicates on corresponding M replicates except for females from second F replicate.

**Fig. 5.**
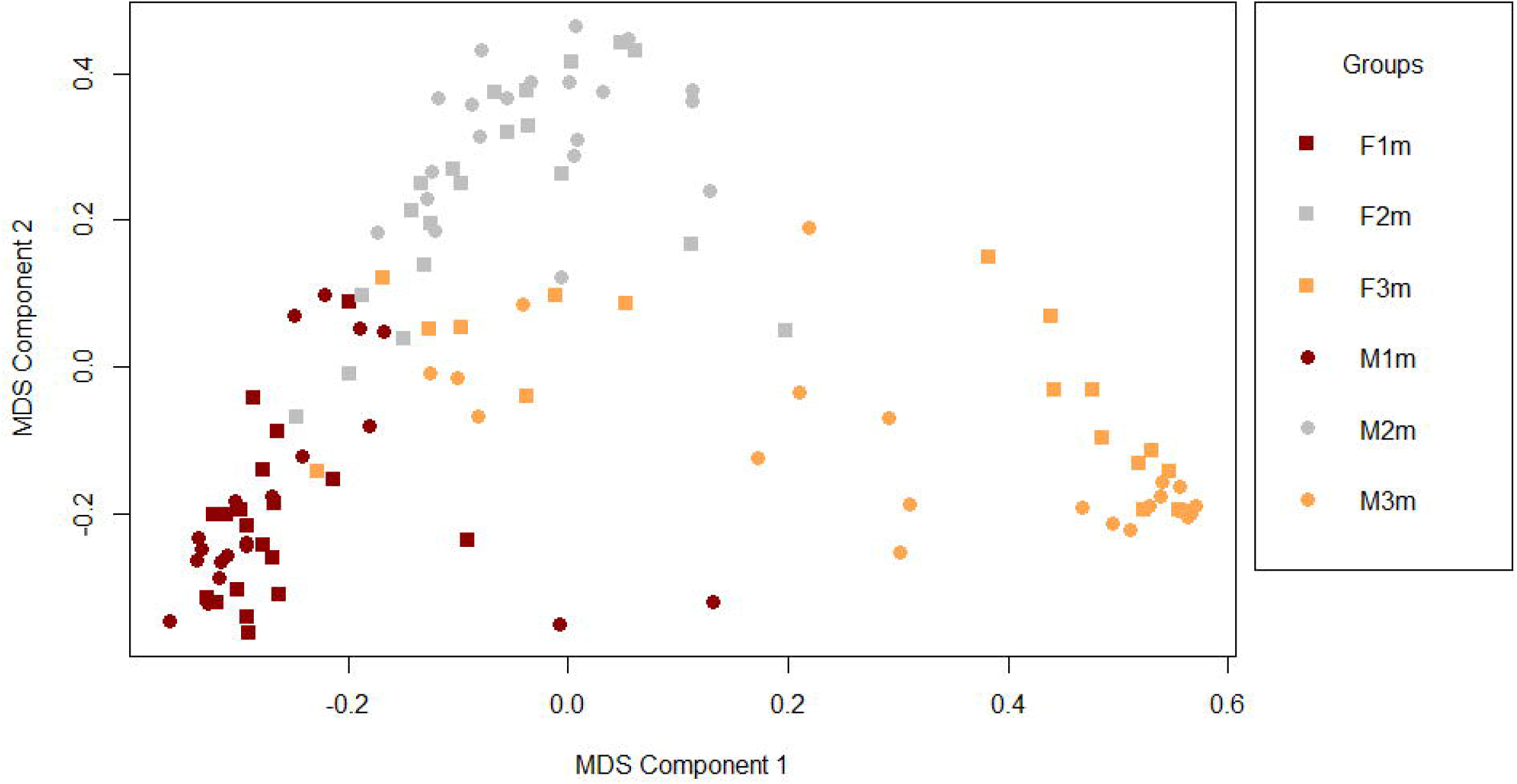
MDS of proximity matrix derived from Random Forest analysis of males from three M and F replicates

**Fig. 6.**
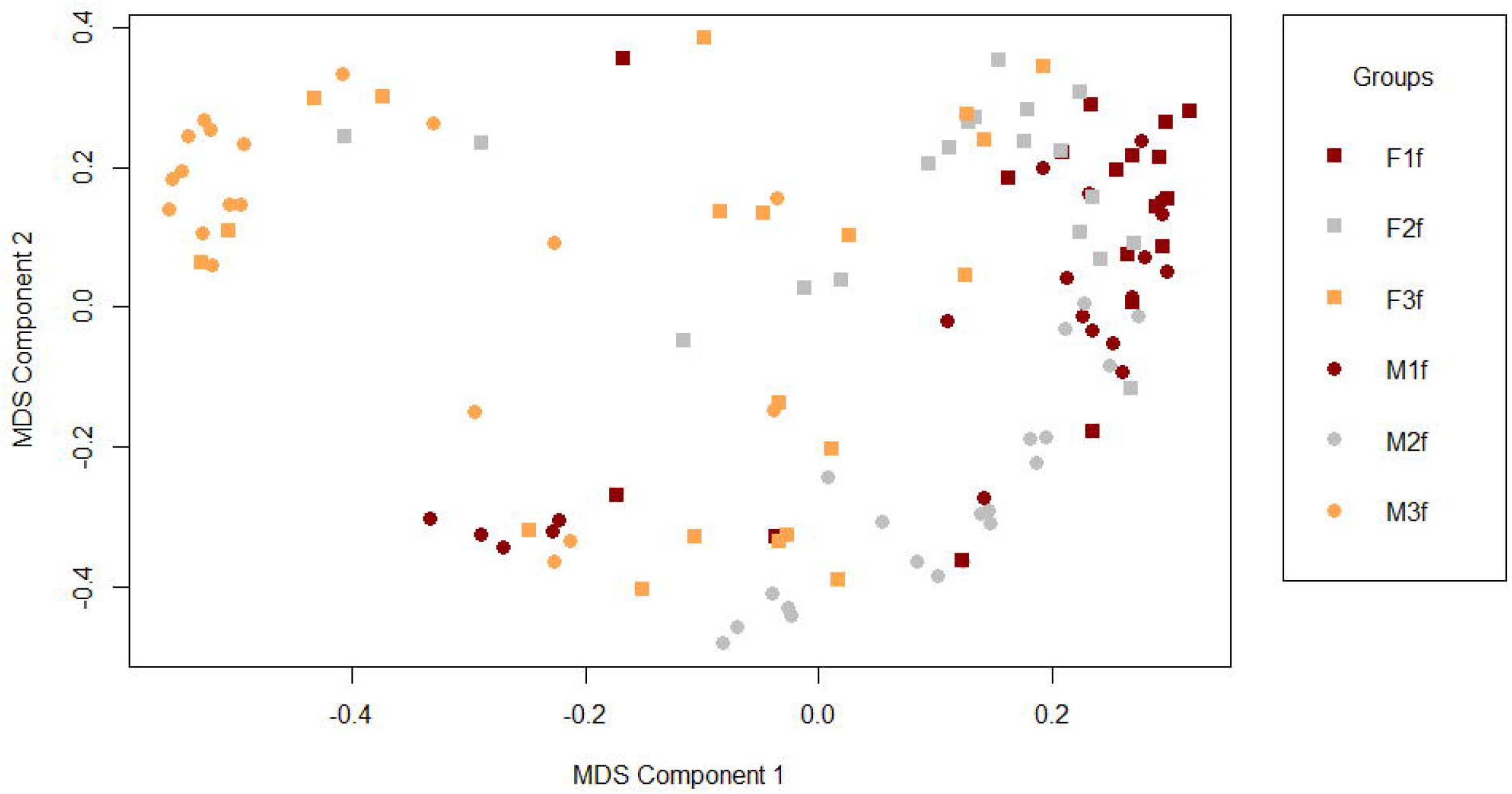
MDS of proximity matrix derived from Random Forest analysis of females from three M and F replicates

We also performed Random Forest analysis with the 16 CHC peaks that were significantly affected by differential operational sex ratios but not time of extraction (see linear mixed effects analysis, Table 2). The MDS plots based on these peaks are presented in S3 Supplementary Information. While there are minor differences between MDS based on 31 and 16 peaks, the overall pattern of scatter (i.e. tightly packed M males and F females in M and F populations respectively; divergence of M and F males and females) still held strong.

## Discussion

Reproductive isolation evolved as a by-product of SAC within each M replicates (8). The allopatric M replicates of *D. melanogaster* MCF population diverged from each other in terms of mate preference, copulation duration and sperm defense ability over 105 generations. However, mating latency that is considered to be a measure of female reluctance to mate did not differ among M replicates. Therefore, it was concluded that the divergent mate preference in the high sexual conflict regime of the MCF population may most likely be due to underlying female choice (8). Since mate preference did not diverge in F replicates under low sexual conflict, we predicted that male and female CHC profiles of the three replicates within the M regime should be more divergent from each other compared to the CHC profiles of the three replicates within the F regime. Here we assume that the putative chemical cues such as the CHCs form the basis of divergent mate choice in M and have also diverged as a by-product of SAC.

Abundances of different CHC peaks that are affected by altered operational sex ratio indicate that the predicted divergence of M replicates may not hold true at least for males (see Table 3 and S2 Supplementary Information). The results indicate that most male CHCs of M population have diverged little from either C and F males. Among the CHCs found in *D. melanogaster* MCF population, male CHCs (such as 7-tricosene, 9-pentacosene and 7-pentacosene) that have known pheromonal roles show no difference among selection lines. On the other hand, M females evolved CHC differences from C females in four peaks but F females did not differ in CHCs from C females at all. However, M females showed differences in peak abundance with F females in many CHCs including the known contact pheromone Heptacosadiene, in spite of F females not significantly differing from C females. Any major differences between M and F populations, therefore, are due to CHC abundance on female cuticle. The distribution of CHC abundance in both males and females shows quite a consistent trend whereby M population has decreased peak abundance while F population has increased peak abundance (see S2 Supplementary Information). The CHC peaks of M males also have a near consistent tighter distribution apart from being less abundant than F males. It appears as though M males are compelled to display a typical CHC profile, possibly due to the highly competitive scenario in accessing mates, indicating an intense sexual selection. Less abundant CHCs in M females, however, could be because of various reasons including (a) to reduce attractiveness (viz. decreasing amount of peak abundances) to harassing males, (b) increased energy/resource investment in resistance to male induced harm, etc. However, at this point, we have no evidence supporting any of the possibilities.

Random Forest analysis (see Fig. 5) shows that the three replicates of M males form 3 distinct groups but contrary to our prediction, F males do not form a single group. Instead, the three F replicates overlap with the three corresponding M replicates. A similar divergence pattern is also reflected in two of the three female replicates of M and F (see Fig. 6). Such CHC divergence pattern indicates that SAC did not result in CHC differentiation in M population in a similar way it had led to divergence in mating behaviour in M population. The within-replicate assortative mating in the high sexual conflict regime may, therefore, be mediated by mate choice on mate attracting cues other than CHCs. It is difficult to ascertain how M and F replicates could have an overlapping CHC divergence pattern though. Since M and F populations do not differ from C population in the abundance and distribution of many CHCs, it is possible that the divergence pattern of M and F reflect that of the C population whose sexes show quite a differentiated profile of CHCs among its three replicates (see S4 Supplementary Information). The pattern of CHC divergence in MCF sexes could otherwise be linked to ancestry. However, since MCF population was established using a large population size (N_e_ (LH_st_) ~ 2250, N_e_ (MCF) ~ 450; (28)), a founder effect during initial establishment is unlikely to have affected the MCF population. This is also confirmed in the linear mixed modelling analysis which shows effect of replicates on the CHC profile is negligible (see Table 2). In conclusion, reproductive isolation (in M regime but not in F regime (8)) as a by-product of SAC cannot be explained by the CHC divergence among males and females of *Drosophila melanogaster* M and F regimes. However, CHC divergence in M and F populations at regime level indicate other possible evolutionary mechanisms at play that are discussed below.

In this study, sexual selection appears to have particularly strong effect on the CHC profiles at the population level (see Fig. 2, Fig. 3 and Fig. 4). It is important to note, here, that CHC profile refers to the combination of different CHCs (CHC bouquet) that acts as a composite pheromonal signal rather than individual CHCs. If we compare the CHC profiles of C and M populations, we see that M females have diverged with respect to C females while M males have converged with respect to C males considerably. Increased sexual selection in M population appears to have resulted in CHC profiles of M males being pushed toward a common optimum point such that M males present a more preferable CHC profile that is under stabilising selection to M females. Similar CHC profiles of the M males may also be explained by suppression of sexually selected CHCs such that variability of male CHCs is reduced over time. This may be true since *D. serrata* males have been found to lower their CHC expression when female:male ratio is decreased (39). We cannot, also, rule out the possibility of the male CHC profile of M population coevolving with other sexually selected trait(s). On the other hand, divergence of CHC profiles of M females could be a result of lack of selection pressure due to abundant mating options or as predicted in populations following Buridan’s ass regime (see below).

The F males and females show a diametrically opposite pattern to that of the M regime. Under low sexual selection, the selection pressure is possibly weaker and hence, absence of directional selective pressures allowed diversification of CHC profiles in F males compared to the C males. In F regime where female:male is high, males may evolve to display their most attractive CHCs to potentially improve mating success (as seen in *D. serrata* (39)) leading to varied CHC profiles in the process. The F females appear to be more tightly grouped in terms of their CHC profile than C females (and also M females). This indicates that F females are probably under selection pressure to maintain similarity of their CHC profiles. The reason for this is not obvious and may be a result of correlated evolution in some other trait affected by sexual selection. Although not experimentally tested, *D. melanogaster* females do possess specific mate attracting contact pheromones that suggest some degree of female control over mating outcome and fitness derived from such mating (17). Another possibility, therefore, exist where F females may also be affected by increased competition due to altered sex ratio where they have to maintain identical profile to be acceptable to males that are fewer in number. Overall, the CHC divergence pattern of M and F regime appears to be strongly influenced by sexual selection although other possible sexual conflict resolution mechanism such as Buridan’s ass paradigm (see below) cannot be ruled out.

Apart from speciation, a population under sexual conflict may follow other evolutionary stable strategy to resolve the conflict between sexes (6). For example, in a large population and/or population with significant rates of mutations, there might be diversification of female traits with respect to male traits (Buridan’s ass regime) such that males, in an attempt to adapt to the variability in female traits, are caught in limbo unable to increase their own fitness in the population as a whole (1). This is relevant to the results from the current study where multivariate analysis show M females of each replicate are more dispersed and M males form a tight group with respect to their CHC profile (see Fig. 3). It is known that the male mating behavior of *Drosophila melanogaster* is partially mediated by female CHCs (17). The M population also meet the criterion for Buridan’s ass regime since the population size is large (N_e_ ~ 450) and the block-wise differences among M replicates (see Fig. 5) suggest that possibility of random mutation accumulation over several generations in allopatry. It is, therefore, possible that sexual conflict on CHC profiles in M regime has followed Buridan’s ass phenomenon where M males have been caught in an uncertain intermediate stage failing to pursue either ends of CHC divergence in M females. The fact that this pattern is repeated in case of females of F regime is a revelation since effect of sexual conflict has been traditionally focused on males rather than females (17).

## Conclusion

Experimental studies and comparative analysis of taxonomic groups suggest that speciation is possible under sexual conflict. The mathematical models support this possibility but also suggest that dynamics of evolution under sexual conflict is quite varied and stochastic to a point that evolution of reproductive isolation in allopatry is only one of the several other outcomes. This study is an example of different reproductive traits being affected differently by sexual conflict such that isolation based on mate preference, copulation duration and sperm defense ability is not reflected in the mate attraction cues such as CHCs. Rather sexual conflict, if any, is probably reconciled in form of Buridan’s ass regime. In fact, *D. melanogaster* individuals from high and low sexual conflict regime appear to be significantly affected by sexual selection on CHC profiles. We conclude that since CHCs form a basis for mate attraction, M males under high sexual selection may be likely to keep their CHC profiles similar to be acceptable by the limited number of females in the male biased regime. The same may also be true for the females of female biased regime or their CHC divergence pattern may be by-product of sexual selection pressure on any other hitherto unknown trait.

## Acknowledgements

We acknowledge Dr. Anoop Ambili, laboratory head of the PRISM group in the Department of Earth and Environmental Sciences, Indian Institute of Science Education and Research Mohali, India for the access to the GC-MS facility. We also thank all the members of Evolutionary Biology Laboratory at Indian Institute of Science Education and Research, Mohali for their numerous small but essential assistance during this study.

