## Supplementary material for "Cuticular hydrocarbon divergence in *Drosophila melanogaster* populations evolving under differential operational sex ratios": SI1

Journal Name: Evolutionary Biology

Dutta^1*^, Tejinder Singh Chechi^1^, Ankit Yadav^2^, Nagaraj Guru Prasad^1^

^1^Department of Biological Sciences, Indian Institute of Science Education and Research, Mohali, India

^2^Department of Earth and Environmental Sciences, Indian Institute of Science Education and Research, Mohali, India

Results from linear mixed effect model analysis of *D. melanogaster* MCF cuticular hydrocarbons

| **Peak ID** | **Model 1 AIC Value** | **Model 2 AIC Value** | **Difference between Model 1 and Model 2** | **Chosen Model** | **Variance of Time in Model 1** | **Variance of Blocks in Model 1** | **Residual Variance in Model 1** |
| --- | --- | --- | --- | --- | --- | --- | --- |
| Peak 1 | 221.3 | 227.6 | p = 0.006 ** | Model 1 | 6.812e-01 | 2.200e-10 | 9.409e-02 |
| Peak 2 | -1251.4 | -1249.6 | p = 0.057 | Model 1 | 8.043e-04 | 4.150e-12 | 1.498e-03 |
| Peak 3 | -826.3 | -827.6 | p = 0.251 | Model 1 | 2.007e-02 | 2.452e-11 | 4.843e-03 |
| Peak 4 | 381.7 | 379.3 | p = 0.459 | Model 1 | 6.405e-02 | 6.913e-08 | 1.557e-01 |
| Peak 5 | -155.4 | -158.4 | p = 0.581 | Model 1 | 0.044 | 0.00 | 0.033 |
| Peak 6 | 14.5 | 19.2 | P = 0.013 | Model 1 | 3.754e-01 | 6.685e-07 | 5.230e-02 |
| Peak 7 | -337.3 | -340.9 | p = 0.794 | Model 1 | 0.006 | 0.004 | 0.020 |
| Peak 8 | 1949.3 | 1947.1 | p = 0.402 | Model 1 | 1.13 | 0.00 | 13.7 |
| Peak 9 | 208.1 | 207.5 | p = 0.180 | Model 1 | 0.068 | 0.00 | 0.094 |
| Peak 10 | -117.9 | -121.1 | p = 0.678 | Model 1 | 0.007 | 0.003 | 0.038 |
| Peak 11 | 1180.8 | 1180.5 | p = 0.162 | Model 1 | 0.348 | 0.128 | 1.52 |
| Peak 12 | 2502.7 | 2502.6 | p = 0.141 | Model 1 | 22.7 | 0.00 | 64.6 |
| Peak 13 | 349.6 | 348.1 | p = 0.285 | Model 1 | 5.011e-02 | 4.173e-11 | 1.425e-01 |
| Peak 14 | 1461.2 | 1458.4 | p = 0.568 | Model 1 | 1.84 | 0.00 | 3.33 |
| Peak 15 | 758.7 | 755.2 | p = 0.793 | Model 1 | 4.197e-01 | 8.259e-08 | 4.484e-01 |
| Peak 16 | 593.0 | 589.5 | p = 0.784 | Model 1 | 0.105 | 0.057 | 0.283 |
| Peak 18 | 1069.4 | 1066.0 | p = 0.718 | Model 1 | 0.263 | 0.00 | 1.11 |
| Peak 19 | 602.7 | 608 | p = 0.010 * | Model 1 | 0.037 | 0.015 | 0.296 |
| Peak 20 | 1469.7 | 1470.3 | p = 0.102 | Model 1 | 0.668 | 0.00 | 3.47 |
| Peak 21 | 2466.1 | 2464.2 | p = 0.358 | Model 1 | 5.43 | 0.00 | 59.5 |
| Peak 22 | 41.0 | 46.1 | p = 0.011 * | Model 1 | 0.008 | 0.00 | 0.060 |
| Peak 23 | 1247.5 | 1250.6 | p = 0.030 * | Model 1 | 7.065e-01 | 2.076e-09 | 1.824e+00 |
| Peak 24 | -552.6 | -556.0 | p = 0.739 | Model 1 | 0.003 | 0.001 | 0.011 |
| Peak 25 | -32.4 | -31.6 | p = 0.090 | Model 1 | 2.859e-02 | 1.356e-09 | 4.772e-02 |
| Peak 26 | 2511.8 | 2508.9 | p = 0.601 | Model 1 | 2.09 | 1.03 | 68.4 |
| Peak 27 | 1467.6 | 1476.3 | p = 0.002 ** | Model 1 | 0.129 | 0.00 | 3.53 |
| Peak 28 | 2521.3 | 2531.4 | p = 0.0009 *** | Model 1 | 3.28 | 2.68 | 69.7 |
| Peak 29 | 246.1 | 248.6 | p = 0.040 * | Model 1 | 0.003 | 0.00 | 0.110 |
| Peak 30 | 1112.4 | 1117.7 | p = 0.009 ** | Model 1 | 0.339 | 0.00 | 1.25 |
| Peak 31 | 352.7 | 350.0 | p = 0.542 | Model 1 | 0.029 | 0.001 | 0.145 |
| Peak 32 | 2122.2 | 2119.4 | p = 0.556 | Model 1 | 2.08 | 0.378 | 22.3 |
| Peak 33 | 1263.4 | 1263.6 | p = 0.120 | Model 1 | 0.000045 | 0.00 | 2.01 |
| Peak 34 | 690.4 | 690.9 | p = 0.105 | Model 1 | 0.044 | 0.002 | 0.382 |
| Peak 35 | -42.5 | -38.4 | p = 0.018 * | Model 1 | 0.010 | 0.00 | 0.047 |
| Peak 36 | 1783.6 | 1780.2 | p = 0.725 | Model 1 | 25.2 | 0.00 | 8.08 |
| Peak 37 | -509.3 | -511.8 | p = 0.478 | Model 1 | 1.444e-02 | 3.136e-08 | 1.217e-02 |
| Peak 38 | 331.7 | 332.2 | p = 0.103 | Model 1 | 0.018 | 0.00 | 0.138 |

Table Description: This table is a partial summary (also refer Table 2 in the main text) of relevant statistic derived from the linear mixed effects analysis involving two models that modelled fixed (Selection Regime, Sex and their interaction) and random (Block and Time) predictor variables on the response variable (Cuticular Hydrocarbon Peak Area) according to the following formulae:

**Model 1 = Selection Regime + Sex + (Selection regime x Sex) + (1|Block) + (1|Time)**

**Model 2 = Selection Regime + Sex + (1|Block) + (1|Time)**

The Selection Regime-Sex interaction term is the only difference between the two models. AIC stands for Akike Information Criterion and lower AIC value is a measure of better fit for a model. The difference between Model 1 and Model 2 in the column 3 of the table is based on ANOVA comparing the two models. The verbatim R output of the linear mixed effects analysis for each cuticular hydrocarbon peak is provided below in form of a R-Markdown document.

### Linear mixed effects analysis of Peak 1

```
Model1 = lmer(CHC_data$Peak_1 ~ Sel.Line + Sex + Sel.Line*Sex + (1|Blocks) + (1|Time), data = CHC_data, REML = FALSE)
summary(Model1)
## Linear mixed model fit by maximum likelihood  ['lmerMod']
## Formula: CHC_data$Peak_1 ~ Sel.Line + Sex + Sel.Line * Sex + (1 | Blocks) +
##     (1 | Time)
##    Data: CHC_data
##
##      AIC      BIC   logLik deviance df.resid
##    221.3    256.1   -101.6    203.3      343
##
## Scaled residuals:
##     Min      1Q  Median      3Q     Max
## -6.3904 -0.3437 -0.0662  0.3268  5.6321
##
## Random effects:
##  Groups   Name        Variance  Std.Dev.
##  Time     (Intercept) 6.812e-01 8.253e-01
##  Blocks   (Intercept) 2.200e-10 1.483e-05
##  Residual             9.409e-02 3.067e-01
## Number of obs: 352, groups:  Time, 6; Blocks, 3
##
## Fixed effects:
##                Estimate Std. Error t value
## (Intercept)     0.97490    0.33930   2.873
## Sel.LineF       0.10377    0.05702   1.820
## Sel.LineM      -0.09212    0.05649  -1.631
## Sexm           -0.01517    0.05649  -0.269
## Sel.LineF:Sexm -0.04740    0.08044  -0.589
## Sel.LineM:Sexm  0.19623    0.07971   2.462
##
## Correlation of Fixed Effects:
##             (Intr) Sl.LnF Sl.LnM Sexm   S.LF:S
## Sel.LineF   -0.083
## Sel.LineM   -0.083  0.496
## Sexm        -0.083  0.496  0.500
## Sel.LnF:Sxm  0.059 -0.709 -0.351 -0.703
## Sel.LnM:Sxm  0.059 -0.351 -0.709 -0.709  0.498
## convergence code: 0
## boundary (singular) fit: see ?isSingular
Anova(Model1)
## Analysis of Deviance Table (Type II Wald chisquare tests)
##
## Response: CHC_data$Peak_1
##                Chisq Df Pr(>Chisq)
## Sel.Line      4.8177  2   0.089919 .
## Sex           1.1912  1   0.275079
## Sel.Line:Sex 10.4380  2   0.005413 **
## ---
## Signif. codes:  0 '***' 0.001 '**' 0.01 '*' 0.05 '.' 0.1 ' ' 1
Model2 = lmer(CHC_data$Peak_1 ~ Sel.Line + Sex + (1|Blocks) + (1|Time), data = CHC_data, REML = FALSE)
summary(Model2)
## Linear mixed model fit by maximum likelihood  ['lmerMod']
## Formula: CHC_data$Peak_1 ~ Sel.Line + Sex + (1 | Blocks) + (1 | Time)
##    Data: CHC_data
##
##      AIC      BIC   logLik deviance df.resid
##    227.6    254.6   -106.8    213.6      345
##
## Scaled residuals:
##     Min      1Q  Median      3Q     Max
## -6.2099 -0.3454 -0.0270  0.2285  5.7100
##
## Random effects:
##  Groups   Name        Variance  Std.Dev.
##  Time     (Intercept) 6.801e-01 0.8246665
##  Blocks   (Intercept) 2.653e-08 0.0001629
##  Residual             9.693e-02 0.3113317
## Number of obs: 352, groups:  Time, 6; Blocks, 3
##
## Fixed effects:
##             Estimate Std. Error t value
## (Intercept) 0.949472   0.338294   2.807
## Sel.LineF   0.079635   0.040810   1.951
## Sel.LineM   0.006617   0.040454   0.164
## Sexm        0.035690   0.033190   1.075
##
## Correlation of Fixed Effects:
##           (Intr) Sl.LnF Sl.LnM
## Sel.LineF -0.059
## Sel.LineM -0.060  0.498
## Sexm      -0.049 -0.004 -0.003
Anova(Model2)
## Analysis of Deviance Table (Type II Wald chisquare tests)
##
## Response: CHC_data$Peak_1
##           Chisq Df Pr(>Chisq)
## Sel.Line 4.6762  2    0.09651 .
## Sex      1.1563  1    0.28222
## ---
## Signif. codes:  0 '***' 0.001 '**' 0.01 '*' 0.05 '.' 0.1 ' ' 1
anova(Model1, Model2)
## Data: CHC_data
## Models:
## Model2: CHC_data$Peak_1 ~ Sel.Line + Sex + (1 | Blocks) + (1 | Time)
## Model1: CHC_data$Peak_1 ~ Sel.Line + Sex + Sel.Line * Sex + (1 | Blocks) +
## Model1:     (1 | Time)
##        Df    AIC    BIC  logLik deviance  Chisq Chi Df Pr(>Chisq)
## Model2  7 227.57 254.62 -106.79   213.57
## Model1  9 221.29 256.06 -101.64   203.29 10.283      2   0.005848 **
## ---
## Signif. codes:  0 '***' 0.001 '**' 0.01 '*' 0.05 '.' 0.1 ' ' 1
```

### Linear mixed effects analysis of Peak 2

```
Model1 = lmer(CHC_data$Peak_2 ~ Sel.Line + Sex + Sel.Line*Sex + (1|Blocks) + (1|Time), data = CHC_data, REML = FALSE)
summary(Model1)
## Linear mixed model fit by maximum likelihood  ['lmerMod']
## Formula: CHC_data$Peak_2 ~ Sel.Line + Sex + Sel.Line * Sex + (1 | Blocks) +
##     (1 | Time)
##    Data: CHC_data
##
##      AIC      BIC   logLik deviance df.resid
##  -1251.4  -1216.6    634.7  -1269.4      343
##
## Scaled residuals:
##     Min      1Q  Median      3Q     Max
## -2.7787 -0.5436 -0.1695  0.2816  5.1915
##
## Random effects:
##  Groups   Name        Variance  Std.Dev.
##  Time     (Intercept) 8.043e-04 2.836e-02
##  Blocks   (Intercept) 4.150e-12 2.037e-06
##  Residual             1.498e-03 3.871e-02
## Number of obs: 352, groups:  Time, 6; Blocks, 3
##
## Fixed effects:
##                 Estimate Std. Error t value
## (Intercept)     0.090001   0.012628   7.127
## Sel.LineF       0.003178   0.007195   0.442
## Sel.LineM      -0.018460   0.007129  -2.589
## Sexm           -0.004859   0.007129  -0.682
## Sel.LineF:Sexm  0.007344   0.010151   0.723
## Sel.LineM:Sexm  0.023651   0.010060   2.351
##
## Correlation of Fixed Effects:
##             (Intr) Sl.LnF Sl.LnM Sexm   S.LF:S
## Sel.LineF   -0.280
## Sel.LineM   -0.282  0.496
## Sexm        -0.282  0.496  0.500
## Sel.LnF:Sxm  0.198 -0.709 -0.351 -0.703
## Sel.LnM:Sxm  0.200 -0.351 -0.709 -0.709  0.498
## convergence code: 0
## boundary (singular) fit: see ?isSingular
Anova(Model1)
## Analysis of Deviance Table (Type II Wald chisquare tests)
##
## Response: CHC_data$Peak_2
##               Chisq Df Pr(>Chisq)
## Sel.Line     7.0227  2    0.02986 *
## Sex          1.7997  1    0.17975
## Sel.Line:Sex 5.7934  2    0.05521 .
## ---
## Signif. codes:  0 '***' 0.001 '**' 0.01 '*' 0.05 '.' 0.1 ' ' 1
Model2 = lmer(CHC_data$Peak_2 ~ Sel.Line + Sex + (1|Blocks) + (1|Time), data = CHC_data, REML = FALSE)
summary(Model2)
## Linear mixed model fit by maximum likelihood  ['lmerMod']
## Formula: CHC_data$Peak_2 ~ Sel.Line + Sex + (1 | Blocks) + (1 | Time)
##    Data: CHC_data
##
##      AIC      BIC   logLik deviance df.resid
##  -1249.6  -1222.6    631.8  -1263.6      345
##
## Scaled residuals:
##     Min      1Q  Median      3Q     Max
## -2.6582 -0.4967 -0.1508  0.2683  5.1089
##
## Random effects:
##  Groups   Name        Variance  Std.Dev.
##  Time     (Intercept) 0.0008036 0.02835
##  Blocks   (Intercept) 0.0000000 0.00000
##  Residual             0.0015235 0.03903
## Number of obs: 352, groups:  Time, 6; Blocks, 3
##
## Fixed effects:
##              Estimate Std. Error t value
## (Intercept)  0.084802   0.012296   6.897
## Sel.LineF    0.006839   0.005116   1.337
## Sel.LineM   -0.006576   0.005072  -1.297
## Sexm         0.005536   0.004161   1.330
##
## Correlation of Fixed Effects:
##           (Intr) Sl.LnF Sl.LnM
## Sel.LineF -0.205
## Sel.LineM -0.207  0.498
## Sexm      -0.169 -0.004 -0.003
## convergence code: 0
## boundary (singular) fit: see ?isSingular
Anova(Model2)
## Analysis of Deviance Table (Type II Wald chisquare tests)
##
## Response: CHC_data$Peak_2
##          Chisq Df Pr(>Chisq)
## Sel.Line 6.907  2    0.03164 *
## Sex      1.770  1    0.18339
## ---
## Signif. codes:  0 '***' 0.001 '**' 0.01 '*' 0.05 '.' 0.1 ' ' 1
anova(Model1, Model2)
## Data: CHC_data
## Models:
## Model2: CHC_data$Peak_2 ~ Sel.Line + Sex + (1 | Blocks) + (1 | Time)
## Model1: CHC_data$Peak_2 ~ Sel.Line + Sex + Sel.Line * Sex + (1 | Blocks) +
## Model1:     (1 | Time)
##        Df     AIC     BIC logLik deviance  Chisq Chi Df Pr(>Chisq)
## Model2  7 -1249.6 -1222.6 631.81  -1263.6
## Model1  9 -1251.4 -1216.6 634.69  -1269.4 5.7454      2    0.05655 .
## ---
## Signif. codes:  0 '***' 0.001 '**' 0.01 '*' 0.05 '.' 0.1 ' ' 1
```

### Linear mixed effects analysis of Peak 3

```
Model1 = lmer(CHC_data$Peak_3 ~ Sel.Line + Sex + Sel.Line*Sex + (1|Blocks) + (1|Time), data = CHC_data, REML = FALSE)
summary(Model1)
## Linear mixed model fit by maximum likelihood  ['lmerMod']
## Formula: CHC_data$Peak_3 ~ Sel.Line + Sex + Sel.Line * Sex + (1 | Blocks) +
##     (1 | Time)
##    Data: CHC_data
##
##      AIC      BIC   logLik deviance df.resid
##   -826.3   -791.6    422.2   -844.3      343
##
## Scaled residuals:
##     Min      1Q  Median      3Q     Max
## -5.3466 -0.4535 -0.0959  0.3329  8.6990
##
## Random effects:
##  Groups   Name        Variance  Std.Dev.
##  Time     (Intercept) 2.007e-02 1.417e-01
##  Blocks   (Intercept) 2.452e-11 4.952e-06
##  Residual             4.843e-03 6.959e-02
## Number of obs: 352, groups:  Time, 6; Blocks, 3
##
## Fixed effects:
##                 Estimate Std. Error t value
## (Intercept)     0.258734   0.058546   4.419
## Sel.LineF       0.022577   0.012935   1.745
## Sel.LineM      -0.015970   0.012816  -1.246
## Sexm            0.054792   0.012816   4.275
## Sel.LineF:Sexm -0.004517   0.018250  -0.248
## Sel.LineM:Sexm  0.023611   0.018085   1.306
##
## Correlation of Fixed Effects:
##             (Intr) Sl.LnF Sl.LnM Sexm   S.LF:S
## Sel.LineF   -0.109
## Sel.LineM   -0.109  0.496
## Sexm        -0.109  0.496  0.500
## Sel.LnF:Sxm  0.077 -0.709 -0.351 -0.703
## Sel.LnM:Sxm  0.078 -0.351 -0.709 -0.709  0.498
## convergence code: 0
## boundary (singular) fit: see ?isSingular
Anova(Model1)
## Analysis of Deviance Table (Type II Wald chisquare tests)
##
## Response: CHC_data$Peak_3
##                Chisq Df Pr(>Chisq)
## Sel.Line      8.1691  2    0.01683 *
## Sex          68.2747  1    < 2e-16 ***
## Sel.Line:Sex  2.7755  2    0.24964
## ---
## Signif. codes:  0 '***' 0.001 '**' 0.01 '*' 0.05 '.' 0.1 ' ' 1
Model2 = lmer(CHC_data$Peak_3 ~ Sel.Line + Sex  + (1|Blocks) + (1|Time), data = CHC_data, REML = FALSE)
summary(Model2)
## Linear mixed model fit by maximum likelihood  ['lmerMod']
## Formula: CHC_data$Peak_3 ~ Sel.Line + Sex + (1 | Blocks) + (1 | Time)
##    Data: CHC_data
##
##      AIC      BIC   logLik deviance df.resid
##   -827.6   -800.5    420.8   -841.6      345
##
## Scaled residuals:
##     Min      1Q  Median      3Q     Max
## -5.2766 -0.4935 -0.0924  0.3453  8.5827
##
## Random effects:
##  Groups   Name        Variance Std.Dev.
##  Time     (Intercept) 0.020056 0.14162
##  Blocks   (Intercept) 0.000000 0.00000
##  Residual             0.004882 0.06987
## Number of obs: 352, groups:  Time, 6; Blocks, 3
##
## Fixed effects:
##              Estimate Std. Error t value
## (Intercept)  0.255481   0.058291   4.383
## Sel.LineF    0.020269   0.009158   2.213
## Sel.LineM   -0.004092   0.009079  -0.451
## Sexm         0.061299   0.007448   8.230
##
## Correlation of Fixed Effects:
##           (Intr) Sl.LnF Sl.LnM
## Sel.LineF -0.077
## Sel.LineM -0.078  0.498
## Sexm      -0.064 -0.004 -0.003
## convergence code: 0
## boundary (singular) fit: see ?isSingular
Anova(Model2)
## Analysis of Deviance Table (Type II Wald chisquare tests)
##
## Response: CHC_data$Peak_3
##            Chisq Df Pr(>Chisq)
## Sel.Line  8.1039  2    0.01739 *
## Sex      67.7304  1    < 2e-16 ***
## ---
## Signif. codes:  0 '***' 0.001 '**' 0.01 '*' 0.05 '.' 0.1 ' ' 1
anova(Model1, Model2)
## Data: CHC_data
## Models:
## Model2: CHC_data$Peak_3 ~ Sel.Line + Sex + (1 | Blocks) + (1 | Time)
## Model1: CHC_data$Peak_3 ~ Sel.Line + Sex + Sel.Line * Sex + (1 | Blocks) +
## Model1:     (1 | Time)
##        Df     AIC     BIC logLik deviance  Chisq Chi Df Pr(>Chisq)
## Model2  7 -827.58 -800.53 420.79  -841.58
## Model1  9 -826.34 -791.57 422.17  -844.34 2.7643      2      0.251
```

### Linear mixed effects analysis of Peak 4

```
Model1 = lmer(CHC_data$Peak_4 ~ Sel.Line + Sex + Sel.Line*Sex + (1|Blocks) + (1|Time), data = CHC_data, REML = FALSE)
summary(Model1)
## Linear mixed model fit by maximum likelihood  ['lmerMod']
## Formula: CHC_data$Peak_4 ~ Sel.Line + Sex + Sel.Line * Sex + (1 | Blocks) +
##     (1 | Time)
##    Data: CHC_data
##
##      AIC      BIC   logLik deviance df.resid
##    381.7    416.5   -181.8    363.7      343
##
## Scaled residuals:
##     Min      1Q  Median      3Q     Max
## -2.9298 -0.6124 -0.1027  0.5072  3.8035
##
## Random effects:
##  Groups   Name        Variance  Std.Dev.
##  Time     (Intercept) 6.405e-02 0.2530714
##  Blocks   (Intercept) 6.913e-08 0.0002629
##  Residual             1.557e-01 0.3946332
## Number of obs: 352, groups:  Time, 6; Blocks, 3
##
## Fixed effects:
##                 Estimate Std. Error t value
## (Intercept)     0.319298   0.115394   2.767
## Sel.LineF       0.007451   0.073351   0.102
## Sel.LineM       0.001398   0.072678   0.019
## Sexm            1.564142   0.072678  21.522
## Sel.LineF:Sexm -0.035802   0.103493  -0.346
## Sel.LineM:Sexm -0.124506   0.102557  -1.214
##
## Correlation of Fixed Effects:
##             (Intr) Sl.LnF Sl.LnM Sexm   S.LF:S
## Sel.LineF   -0.312
## Sel.LineM   -0.315  0.496
## Sexm        -0.315  0.496  0.500
## Sel.LnF:Sxm  0.221 -0.709 -0.351 -0.703
## Sel.LnM:Sxm  0.223 -0.351 -0.709 -0.709  0.498
## convergence code: 0
## Model failed to converge with max|grad| = 0.00460684 (tol = 0.002, component 1)
Anova(Model1)
## Analysis of Deviance Table (Type II Wald chisquare tests)
##
## Response: CHC_data$Peak_4
##                  Chisq Df Pr(>Chisq)
## Sel.Line        1.6283  2     0.4430
## Sex          1288.8561  1     <2e-16 ***
## Sel.Line:Sex    1.5627  2     0.4578
## ---
## Signif. codes:  0 '***' 0.001 '**' 0.01 '*' 0.05 '.' 0.1 ' ' 1
Model2 = lmer(CHC_data$Peak_4 ~ Sel.Line + Sex  + (1|Blocks) + (1|Time), data = CHC_data, REML = FALSE)
summary(Model2)
## Linear mixed model fit by maximum likelihood  ['lmerMod']
## Formula: CHC_data$Peak_4 ~ Sel.Line + Sex + (1 | Blocks) + (1 | Time)
##    Data: CHC_data
##
##      AIC      BIC   logLik deviance df.resid
##    379.3    406.3   -182.6    365.3      345
##
## Scaled residuals:
##     Min      1Q  Median      3Q     Max
## -2.8990 -0.6059 -0.0985  0.5169  3.8629
##
## Random effects:
##  Groups   Name        Variance  Std.Dev.
##  Time     (Intercept) 6.420e-02 0.253369
##  Blocks   (Intercept) 1.643e-06 0.001282
##  Residual             1.564e-01 0.395515
## Number of obs: 352, groups:  Time, 6; Blocks, 3
##
## Fixed effects:
##             Estimate Std. Error t value
## (Intercept)  0.34620    0.11167    3.10
## Sel.LineF   -0.01038    0.05184   -0.20
## Sel.LineM   -0.06116    0.05139   -1.19
## Sexm         1.51035    0.04216   35.82
##
## Correlation of Fixed Effects:
##           (Intr) Sl.LnF Sl.LnM
## Sel.LineF -0.228
## Sel.LineM -0.230  0.498
## Sexm      -0.189 -0.004 -0.003
## convergence code: 0
## Model failed to converge with max|grad| = 0.00237822 (tol = 0.002, component 1)
Anova(Model2)
## Analysis of Deviance Table (Type II Wald chisquare tests)
##
## Response: CHC_data$Peak_4
##              Chisq Df Pr(>Chisq)
## Sel.Line    1.6211  2     0.4446
## Sex      1283.1154  1     <2e-16 ***
## ---
## Signif. codes:  0 '***' 0.001 '**' 0.01 '*' 0.05 '.' 0.1 ' ' 1
anova(Model1, Model2)
## Data: CHC_data
## Models:
## Model2: CHC_data$Peak_4 ~ Sel.Line + Sex + (1 | Blocks) + (1 | Time)
## Model1: CHC_data$Peak_4 ~ Sel.Line + Sex + Sel.Line * Sex + (1 | Blocks) +
## Model1:     (1 | Time)
##        Df    AIC    BIC  logLik deviance  Chisq Chi Df Pr(>Chisq)
## Model2  7 379.26 406.30 -182.63   365.26
## Model1  9 381.70 416.47 -181.85   363.70 1.5592      2     0.4586
```

### Linear mixed effects analysis of Peak 5

```
Model1 = lmer(CHC_data$Peak_5 ~ Sel.Line + Sex + Sel.Line*Sex + (1|Blocks) + (1|Time), data = CHC_data, REML = FALSE)
summary(Model1)
## Linear mixed model fit by maximum likelihood  ['lmerMod']
## Formula: CHC_data$Peak_5 ~ Sel.Line + Sex + Sel.Line * Sex + (1 | Blocks) +
##     (1 | Time)
##    Data: CHC_data
##
##      AIC      BIC   logLik deviance df.resid
##   -155.4   -120.7     86.7   -173.4      343
##
## Scaled residuals:
##     Min      1Q  Median      3Q     Max
## -2.6649 -0.4533 -0.1068  0.1610 11.7423
##
## Random effects:
##  Groups   Name        Variance Std.Dev.
##  Time     (Intercept) 0.0443   0.2105
##  Blocks   (Intercept) 0.0000   0.0000
##  Residual             0.0332   0.1822
## Number of obs: 352, groups:  Time, 6; Blocks, 3
##
## Fixed effects:
##                 Estimate Std. Error t value
## (Intercept)     0.393563   0.089140   4.415
## Sel.LineF       0.082332   0.033869   2.431
## Sel.LineM       0.040444   0.033558   1.205
## Sexm            0.003928   0.033558   0.117
## Sel.LineF:Sexm -0.040748   0.047787  -0.853
## Sel.LineM:Sexm  0.004565   0.047354   0.096
##
## Correlation of Fixed Effects:
##             (Intr) Sl.LnF Sl.LnM Sexm   S.LF:S
## Sel.LineF   -0.187
## Sel.LineM   -0.188  0.496
## Sexm        -0.188  0.496  0.500
## Sel.LnF:Sxm  0.132 -0.709 -0.351 -0.703
## Sel.LnM:Sxm  0.133 -0.351 -0.709 -0.709  0.498
## convergence code: 0
## boundary (singular) fit: see ?isSingular
Anova(Model1)
## Analysis of Deviance Table (Type II Wald chisquare tests)
##
## Response: CHC_data$Peak_5
##               Chisq Df Pr(>Chisq)
## Sel.Line     7.0565  2    0.02936 *
## Sex          0.1628  1    0.68658
## Sel.Line:Sex 1.0879  2    0.58044
## ---
## Signif. codes:  0 '***' 0.001 '**' 0.01 '*' 0.05 '.' 0.1 ' ' 1
Model2 = lmer(CHC_data$Peak_5 ~ Sel.Line + Sex  + (1|Blocks) + (1|Time), data = CHC_data, REML = FALSE)
summary(Model2)
## Linear mixed model fit by maximum likelihood  ['lmerMod']
## Formula: CHC_data$Peak_5 ~ Sel.Line + Sex + (1 | Blocks) + (1 | Time)
##    Data: CHC_data
##
##      AIC      BIC   logLik deviance df.resid
##   -158.4   -131.3     86.2   -172.4      345
##
## Scaled residuals:
##     Min      1Q  Median      3Q     Max
## -2.6897 -0.4278 -0.1123  0.1542 11.6761
##
## Random effects:
##  Groups   Name        Variance  Std.Dev.
##  Time     (Intercept) 4.416e-02 2.102e-01
##  Blocks   (Intercept) 9.384e-10 3.063e-05
##  Residual             3.331e-02 1.825e-01
## Number of obs: 352, groups:  Time, 6; Blocks, 3
##
## Fixed effects:
##              Estimate Std. Error t value
## (Intercept)  0.399452   0.087964   4.541
## Sel.LineF    0.061819   0.023923   2.584
## Sel.LineM    0.042792   0.023714   1.804
## Sexm        -0.007838   0.019456  -0.403
##
## Correlation of Fixed Effects:
##           (Intr) Sl.LnF Sl.LnM
## Sel.LineF -0.134
## Sel.LineM -0.135  0.498
## Sexm      -0.111 -0.004 -0.003
Anova(Model2)
## Analysis of Deviance Table (Type II Wald chisquare tests)
##
## Response: CHC_data$Peak_5
##           Chisq Df Pr(>Chisq)
## Sel.Line 7.0339  2    0.02969 *
## Sex      0.1623  1    0.68705
## ---
## Signif. codes:  0 '***' 0.001 '**' 0.01 '*' 0.05 '.' 0.1 ' ' 1
anova(Model1, Model2)
## Data: CHC_data
## Models:
## Model2: CHC_data$Peak_5 ~ Sel.Line + Sex + (1 | Blocks) + (1 | Time)
## Model1: CHC_data$Peak_5 ~ Sel.Line + Sex + Sel.Line * Sex + (1 | Blocks) +
## Model1:     (1 | Time)
##        Df     AIC     BIC logLik deviance  Chisq Chi Df Pr(>Chisq)
## Model2  7 -158.36 -131.31 86.178  -172.36
## Model1  9 -155.44 -120.67 86.721  -173.44 1.0861      2      0.581
```

### Linear mixed effects analysis of Peak 6

```
Model1 = lmer(CHC_data$Peak_6 ~ Sel.Line + Sex + Sel.Line*Sex + (1|Blocks) + (1|Time), data = CHC_data, REML = FALSE)
summary(Model1)
## Linear mixed model fit by maximum likelihood  ['lmerMod']
## Formula: CHC_data$Peak_6 ~ Sel.Line + Sex + Sel.Line * Sex + (1 | Blocks) +
##     (1 | Time)
##    Data: CHC_data
##
##      AIC      BIC   logLik deviance df.resid
##     14.5     49.3      1.7     -3.5      343
##
## Scaled residuals:
##     Min      1Q  Median      3Q     Max
## -7.0061 -0.3785 -0.1202  0.2092  7.8337
##
## Random effects:
##  Groups   Name        Variance  Std.Dev.
##  Time     (Intercept) 3.754e-01 0.6126672
##  Blocks   (Intercept) 6.685e-07 0.0008176
##  Residual             5.230e-02 0.2286914
## Number of obs: 352, groups:  Time, 6; Blocks, 3
##
## Fixed effects:
##                 Estimate Std. Error t value
## (Intercept)     0.785881   0.251888   3.120
## Sel.LineF       0.095927   0.042509   2.257
## Sel.LineM      -0.043906   0.042118  -1.042
## Sexm            0.007006   0.042118   0.166
## Sel.LineF:Sexm -0.080702   0.059976  -1.346
## Sel.LineM:Sexm  0.095852   0.059432   1.613
##
## Correlation of Fixed Effects:
##             (Intr) Sl.LnF Sl.LnM Sexm   S.LF:S
## Sel.LineF   -0.083
## Sel.LineM   -0.084  0.496
## Sexm        -0.084  0.496  0.500
## Sel.LnF:Sxm  0.059 -0.709 -0.351 -0.703
## Sel.LnM:Sxm  0.059 -0.351 -0.709 -0.709  0.498
## convergence code: 0
## Model failed to converge with max|grad| = 0.00528124 (tol = 0.002, component 1)
Anova(Model1)
## Analysis of Deviance Table (Type II Wald chisquare tests)
##
## Response: CHC_data$Peak_6
##               Chisq Df Pr(>Chisq)
## Sel.Line     4.1731  2    0.12412
## Sex          0.2867  1    0.59236
## Sel.Line:Sex 8.7385  2    0.01266 *
## ---
## Signif. codes:  0 '***' 0.001 '**' 0.01 '*' 0.05 '.' 0.1 ' ' 1
Model2 = lmer(CHC_data$Peak_6 ~ Sel.Line + Sex  + (1|Blocks) + (1|Time), data = CHC_data, REML = FALSE)
summary(Model2)
## Linear mixed model fit by maximum likelihood  ['lmerMod']
## Formula: CHC_data$Peak_6 ~ Sel.Line + Sex + (1 | Blocks) + (1 | Time)
##    Data: CHC_data
##
##      AIC      BIC   logLik deviance df.resid
##     19.2     46.2     -2.6      5.2      345
##
## Scaled residuals:
##     Min      1Q  Median      3Q     Max
## -6.8998 -0.4089 -0.1043  0.2033  7.5344
##
## Random effects:
##  Groups   Name        Variance Std.Dev.
##  Time     (Intercept) 0.37523  0.6126
##  Blocks   (Intercept) 0.00000  0.0000
##  Residual             0.05362  0.2316
## Number of obs: 352, groups:  Time, 6; Blocks, 3
##
## Fixed effects:
##             Estimate Std. Error t value
## (Intercept) 0.782869   0.251289   3.115
## Sel.LineF   0.055174   0.030353   1.818
## Sel.LineM   0.004398   0.030089   0.146
## Sexm        0.013054   0.024686   0.529
##
## Correlation of Fixed Effects:
##           (Intr) Sl.LnF Sl.LnM
## Sel.LineF -0.059
## Sel.LineM -0.060  0.498
## Sexm      -0.049 -0.004 -0.003
## convergence code: 0
## boundary (singular) fit: see ?isSingular
Anova(Model2)
## Analysis of Deviance Table (Type II Wald chisquare tests)
##
## Response: CHC_data$Peak_6
##           Chisq Df Pr(>Chisq)
## Sel.Line 4.0701  2     0.1307
## Sex      0.2796  1     0.5970
anova(Model1, Model2)
## Data: CHC_data
## Models:
## Model2: CHC_data$Peak_6 ~ Sel.Line + Sex + (1 | Blocks) + (1 | Time)
## Model1: CHC_data$Peak_6 ~ Sel.Line + Sex + Sel.Line * Sex + (1 | Blocks) +
## Model1:     (1 | Time)
##        Df    AIC    BIC  logLik deviance Chisq Chi Df Pr(>Chisq)
## Model2  7 19.164 46.210 -2.5821   5.1642
## Model1  9 14.534 49.307  1.7329  -3.4657  8.63      2    0.01337 *
## ---
## Signif. codes:  0 '***' 0.001 '**' 0.01 '*' 0.05 '.' 0.1 ' ' 1
```

### Linear mixed effects analysis of Peak 7

```
Model1 = lmer(CHC_data$Peak_7 ~ Sel.Line + Sex + Sel.Line*Sex + (1|Blocks) + (1|Time), data = CHC_data, REML = FALSE)
summary(Model1)
## Linear mixed model fit by maximum likelihood  ['lmerMod']
## Formula: CHC_data$Peak_7 ~ Sel.Line + Sex + Sel.Line * Sex + (1 | Blocks) +
##     (1 | Time)
##    Data: CHC_data
##
##      AIC      BIC   logLik deviance df.resid
##   -337.3   -302.6    177.7   -355.3      343
##
## Scaled residuals:
##     Min      1Q  Median      3Q     Max
## -2.1732 -0.6516 -0.1271  0.5678  9.1571
##
## Random effects:
##  Groups   Name        Variance Std.Dev.
##  Time     (Intercept) 0.006414 0.08009
##  Blocks   (Intercept) 0.003499 0.05915
##  Residual             0.020157 0.14198
## Number of obs: 352, groups:  Time, 6; Blocks, 3
##
## Fixed effects:
##                 Estimate Std. Error t value
## (Intercept)     0.144584   0.050766   2.848
## Sel.LineF       0.019206   0.026389   0.728
## Sel.LineM       0.015171   0.026147   0.580
## Sexm            0.411351   0.026147  15.732
## Sel.LineF:Sexm  0.007845   0.037233   0.211
## Sel.LineM:Sexm -0.016772   0.036896  -0.455
##
## Correlation of Fixed Effects:
##             (Intr) Sl.LnF Sl.LnM Sexm   S.LF:S
## Sel.LineF   -0.255
## Sel.LineM   -0.258  0.496
## Sexm        -0.258  0.496  0.500
## Sel.LnF:Sxm  0.181 -0.709 -0.351 -0.703
## Sel.LnM:Sxm  0.183 -0.351 -0.709 -0.709  0.498
Anova(Model1)
## Analysis of Deviance Table (Type II Wald chisquare tests)
##
## Response: CHC_data$Peak_7
##                 Chisq Df Pr(>Chisq)
## Sel.Line       1.6381  2     0.4409
## Sex          727.5234  1     <2e-16 ***
## Sel.Line:Sex   0.4606  2     0.7943
## ---
## Signif. codes:  0 '***' 0.001 '**' 0.01 '*' 0.05 '.' 0.1 ' ' 1
Model2 = lmer(CHC_data$Peak_7 ~ Sel.Line + Sex  + (1|Blocks) + (1|Time), data = CHC_data, REML = FALSE)
summary(Model2)
## Linear mixed model fit by maximum likelihood  ['lmerMod']
## Formula: CHC_data$Peak_7 ~ Sel.Line + Sex + (1 | Blocks) + (1 | Time)
##    Data: CHC_data
##
##      AIC      BIC   logLik deviance df.resid
##   -340.9   -313.8    177.4   -354.9      345
##
## Scaled residuals:
##     Min      1Q  Median      3Q     Max
## -2.1322 -0.6504 -0.1389  0.5695  9.1993
##
## Random effects:
##  Groups   Name        Variance Std.Dev.
##  Time     (Intercept) 0.006433 0.08021
##  Blocks   (Intercept) 0.003515 0.05929
##  Residual             0.020183 0.14207
## Number of obs: 352, groups:  Time, 6; Blocks, 3
##
## Fixed effects:
##             Estimate Std. Error t value
## (Intercept) 0.146137   0.049723   2.939
## Sel.LineF   0.023179   0.018622   1.245
## Sel.LineM   0.006727   0.018460   0.364
## Sexm        0.408243   0.015145  26.955
##
## Correlation of Fixed Effects:
##           (Intr) Sl.LnF Sl.LnM
## Sel.LineF -0.184
## Sel.LineM -0.186  0.498
## Sexm      -0.152 -0.004 -0.003
## convergence code: 0
## Model failed to converge with max|grad| = 0.00456241 (tol = 0.002, component 1)
Anova(Model2)
## Analysis of Deviance Table (Type II Wald chisquare tests)
##
## Response: CHC_data$Peak_7
##            Chisq Df Pr(>Chisq)
## Sel.Line   1.636  2     0.4413
## Sex      726.599  1     <2e-16 ***
## ---
## Signif. codes:  0 '***' 0.001 '**' 0.01 '*' 0.05 '.' 0.1 ' ' 1
anova(Model1, Model2)
## Data: CHC_data
## Models:
## Model2: CHC_data$Peak_7 ~ Sel.Line + Sex + (1 | Blocks) + (1 | Time)
## Model1: CHC_data$Peak_7 ~ Sel.Line + Sex + Sel.Line * Sex + (1 | Blocks) +
## Model1:     (1 | Time)
##        Df     AIC     BIC logLik deviance  Chisq Chi Df Pr(>Chisq)
## Model2  7 -340.88 -313.84 177.44  -354.88
## Model1  9 -337.34 -302.57 177.67  -355.34 0.4603      2     0.7944
```

### Linear mixed effects analysis of Peak 8

```
Model1 = lmer(CHC_data$Peak_8 ~ Sel.Line + Sex + Sel.Line*Sex + (1|Blocks) + (1|Time), data = CHC_data, REML = FALSE)
summary(Model1)
## Linear mixed model fit by maximum likelihood  ['lmerMod']
## Formula: CHC_data$Peak_8 ~ Sel.Line + Sex + Sel.Line * Sex + (1 | Blocks) +
##     (1 | Time)
##    Data: CHC_data
##
##      AIC      BIC   logLik deviance df.resid
##   1949.3   1984.0   -965.6   1931.3      343
##
## Scaled residuals:
##     Min      1Q  Median      3Q     Max
## -2.2351 -0.4271  0.0627  0.3530  5.9934
##
## Random effects:
##  Groups   Name        Variance Std.Dev.
##  Time     (Intercept)  1.132   1.064
##  Blocks   (Intercept)  0.000   0.000
##  Residual             13.716   3.704
## Number of obs: 352, groups:  Time, 6; Blocks, 3
##
## Fixed effects:
##                 Estimate Std. Error t value
## (Intercept)     0.065933   0.649033   0.102
## Sel.LineF      -0.031613   0.688315  -0.046
## Sel.LineM      -0.004053   0.682047  -0.006
## Sexm            7.079137   0.682048  10.379
## Sel.LineF:Sexm  1.313417   0.971177   1.352
## Sel.LineM:Sexm  0.671984   0.962458   0.698
##
## Correlation of Fixed Effects:
##             (Intr) Sl.LnF Sl.LnM Sexm   S.LF:S
## Sel.LineF   -0.521
## Sel.LineM   -0.526  0.496
## Sexm        -0.526  0.496  0.500
## Sel.LnF:Sxm  0.369 -0.709 -0.351 -0.703
## Sel.LnM:Sxm  0.372 -0.351 -0.709 -0.709  0.498
## convergence code: 0
## boundary (singular) fit: see ?isSingular
Anova(Model1)
## Analysis of Deviance Table (Type II Wald chisquare tests)
##
## Response: CHC_data$Peak_8
##                 Chisq Df Pr(>Chisq)
## Sel.Line       1.6779  2     0.4322
## Sex          383.8449  1     <2e-16 ***
## Sel.Line:Sex   1.8298  2     0.4006
## ---
## Signif. codes:  0 '***' 0.001 '**' 0.01 '*' 0.05 '.' 0.1 ' ' 1
Model2 = lmer(CHC_data$Peak_8 ~ Sel.Line + Sex  + (1|Blocks) + (1|Time), data = CHC_data, REML = FALSE)
summary(Model2)
## Linear mixed model fit by maximum likelihood  ['lmerMod']
## Formula: CHC_data$Peak_8 ~ Sel.Line + Sex + (1 | Blocks) + (1 | Time)
##    Data: CHC_data
##
##      AIC      BIC   logLik deviance df.resid
##   1947.1   1974.1   -966.5   1933.1      345
##
## Scaled residuals:
##     Min      1Q  Median      3Q     Max
## -2.1398 -0.3423  0.0355  0.3504  5.8928
##
## Random effects:
##  Groups   Name        Variance  Std.Dev.
##  Time     (Intercept) 1.121e+00 1.0585946
##  Blocks   (Intercept) 5.641e-09 0.0000751
##  Residual             1.379e+01 3.7135850
## Number of obs: 352, groups:  Time, 6; Blocks, 3
##
## Fixed effects:
##             Estimate Std. Error t value
## (Intercept)  -0.2623     0.5855  -0.448
## Sel.LineF     0.6283     0.4868   1.291
## Sel.LineM     0.3321     0.4825   0.688
## Sexm          7.7353     0.3959  19.539
##
## Correlation of Fixed Effects:
##           (Intr) Sl.LnF Sl.LnM
## Sel.LineF -0.409
## Sel.LineM -0.413  0.498
## Sexm      -0.338 -0.004 -0.003
## convergence code: 0
## boundary (singular) fit: see ?isSingular
Anova(Model2)
## Analysis of Deviance Table (Type II Wald chisquare tests)
##
## Response: CHC_data$Peak_8
##             Chisq Df Pr(>Chisq)
## Sel.Line   1.6688  2     0.4341
## Sex      381.7765  1     <2e-16 ***
## ---
## Signif. codes:  0 '***' 0.001 '**' 0.01 '*' 0.05 '.' 0.1 ' ' 1
anova(Model1, Model2)
## Data: CHC_data
## Models:
## Model2: CHC_data$Peak_8 ~ Sel.Line + Sex + (1 | Blocks) + (1 | Time)
## Model1: CHC_data$Peak_8 ~ Sel.Line + Sex + Sel.Line * Sex + (1 | Blocks) +
## Model1:     (1 | Time)
##        Df    AIC    BIC  logLik deviance  Chisq Chi Df Pr(>Chisq)
## Model2  7 1947.1 1974.1 -966.55   1933.1
## Model1  9 1949.3 1984.0 -965.63   1931.3 1.8246      2     0.4016
```

### Linear mixed effects analysis of Peak 9

```
Model1 = lmer(CHC_data$Peak_9 ~ Sel.Line + Sex + Sel.Line*Sex + (1|Blocks) + (1|Time), data = CHC_data, REML = FALSE)
summary(Model1)
## Linear mixed model fit by maximum likelihood  ['lmerMod']
## Formula: CHC_data$Peak_9 ~ Sel.Line + Sex + Sel.Line * Sex + (1 | Blocks) +
##     (1 | Time)
##    Data: CHC_data
##
##      AIC      BIC   logLik deviance df.resid
##    208.1    242.8    -95.0    190.1      343
##
## Scaled residuals:
##     Min      1Q  Median      3Q     Max
## -3.3281 -0.5804 -0.1318  0.5448  7.1126
##
## Random effects:
##  Groups   Name        Variance Std.Dev.
##  Time     (Intercept) 0.06844  0.2616
##  Blocks   (Intercept) 0.00000  0.0000
##  Residual             0.09421  0.3069
## Number of obs: 352, groups:  Time, 6; Blocks, 3
##
## Fixed effects:
##                Estimate Std. Error t value
## (Intercept)     0.63925    0.11404   5.606
## Sel.LineF       0.03665    0.05705   0.642
## Sel.LineM      -0.01050    0.05653  -0.186
## Sexm            1.24157    0.05653  21.964
## Sel.LineF:Sexm -0.04233    0.08049  -0.526
## Sel.LineM:Sexm -0.14399    0.07977  -1.805
##
## Correlation of Fixed Effects:
##             (Intr) Sl.LnF Sl.LnM Sexm   S.LF:S
## Sel.LineF   -0.246
## Sel.LineM   -0.248  0.496
## Sexm        -0.248  0.496  0.500
## Sel.LnF:Sxm  0.174 -0.709 -0.351 -0.703
## Sel.LnM:Sxm  0.176 -0.351 -0.709 -0.709  0.498
## convergence code: 0
## boundary (singular) fit: see ?isSingular
Anova(Model1)
## Analysis of Deviance Table (Type II Wald chisquare tests)
##
## Response: CHC_data$Peak_9
##                  Chisq Df Pr(>Chisq)
## Sel.Line        7.0002  2    0.03019 *
## Sex          1298.4240  1    < 2e-16 ***
## Sel.Line:Sex    3.4433  2    0.17877
## ---
## Signif. codes:  0 '***' 0.001 '**' 0.01 '*' 0.05 '.' 0.1 ' ' 1
Model2 = lmer(CHC_data$Peak_9 ~ Sel.Line + Sex  + (1|Blocks) + (1|Time), data = CHC_data, REML = FALSE)
summary(Model2)
## Linear mixed model fit by maximum likelihood  ['lmerMod']
## Formula: CHC_data$Peak_9 ~ Sel.Line + Sex + (1 | Blocks) + (1 | Time)
##    Data: CHC_data
##
##      AIC      BIC   logLik deviance df.resid
##    207.5    234.5    -96.8    193.5      345
##
## Scaled residuals:
##     Min      1Q  Median      3Q     Max
## -3.2766 -0.5788 -0.1335  0.5378  7.2116
##
## Random effects:
##  Groups   Name        Variance  Std.Dev.
##  Time     (Intercept) 6.828e-02 2.613e-01
##  Blocks   (Intercept) 1.005e-11 3.171e-06
##  Residual             9.515e-02 3.085e-01
## Number of obs: 352, groups:  Time, 6; Blocks, 3
##
## Fixed effects:
##             Estimate Std. Error t value
## (Intercept)  0.67051    0.11161   6.008
## Sel.LineF    0.01555    0.04043   0.385
## Sel.LineM   -0.08285    0.04008  -2.067
## Sexm         1.17906    0.03288  35.855
##
## Correlation of Fixed Effects:
##           (Intr) Sl.LnF Sl.LnM
## Sel.LineF -0.178
## Sel.LineM -0.180  0.498
## Sexm      -0.147 -0.004 -0.003
## convergence code: 0
## boundary (singular) fit: see ?isSingular
Anova(Model2)
## Analysis of Deviance Table (Type II Wald chisquare tests)
##
## Response: CHC_data$Peak_9
##              Chisq Df Pr(>Chisq)
## Sel.Line    6.9308  2    0.03126 *
## Sex      1285.5828  1    < 2e-16 ***
## ---
## Signif. codes:  0 '***' 0.001 '**' 0.01 '*' 0.05 '.' 0.1 ' ' 1
anova(Model1, Model2)
## Data: CHC_data
## Models:
## Model2: CHC_data$Peak_9 ~ Sel.Line + Sex + (1 | Blocks) + (1 | Time)
## Model1: CHC_data$Peak_9 ~ Sel.Line + Sex + Sel.Line * Sex + (1 | Blocks) +
## Model1:     (1 | Time)
##        Df    AIC    BIC  logLik deviance  Chisq Chi Df Pr(>Chisq)
## Model2  7 207.50 234.55 -96.750   193.50
## Model1  9 208.07 242.85 -95.037   190.07 3.4261      2     0.1803
```

### Linear mixed effects analysis of Peak 10

```
Model1 = lmer(CHC_data$Peak_10 ~ Sel.Line + Sex + Sel.Line*Sex + (1|Blocks) + (1|Time), data = CHC_data, REML = FALSE)
summary(Model1)
## Linear mixed model fit by maximum likelihood  ['lmerMod']
## Formula: CHC_data$Peak_10 ~ Sel.Line + Sex + Sel.Line * Sex + (1 | Blocks) +
##     (1 | Time)
##    Data: CHC_data
##
##      AIC      BIC   logLik deviance df.resid
##   -117.9    -83.1     67.9   -135.9      343
##
## Scaled residuals:
##     Min      1Q  Median      3Q     Max
## -1.9474 -0.6020 -0.1392  0.3815  6.7254
##
## Random effects:
##  Groups   Name        Variance Std.Dev.
##  Time     (Intercept) 0.006667 0.08165
##  Blocks   (Intercept) 0.003372 0.05807
##  Residual             0.037974 0.19487
## Number of obs: 352, groups:  Time, 6; Blocks, 3
##
## Fixed effects:
##                Estimate Std. Error t value
## (Intercept)     0.48535    0.05366   9.045
## Sel.LineF       0.01384    0.03622   0.382
## Sel.LineM      -0.01991    0.03589  -0.555
## Sexm            0.04083    0.03589   1.138
## Sel.LineF:Sexm -0.01358    0.05110  -0.266
## Sel.LineM:Sexm  0.03024    0.05064   0.597
##
## Correlation of Fixed Effects:
##             (Intr) Sl.LnF Sl.LnM Sexm   S.LF:S
## Sel.LineF   -0.332
## Sel.LineM   -0.334  0.496
## Sexm        -0.334  0.496  0.500
## Sel.LnF:Sxm  0.235 -0.709 -0.351 -0.703
## Sel.LnM:Sxm  0.237 -0.351 -0.709 -0.709  0.498
Anova(Model1)
## Analysis of Deviance Table (Type II Wald chisquare tests)
##
## Response: CHC_data$Peak_10
##               Chisq Df Pr(>Chisq)
## Sel.Line     0.2113  2    0.89972
## Sex          5.0360  1    0.02483 *
## Sel.Line:Sex 0.7778  2    0.67780
## ---
## Signif. codes:  0 '***' 0.001 '**' 0.01 '*' 0.05 '.' 0.1 ' ' 1
Model2 = lmer(CHC_data$Peak_10 ~ Sel.Line + Sex  + (1|Blocks) + (1|Time), data = CHC_data, REML = FALSE)
summary(Model2)
## Linear mixed model fit by maximum likelihood  ['lmerMod']
## Formula: CHC_data$Peak_10 ~ Sel.Line + Sex + (1 | Blocks) + (1 | Time)
##    Data: CHC_data
##
##      AIC      BIC   logLik deviance df.resid
##   -121.1    -94.1     67.6   -135.1      345
##
## Scaled residuals:
##     Min      1Q  Median      3Q     Max
## -1.8836 -0.5773 -0.1271  0.4053  6.6683
##
## Random effects:
##  Groups   Name        Variance Std.Dev.
##  Time     (Intercept) 0.006706 0.08189
##  Blocks   (Intercept) 0.003333 0.05774
##  Residual             0.038059 0.19509
## Number of obs: 352, groups:  Time, 6; Blocks, 3
##
## Fixed effects:
##              Estimate Std. Error t value
## (Intercept)  0.482454   0.051572   9.355
## Sel.LineF    0.006961   0.025572   0.272
## Sel.LineM   -0.004692   0.025349  -0.185
## Sexm         0.046620   0.020797   2.242
##
## Correlation of Fixed Effects:
##           (Intr) Sl.LnF Sl.LnM
## Sel.LineF -0.244
## Sel.LineM -0.246  0.498
## Sexm      -0.202 -0.004 -0.003
Anova(Model2)
## Analysis of Deviance Table (Type II Wald chisquare tests)
##
## Response: CHC_data$Peak_10
##           Chisq Df Pr(>Chisq)
## Sel.Line 0.2108  2    0.89995
## Sex      5.0249  1    0.02499 *
## ---
## Signif. codes:  0 '***' 0.001 '**' 0.01 '*' 0.05 '.' 0.1 ' ' 1
anova(Model1, Model2)
## Data: CHC_data
## Models:
## Model2: CHC_data$Peak_10 ~ Sel.Line + Sex + (1 | Blocks) + (1 | Time)
## Model1: CHC_data$Peak_10 ~ Sel.Line + Sex + Sel.Line * Sex + (1 | Blocks) +
## Model1:     (1 | Time)
##        Df     AIC     BIC logLik deviance  Chisq Chi Df Pr(>Chisq)
## Model2  7 -121.12 -94.074 67.560  -135.12
## Model1  9 -117.90 -83.124 67.948  -135.90 0.7769      2     0.6781
```

### Linear mixed effects analysis of Peak 11

```
Model1 = lmer(CHC_data$Peak_11 ~ Sel.Line + Sex + Sel.Line*Sex + (1|Blocks) + (1|Time), data = CHC_data, REML = FALSE)
summary(Model1)
## Linear mixed model fit by maximum likelihood  ['lmerMod']
## Formula: CHC_data$Peak_11 ~ Sel.Line + Sex + Sel.Line * Sex + (1 | Blocks) +
##     (1 | Time)
##    Data: CHC_data
##
##      AIC      BIC   logLik deviance df.resid
##   1180.8   1215.6   -581.4   1162.8      343
##
## Scaled residuals:
##     Min      1Q  Median      3Q     Max
## -2.9682 -0.5607 -0.0861  0.4306  6.5237
##
## Random effects:
##  Groups   Name        Variance Std.Dev.
##  Time     (Intercept) 0.3475   0.5895
##  Blocks   (Intercept) 0.1279   0.3576
##  Residual             1.5154   1.2310
## Number of obs: 352, groups:  Time, 6; Blocks, 3
##
## Fixed effects:
##                Estimate Std. Error t value
## (Intercept)     0.37248    0.35532   1.048
## Sel.LineF       0.08272    0.22881   0.362
## Sel.LineM       0.02287    0.22671   0.101
## Sexm            4.75747    0.22671  20.985
## Sel.LineF:Sexm  0.44605    0.32283   1.382
## Sel.LineM:Sexm -0.14648    0.31991  -0.458
##
## Correlation of Fixed Effects:
##             (Intr) Sl.LnF Sl.LnM Sexm   S.LF:S
## Sel.LineF   -0.316
## Sel.LineM   -0.319  0.496
## Sexm        -0.319  0.496  0.500
## Sel.LnF:Sxm  0.224 -0.709 -0.351 -0.703
## Sel.LnM:Sxm  0.226 -0.351 -0.709 -0.709  0.498
## convergence code: 0
## Model failed to converge with max|grad| = 0.00202033 (tol = 0.002, component 1)
Anova(Model1)
## Analysis of Deviance Table (Type II Wald chisquare tests)
##
## Response: CHC_data$Peak_11
##                  Chisq Df Pr(>Chisq)
## Sel.Line        5.7731  2    0.05577 .
## Sex          1367.8679  1    < 2e-16 ***
## Sel.Line:Sex    3.6545  2    0.16086
## ---
## Signif. codes:  0 '***' 0.001 '**' 0.01 '*' 0.05 '.' 0.1 ' ' 1
Model2 = lmer(CHC_data$Peak_11 ~ Sel.Line + Sex  + (1|Blocks) + (1|Time), data = CHC_data, REML = FALSE)
summary(Model2)
## Linear mixed model fit by maximum likelihood  ['lmerMod']
## Formula: CHC_data$Peak_11 ~ Sel.Line + Sex + (1 | Blocks) + (1 | Time)
##    Data: CHC_data
##
##      AIC      BIC   logLik deviance df.resid
##   1180.5   1207.5   -583.2   1166.5      345
##
## Scaled residuals:
##     Min      1Q  Median      3Q     Max
## -2.8079 -0.5335 -0.0905  0.4332  6.6328
##
## Random effects:
##  Groups   Name        Variance Std.Dev.
##  Time     (Intercept) 0.3446   0.5871
##  Blocks   (Intercept) 0.1334   0.3653
##  Residual             1.5313   1.2375
## Number of obs: 352, groups:  Time, 6; Blocks, 3
##
## Fixed effects:
##             Estimate Std. Error t value
## (Intercept)  0.32433    0.34532   0.939
## Sel.LineF    0.30739    0.16221   1.895
## Sel.LineM   -0.05138    0.16079  -0.320
## Sexm         4.85364    0.13192  36.792
##
## Correlation of Fixed Effects:
##           (Intr) Sl.LnF Sl.LnM
## Sel.LineF -0.231
## Sel.LineM -0.233  0.498
## Sexm      -0.191 -0.004 -0.003
## convergence code: 0
## Model failed to converge with max|grad| = 0.0153708 (tol = 0.002, component 1)
Anova(Model2)
## Analysis of Deviance Table (Type II Wald chisquare tests)
##
## Response: CHC_data$Peak_11
##              Chisq Df Pr(>Chisq)
## Sel.Line    5.7132  2    0.05746 .
## Sex      1353.6463  1    < 2e-16 ***
## ---
## Signif. codes:  0 '***' 0.001 '**' 0.01 '*' 0.05 '.' 0.1 ' ' 1
anova(Model1, Model2)
## Data: CHC_data
## Models:
## Model2: CHC_data$Peak_11 ~ Sel.Line + Sex + (1 | Blocks) + (1 | Time)
## Model1: CHC_data$Peak_11 ~ Sel.Line + Sex + Sel.Line * Sex + (1 | Blocks) +
## Model1:     (1 | Time)
##        Df    AIC    BIC  logLik deviance  Chisq Chi Df Pr(>Chisq)
## Model2  7 1180.5 1207.5 -583.24   1166.5
## Model1  9 1180.8 1215.6 -581.42   1162.8 3.6351      2     0.1624
```

### Linear mixed effects analysis of Peak 12

```
Model1 = lmer(CHC_data$Peak_12 ~ Sel.Line + Sex + Sel.Line*Sex + (1|Blocks) + (1|Time), data = CHC_data, REML = FALSE)
summary(Model1)
## Linear mixed model fit by maximum likelihood  ['lmerMod']
## Formula: CHC_data$Peak_12 ~ Sel.Line + Sex + Sel.Line * Sex + (1 | Blocks) +
##     (1 | Time)
##    Data: CHC_data
##
##      AIC      BIC   logLik deviance df.resid
##   2502.7   2537.5  -1242.3   2484.7      343
##
## Scaled residuals:
##     Min      1Q  Median      3Q     Max
## -3.5291 -0.5802 -0.0588  0.5224  4.7621
##
## Random effects:
##  Groups   Name        Variance Std.Dev.
##  Time     (Intercept) 22.66    4.760
##  Blocks   (Intercept)  0.00    0.000
##  Residual             64.62    8.039
## Number of obs: 352, groups:  Time, 6; Blocks, 3
##
## Fixed effects:
##                Estimate Std. Error t value
## (Intercept)      5.1184     2.2075   2.319
## Sel.LineF        0.8515     1.4941   0.570
## Sel.LineM        0.6156     1.4804   0.416
## Sexm            40.3316     1.4804  27.243
## Sel.LineF:Sexm   2.9071     2.1081   1.379
## Sel.LineM:Sexm  -1.1550     2.0891  -0.553
##
## Correlation of Fixed Effects:
##             (Intr) Sl.LnF Sl.LnM Sexm   S.LF:S
## Sel.LineF   -0.333
## Sel.LineM   -0.335  0.496
## Sexm        -0.335  0.496  0.500
## Sel.LnF:Sxm  0.236 -0.709 -0.351 -0.703
## Sel.LnM:Sxm  0.238 -0.351 -0.709 -0.709  0.498
## convergence code: 0
## boundary (singular) fit: see ?isSingular
Anova(Model1)
## Analysis of Deviance Table (Type II Wald chisquare tests)
##
## Response: CHC_data$Peak_12
##                  Chisq Df Pr(>Chisq)
## Sel.Line        6.3392  2    0.04202 *
## Sex          2276.7721  1    < 2e-16 ***
## Sel.Line:Sex    3.9440  2    0.13918
## ---
## Signif. codes:  0 '***' 0.001 '**' 0.01 '*' 0.05 '.' 0.1 ' ' 1
Model2 = lmer(CHC_data$Peak_12 ~ Sel.Line + Sex  + (1|Blocks) + (1|Time), data = CHC_data, REML = FALSE)
summary(Model2)
## Linear mixed model fit by maximum likelihood  ['lmerMod']
## Formula: CHC_data$Peak_12 ~ Sel.Line + Sex + (1 | Blocks) + (1 | Time)
##    Data: CHC_data
##
##      AIC      BIC   logLik deviance df.resid
##   2502.6   2529.6  -1244.3   2488.6      345
##
## Scaled residuals:
##     Min      1Q  Median      3Q     Max
## -3.3609 -0.5569 -0.1036  0.5258  4.8833
##
## Random effects:
##  Groups   Name        Variance Std.Dev.
##  Time     (Intercept) 22.61    4.755
##  Blocks   (Intercept)  0.00    0.000
##  Residual             65.36    8.084
## Number of obs: 352, groups:  Time, 6; Blocks, 3
##
## Fixed effects:
##             Estimate Std. Error t value
## (Intercept)  4.83855    2.12314   2.279
## Sel.LineF    2.31605    1.05970   2.186
## Sel.LineM    0.03101    1.05047   0.030
## Sexm        40.89063    0.86185  47.445
##
## Correlation of Fixed Effects:
##           (Intr) Sl.LnF Sl.LnM
## Sel.LineF -0.246
## Sel.LineM -0.248  0.498
## Sexm      -0.203 -0.004 -0.003
## convergence code: 0
## boundary (singular) fit: see ?isSingular
Anova(Model2)
## Analysis of Deviance Table (Type II Wald chisquare tests)
##
## Response: CHC_data$Peak_12
##              Chisq Df Pr(>Chisq)
## Sel.Line    6.2674  2    0.04356 *
## Sex      2251.0431  1    < 2e-16 ***
## ---
## Signif. codes:  0 '***' 0.001 '**' 0.01 '*' 0.05 '.' 0.1 ' ' 1
anova(Model1, Model2)
## Data: CHC_data
## Models:
## Model2: CHC_data$Peak_12 ~ Sel.Line + Sex + (1 | Blocks) + (1 | Time)
## Model1: CHC_data$Peak_12 ~ Sel.Line + Sex + Sel.Line * Sex + (1 | Blocks) +
## Model1:     (1 | Time)
##        Df    AIC    BIC  logLik deviance  Chisq Chi Df Pr(>Chisq)
## Model2  7 2502.6 2529.7 -1244.3   2488.6
## Model1  9 2502.7 2537.4 -1242.3   2484.7 3.9216      2     0.1407
```

### Linear mixed effects analysis of Peak 13

```
Model1 = lmer(CHC_data$Peak_13 ~ Sel.Line + Sex + Sel.Line*Sex + (1|Blocks) + (1|Time), data = CHC_data, REML = FALSE)
summary(Model1)
## Linear mixed model fit by maximum likelihood  ['lmerMod']
## Formula: CHC_data$Peak_13 ~ Sel.Line + Sex + Sel.Line * Sex + (1 | Blocks) +
##     (1 | Time)
##    Data: CHC_data
##
##      AIC      BIC   logLik deviance df.resid
##    349.6    384.4   -165.8    331.6      343
##
## Scaled residuals:
##     Min      1Q  Median      3Q     Max
## -3.5235 -0.5641  0.0131  0.4518  5.3352
##
## Random effects:
##  Groups   Name        Variance  Std.Dev.
##  Time     (Intercept) 5.011e-02 2.239e-01
##  Blocks   (Intercept) 4.173e-11 6.460e-06
##  Residual             1.425e-01 3.775e-01
## Number of obs: 352, groups:  Time, 6; Blocks, 3
##
## Fixed effects:
##                Estimate Std. Error t value
## (Intercept)     0.36444    0.10378   3.512
## Sel.LineF       0.04093    0.07017   0.583
## Sel.LineM       0.01400    0.06953   0.201
## Sexm            1.70645    0.06953  24.542
## Sel.LineF:Sexm  0.15027    0.09901   1.518
## Sel.LineM:Sexm  0.03444    0.09812   0.351
##
## Correlation of Fixed Effects:
##             (Intr) Sl.LnF Sl.LnM Sexm   S.LF:S
## Sel.LineF   -0.332
## Sel.LineM   -0.335  0.496
## Sexm        -0.335  0.496  0.500
## Sel.LnF:Sxm  0.236 -0.709 -0.351 -0.703
## Sel.LnM:Sxm  0.237 -0.351 -0.709 -0.709  0.498
## convergence code: 0
## boundary (singular) fit: see ?isSingular
Anova(Model1)
## Analysis of Deviance Table (Type II Wald chisquare tests)
##
## Response: CHC_data$Peak_13
##                  Chisq Df Pr(>Chisq)
## Sel.Line        5.9262  2    0.05166 .
## Sex          1927.8060  1    < 2e-16 ***
## Sel.Line:Sex    2.5212  2    0.28349
## ---
## Signif. codes:  0 '***' 0.001 '**' 0.01 '*' 0.05 '.' 0.1 ' ' 1
Model2 = lmer(CHC_data$Peak_13 ~ Sel.Line + Sex  + (1|Blocks) + (1|Time), data = CHC_data, REML = FALSE)
summary(Model2)
## Linear mixed model fit by maximum likelihood  ['lmerMod']
## Formula: CHC_data$Peak_13 ~ Sel.Line + Sex + (1 | Blocks) + (1 | Time)
##    Data: CHC_data
##
##      AIC      BIC   logLik deviance df.resid
##    348.1    375.2   -167.1    334.1      345
##
## Scaled residuals:
##     Min      1Q  Median      3Q     Max
## -3.3907 -0.5917 -0.0076  0.4630  5.4334
##
## Random effects:
##  Groups   Name        Variance  Std.Dev.
##  Time     (Intercept) 4.992e-02 0.2234330
##  Blocks   (Intercept) 3.341e-08 0.0001828
##  Residual             1.436e-01 0.3789264
## Number of obs: 352, groups:  Time, 6; Blocks, 3
##
## Fixed effects:
##             Estimate Std. Error t value
## (Intercept)  0.33406    0.09973   3.350
## Sel.LineF    0.11650    0.04967   2.345
## Sel.LineM    0.03112    0.04924   0.632
## Sexm         1.76718    0.04040  43.746
##
## Correlation of Fixed Effects:
##           (Intr) Sl.LnF Sl.LnM
## Sel.LineF -0.245
## Sel.LineM -0.247  0.498
## Sexm      -0.203 -0.004 -0.003
## convergence code: 0
## Model failed to converge with max|grad| = 0.00215915 (tol = 0.002, component 1)
Anova(Model2)
## Analysis of Deviance Table (Type II Wald chisquare tests)
##
## Response: CHC_data$Peak_13
##              Chisq Df Pr(>Chisq)
## Sel.Line    5.8829  2    0.05279 .
## Sex      1913.7499  1    < 2e-16 ***
## ---
## Signif. codes:  0 '***' 0.001 '**' 0.01 '*' 0.05 '.' 0.1 ' ' 1
anova(Model1, Model2)
## Data: CHC_data
## Models:
## Model2: CHC_data$Peak_13 ~ Sel.Line + Sex + (1 | Blocks) + (1 | Time)
## Model1: CHC_data$Peak_13 ~ Sel.Line + Sex + Sel.Line * Sex + (1 | Blocks) +
## Model1:     (1 | Time)
##        Df    AIC    BIC  logLik deviance  Chisq Chi Df Pr(>Chisq)
## Model2  7 348.14 375.19 -167.07   334.14
## Model1  9 349.63 384.40 -165.81   331.63 2.5118      2     0.2848
```

### Linear mixed effects analysis of Peak 14

```
Model1 = lmer(CHC_data$Peak_14 ~ Sel.Line + Sex + Sel.Line*Sex + (1|Blocks) + (1|Time), data = CHC_data, REML = FALSE)
summary(Model1)
## Linear mixed model fit by maximum likelihood  ['lmerMod']
## Formula: CHC_data$Peak_14 ~ Sel.Line + Sex + Sel.Line * Sex + (1 | Blocks) +
##     (1 | Time)
##    Data: CHC_data
##
##      AIC      BIC   logLik deviance df.resid
##   1461.2   1496.0   -721.6   1443.2      343
##
## Scaled residuals:
##     Min      1Q  Median      3Q     Max
## -4.1535 -0.5908 -0.0426  0.5691  4.7887
##
## Random effects:
##  Groups   Name        Variance Std.Dev.
##  Time     (Intercept) 1.836    1.355
##  Blocks   (Intercept) 0.000    0.000
##  Residual             3.328    1.824
## Number of obs: 352, groups:  Time, 6; Blocks, 3
##
## Fixed effects:
##                Estimate Std. Error t value
## (Intercept)     5.88114    0.60205   9.769
## Sel.LineF       0.01981    0.33910   0.058
## Sel.LineM      -0.50222    0.33598  -1.495
## Sexm            6.06109    0.33598  18.040
## Sel.LineF:Sexm -0.37100    0.47844  -0.775
## Sel.LineM:Sexm -0.48301    0.47411  -1.019
##
## Correlation of Fixed Effects:
##             (Intr) Sl.LnF Sl.LnM Sexm   S.LF:S
## Sel.LineF   -0.277
## Sel.LineM   -0.279  0.496
## Sexm        -0.279  0.496  0.500
## Sel.LnF:Sxm  0.196 -0.709 -0.351 -0.703
## Sel.LnM:Sxm  0.198 -0.351 -0.709 -0.709  0.498
## convergence code: 0
## boundary (singular) fit: see ?isSingular
Anova(Model1)
## Analysis of Deviance Table (Type II Wald chisquare tests)
##
## Response: CHC_data$Peak_14
##                 Chisq Df Pr(>Chisq)
## Sel.Line      10.8718  2   0.004357 **
## Sex          882.1980  1  < 2.2e-16 ***
## Sel.Line:Sex   1.1336  2   0.567349
## ---
## Signif. codes:  0 '***' 0.001 '**' 0.01 '*' 0.05 '.' 0.1 ' ' 1
Model2 = lmer(CHC_data$Peak_14 ~ Sel.Line + Sex  + (1|Blocks) + (1|Time), data = CHC_data, REML = FALSE)
summary(Model2)
## Linear mixed model fit by maximum likelihood  ['lmerMod']
## Formula: CHC_data$Peak_14 ~ Sel.Line + Sex + (1 | Blocks) + (1 | Time)
##    Data: CHC_data
##
##      AIC      BIC   logLik deviance df.resid
##   1458.4   1485.4   -722.2   1444.4      345
##
## Scaled residuals:
##     Min      1Q  Median      3Q     Max
## -4.1697 -0.6035 -0.0459  0.5382  4.8601
##
## Random effects:
##  Groups   Name        Variance  Std.Dev.
##  Time     (Intercept) 1.836e+00 1.3549021
##  Blocks   (Intercept) 4.179e-07 0.0006465
##  Residual             3.339e+00 1.8273303
## Number of obs: 352, groups:  Time, 6; Blocks, 3
##
## Fixed effects:
##             Estimate Std. Error t value
## (Intercept)   6.0234     0.5863  10.274
## Sel.LineF    -0.1662     0.2395  -0.694
## Sel.LineM    -0.7446     0.2374  -3.136
## Sexm          5.7766     0.1948  29.653
##
## Correlation of Fixed Effects:
##           (Intr) Sl.LnF Sl.LnM
## Sel.LineF -0.201
## Sel.LineM -0.203  0.498
## Sexm      -0.166 -0.004 -0.003
Anova(Model2)
## Analysis of Deviance Table (Type II Wald chisquare tests)
##
## Response: CHC_data$Peak_14
##            Chisq Df Pr(>Chisq)
## Sel.Line  10.836  2   0.004435 **
## Sex      879.316  1  < 2.2e-16 ***
## ---
## Signif. codes:  0 '***' 0.001 '**' 0.01 '*' 0.05 '.' 0.1 ' ' 1
anova(Model1, Model2)
## Data: CHC_data
## Models:
## Model2: CHC_data$Peak_14 ~ Sel.Line + Sex + (1 | Blocks) + (1 | Time)
## Model1: CHC_data$Peak_14 ~ Sel.Line + Sex + Sel.Line * Sex + (1 | Blocks) +
## Model1:     (1 | Time)
##        Df    AIC    BIC  logLik deviance  Chisq Chi Df Pr(>Chisq)
## Model2  7 1458.4 1485.4 -722.18   1444.4
## Model1  9 1461.2 1496.0 -721.62   1443.2 1.1317      2     0.5679
```

### Linear mixed effects analysis of Peak 15

```
Model1 = lmer(CHC_data$Peak_15 ~ Sel.Line + Sex + Sel.Line*Sex + (1|Blocks) + (1|Time), data = CHC_data, REML = FALSE)
summary(Model1)
## Linear mixed model fit by maximum likelihood  ['lmerMod']
## Formula: CHC_data$Peak_15 ~ Sel.Line + Sex + Sel.Line * Sex + (1 | Blocks) +
##     (1 | Time)
##    Data: CHC_data
##
##      AIC      BIC   logLik deviance df.resid
##    758.7    793.5   -370.4    740.7      343
##
## Scaled residuals:
##     Min      1Q  Median      3Q     Max
## -3.2792 -0.4961 -0.0608  0.4010 10.7782
##
## Random effects:
##  Groups   Name        Variance  Std.Dev.
##  Time     (Intercept) 4.197e-01 0.6478318
##  Blocks   (Intercept) 8.259e-08 0.0002874
##  Residual             4.484e-01 0.6695904
## Number of obs: 352, groups:  Time, 6; Blocks, 3
##
## Fixed effects:
##                Estimate Std. Error t value
## (Intercept)     1.36378    0.27848   4.897
## Sel.LineF       0.35830    0.12446   2.879
## Sel.LineM      -0.03125    0.12332  -0.253
## Sexm            1.60098    0.12332  12.983
## Sel.LineF:Sexm -0.11929    0.17560  -0.679
## Sel.LineM:Sexm -0.05284    0.17401  -0.304
##
## Correlation of Fixed Effects:
##             (Intr) Sl.LnF Sl.LnM Sexm   S.LF:S
## Sel.LineF   -0.220
## Sel.LineM   -0.221  0.496
## Sexm        -0.221  0.496  0.500
## Sel.LnF:Sxm  0.156 -0.709 -0.351 -0.703
## Sel.LnM:Sxm  0.157 -0.351 -0.709 -0.709  0.498
Anova(Model1)
## Analysis of Deviance Table (Type II Wald chisquare tests)
##
## Response: CHC_data$Peak_15
##                 Chisq Df Pr(>Chisq)
## Sel.Line      18.9319  2  7.744e-05 ***
## Sex          467.9467  1  < 2.2e-16 ***
## Sel.Line:Sex   0.4631  2     0.7933
## ---
## Signif. codes:  0 '***' 0.001 '**' 0.01 '*' 0.05 '.' 0.1 ' ' 1
Model2 = lmer(CHC_data$Peak_15 ~ Sel.Line + Sex  + (1|Blocks) + (1|Time), data = CHC_data, REML = FALSE)
summary(Model2)
## Linear mixed model fit by maximum likelihood  ['lmerMod']
## Formula: CHC_data$Peak_15 ~ Sel.Line + Sex + (1 | Blocks) + (1 | Time)
##    Data: CHC_data
##
##      AIC      BIC   logLik deviance df.resid
##    755.2    782.2   -370.6    741.2      345
##
## Scaled residuals:
##     Min      1Q  Median      3Q     Max
## -3.2331 -0.4872 -0.0623  0.4059 10.8193
##
## Random effects:
##  Groups   Name        Variance  Std.Dev.
##  Time     (Intercept) 4.185e-01 0.6469073
##  Blocks   (Intercept) 3.072e-08 0.0001753
##  Residual             4.490e-01 0.6700544
## Number of obs: 352, groups:  Time, 6; Blocks, 3
##
## Fixed effects:
##             Estimate Std. Error t value
## (Intercept)  1.39221    0.27355   5.089
## Sel.LineF    0.29834    0.08783   3.397
## Sel.LineM   -0.05767    0.08707  -0.662
## Sexm         1.54415    0.07143  21.617
##
## Correlation of Fixed Effects:
##           (Intr) Sl.LnF Sl.LnM
## Sel.LineF -0.158
## Sel.LineM -0.159  0.498
## Sexm      -0.131 -0.004 -0.003
Anova(Model2)
## Analysis of Deviance Table (Type II Wald chisquare tests)
##
## Response: CHC_data$Peak_15
##            Chisq Df Pr(>Chisq)
## Sel.Line  18.905  2  7.847e-05 ***
## Sex      467.299  1  < 2.2e-16 ***
## ---
## Signif. codes:  0 '***' 0.001 '**' 0.01 '*' 0.05 '.' 0.1 ' ' 1
anova(Model1, Model2)
## Data: CHC_data
## Models:
## Model2: CHC_data$Peak_15 ~ Sel.Line + Sex + (1 | Blocks) + (1 | Time)
## Model1: CHC_data$Peak_15 ~ Sel.Line + Sex + Sel.Line * Sex + (1 | Blocks) +
## Model1:     (1 | Time)
##        Df    AIC    BIC  logLik deviance  Chisq Chi Df Pr(>Chisq)
## Model2  7 755.17 782.22 -370.59   741.17
## Model1  9 758.71 793.48 -370.35   740.71 0.4627      2     0.7934
```

### Linear mixed effects analysis of Peak 16

```
Model1 = lmer(CHC_data$Peak_16 ~ Sel.Line + Sex + Sel.Line*Sex + (1|Blocks) + (1|Time), data = CHC_data, REML = FALSE)
summary(Model1)
## Linear mixed model fit by maximum likelihood  ['lmerMod']
## Formula: CHC_data$Peak_16 ~ Sel.Line + Sex + Sel.Line * Sex + (1 | Blocks) +
##     (1 | Time)
##    Data: CHC_data
##
##      AIC      BIC   logLik deviance df.resid
##    593.0    627.8   -287.5    575.0      343
##
## Scaled residuals:
##     Min      1Q  Median      3Q     Max
## -3.2810 -0.6930  0.0787  0.7279  2.3443
##
## Random effects:
##  Groups   Name        Variance Std.Dev.
##  Time     (Intercept) 0.10543  0.3247
##  Blocks   (Intercept) 0.05653  0.2378
##  Residual             0.28258  0.5316
## Number of obs: 352, groups:  Time, 6; Blocks, 3
##
## Fixed effects:
##                Estimate Std. Error t value
## (Intercept)     1.63525    0.20300   8.056
## Sel.LineF      -0.01288    0.09881  -0.130
## Sel.LineM      -0.15228    0.09790  -1.555
## Sexm            0.67807    0.09790   6.926
## Sel.LineF:Sexm  0.03415    0.13941   0.245
## Sel.LineM:Sexm  0.09513    0.13815   0.689
##
## Correlation of Fixed Effects:
##             (Intr) Sl.LnF Sl.LnM Sexm   S.LF:S
## Sel.LineF   -0.239
## Sel.LineM   -0.241  0.496
## Sexm        -0.241  0.496  0.500
## Sel.LnF:Sxm  0.170 -0.709 -0.351 -0.703
## Sel.LnM:Sxm  0.171 -0.351 -0.709 -0.709  0.498
Anova(Model1)
## Analysis of Deviance Table (Type II Wald chisquare tests)
##
## Response: CHC_data$Peak_16
##                 Chisq Df Pr(>Chisq)
## Sel.Line       3.1673  2     0.2052
## Sex          162.0395  1     <2e-16 ***
## Sel.Line:Sex   0.4869  2     0.7839
## ---
## Signif. codes:  0 '***' 0.001 '**' 0.01 '*' 0.05 '.' 0.1 ' ' 1
Model2 = lmer(CHC_data$Peak_16 ~ Sel.Line + Sex  + (1|Blocks) + (1|Time), data = CHC_data, REML = FALSE)
summary(Model2)
## Linear mixed model fit by maximum likelihood  ['lmerMod']
## Formula: CHC_data$Peak_16 ~ Sel.Line + Sex + (1 | Blocks) + (1 | Time)
##    Data: CHC_data
##
##      AIC      BIC   logLik deviance df.resid
##    589.5    616.5   -287.7    575.5      345
##
## Scaled residuals:
##     Min      1Q  Median      3Q     Max
## -3.3187 -0.7294  0.0741  0.7310  2.2943
##
## Random effects:
##  Groups   Name        Variance Std.Dev.
##  Time     (Intercept) 0.10550  0.3248
##  Blocks   (Intercept) 0.05639  0.2375
##  Residual             0.28298  0.5320
## Number of obs: 352, groups:  Time, 6; Blocks, 3
##
## Fixed effects:
##              Estimate Std. Error t value
## (Intercept)  1.613584   0.198951   8.110
## Sel.LineF    0.004164   0.069729   0.060
## Sel.LineM   -0.104488   0.069122  -1.512
## Sexm         0.721386   0.056710  12.721
##
## Correlation of Fixed Effects:
##           (Intr) Sl.LnF Sl.LnM
## Sel.LineF -0.172
## Sel.LineM -0.174  0.498
## Sexm      -0.143 -0.004 -0.003
Anova(Model2)
## Analysis of Deviance Table (Type II Wald chisquare tests)
##
## Response: CHC_data$Peak_16
##             Chisq Df Pr(>Chisq)
## Sel.Line   3.1628  2     0.2057
## Sex      161.8114  1     <2e-16 ***
## ---
## Signif. codes:  0 '***' 0.001 '**' 0.01 '*' 0.05 '.' 0.1 ' ' 1
anova(Model1, Model2)
## Data: CHC_data
## Models:
## Model2: CHC_data$Peak_16 ~ Sel.Line + Sex + (1 | Blocks) + (1 | Time)
## Model1: CHC_data$Peak_16 ~ Sel.Line + Sex + Sel.Line * Sex + (1 | Blocks) +
## Model1:     (1 | Time)
##        Df    AIC    BIC  logLik deviance  Chisq Chi Df Pr(>Chisq)
## Model2  7 589.47 616.52 -287.74   575.47
## Model1  9 592.98 627.76 -287.49   574.98 0.4866      2      0.784
```

### Linear mixed effects analysis of Peak 18

```
Model1 = lmer(CHC_data$Peak_18 ~ Sel.Line + Sex + Sel.Line*Sex + (1|Blocks) + (1|Time), data = CHC_data, REML = FALSE)
summary(Model1)
## Linear mixed model fit by maximum likelihood  ['lmerMod']
## Formula: CHC_data$Peak_18 ~ Sel.Line + Sex + Sel.Line * Sex + (1 | Blocks) +
##     (1 | Time)
##    Data: CHC_data
##
##      AIC      BIC   logLik deviance df.resid
##   1069.4   1104.2   -525.7   1051.4      343
##
## Scaled residuals:
##     Min      1Q  Median      3Q     Max
## -2.6806 -0.5873 -0.0441  0.5179  4.2770
##
## Random effects:
##  Groups   Name        Variance Std.Dev.
##  Time     (Intercept) 0.2627   0.5126
##  Blocks   (Intercept) 0.0000   0.0000
##  Residual             1.1085   1.0528
## Number of obs: 352, groups:  Time, 6; Blocks, 3
##
## Fixed effects:
##                 Estimate Std. Error t value
## (Intercept)     1.810680   0.250180   7.238
## Sel.LineF       0.066678   0.195687   0.341
## Sel.LineM      -0.160451   0.193895  -0.828
## Sexm            1.946889   0.193895  10.041
## Sel.LineF:Sexm -0.197209   0.276101  -0.714
## Sel.LineM:Sexm -0.004784   0.273607  -0.017
##
## Correlation of Fixed Effects:
##             (Intr) Sl.LnF Sl.LnM Sexm   S.LF:S
## Sel.LineF   -0.384
## Sel.LineM   -0.388  0.496
## Sexm        -0.388  0.496  0.500
## Sel.LnF:Sxm  0.272 -0.709 -0.351 -0.703
## Sel.LnM:Sxm  0.275 -0.351 -0.709 -0.709  0.498
## convergence code: 0
## boundary (singular) fit: see ?isSingular
Anova(Model1)
## Analysis of Deviance Table (Type II Wald chisquare tests)
##
## Response: CHC_data$Peak_18
##                 Chisq Df Pr(>Chisq)
## Sel.Line       1.5812  2     0.4536
## Sex          280.8199  1     <2e-16 ***
## Sel.Line:Sex   0.6622  2     0.7181
## ---
## Signif. codes:  0 '***' 0.001 '**' 0.01 '*' 0.05 '.' 0.1 ' ' 1
Model2 = lmer(CHC_data$Peak_18 ~ Sel.Line + Sex  + (1|Blocks) + (1|Time), data = CHC_data, REML = FALSE)
summary(Model2)
## Linear mixed model fit by maximum likelihood  ['lmerMod']
## Formula: CHC_data$Peak_18 ~ Sel.Line + Sex + (1 | Blocks) + (1 | Time)
##    Data: CHC_data
##
##      AIC      BIC   logLik deviance df.resid
##   1066.0   1093.1   -526.0   1052.0      345
##
## Scaled residuals:
##     Min      1Q  Median      3Q     Max
## -2.6483 -0.5948 -0.0663  0.5140  4.3002
##
## Random effects:
##  Groups   Name        Variance  Std.Dev.
##  Time     (Intercept) 2.626e-01 0.5124486
##  Blocks   (Intercept) 2.595e-08 0.0001611
##  Residual             1.111e+00 1.0538420
## Number of obs: 352, groups:  Time, 6; Blocks, 3
##
## Fixed effects:
##             Estimate Std. Error t value
## (Intercept)  1.84373    0.23736   7.768
## Sel.LineF   -0.03256    0.13814  -0.236
## Sel.LineM   -0.16260    0.13693  -1.187
## Sexm         1.88085    0.11235  16.742
##
## Correlation of Fixed Effects:
##           (Intr) Sl.LnF Sl.LnM
## Sel.LineF -0.286
## Sel.LineM -0.289  0.498
## Sexm      -0.237 -0.004 -0.003
Anova(Model2)
## Analysis of Deviance Table (Type II Wald chisquare tests)
##
## Response: CHC_data$Peak_18
##             Chisq Df Pr(>Chisq)
## Sel.Line   1.5782  2     0.4543
## Sex      280.2818  1     <2e-16 ***
## ---
## Signif. codes:  0 '***' 0.001 '**' 0.01 '*' 0.05 '.' 0.1 ' ' 1
anova(Model1, Model2)
## Data: CHC_data
## Models:
## Model2: CHC_data$Peak_18 ~ Sel.Line + Sex + (1 | Blocks) + (1 | Time)
## Model1: CHC_data$Peak_18 ~ Sel.Line + Sex + Sel.Line * Sex + (1 | Blocks) +
## Model1:     (1 | Time)
##        Df    AIC    BIC  logLik deviance  Chisq Chi Df Pr(>Chisq)
## Model2  7 1066.0 1093.1 -526.02   1052.0
## Model1  9 1069.4 1104.2 -525.69   1051.4 0.6615      2     0.7184
```

### Linear mixed effects analysis of Peak 19

```
Model1 = lmer(CHC_data$Peak_19 ~ Sel.Line + Sex + Sel.Line*Sex + (1|Blocks) + (1|Time), data = CHC_data, REML = FALSE)
summary(Model1)
## Linear mixed model fit by maximum likelihood  ['lmerMod']
## Formula: CHC_data$Peak_19 ~ Sel.Line + Sex + Sel.Line * Sex + (1 | Blocks) +
##     (1 | Time)
##    Data: CHC_data
##
##      AIC      BIC   logLik deviance df.resid
##    602.7    637.5   -292.4    584.7      343
##
## Scaled residuals:
##     Min      1Q  Median      3Q     Max
## -2.1534 -0.5377 -0.1352  0.3656  6.9886
##
## Random effects:
##  Groups   Name        Variance Std.Dev.
##  Time     (Intercept) 0.03654  0.1911
##  Blocks   (Intercept) 0.01487  0.1220
##  Residual             0.29604  0.5441
## Number of obs: 352, groups:  Time, 6; Blocks, 3
##
## Fixed effects:
##                 Estimate Std. Error t value
## (Intercept)    -0.007380   0.126762  -0.058
## Sel.LineF       0.015569   0.101125   0.154
## Sel.LineM       0.008751   0.100201   0.087
## Sexm            1.980506   0.100201  19.765
## Sel.LineF:Sexm -0.407392   0.142682  -2.855
## Sel.LineM:Sexm -0.337877   0.141396  -2.390
##
## Correlation of Fixed Effects:
##             (Intr) Sl.LnF Sl.LnM Sexm   S.LF:S
## Sel.LineF   -0.392
## Sel.LineM   -0.395  0.496
## Sexm        -0.395  0.496  0.500
## Sel.LnF:Sxm  0.278 -0.709 -0.351 -0.703
## Sel.LnM:Sxm  0.280 -0.351 -0.709 -0.709  0.498
## convergence code: 0
## Model failed to converge with max|grad| = 0.00297408 (tol = 0.002, component 1)
Anova(Model1)
## Analysis of Deviance Table (Type II Wald chisquare tests)
##
## Response: CHC_data$Peak_19
##                 Chisq Df Pr(>Chisq)
## Sel.Line       8.2253  2   0.016364 *
## Sex          892.8708  1  < 2.2e-16 ***
## Sel.Line:Sex   9.3985  2   0.009102 **
## ---
## Signif. codes:  0 '***' 0.001 '**' 0.01 '*' 0.05 '.' 0.1 ' ' 1
Model2 = lmer(CHC_data$Peak_19 ~ Sel.Line + Sex  + (1|Blocks) + (1|Time), data = CHC_data, REML = FALSE)
summary(Model2)
## Linear mixed model fit by maximum likelihood  ['lmerMod']
## Formula: CHC_data$Peak_19 ~ Sel.Line + Sex + (1 | Blocks) + (1 | Time)
##    Data: CHC_data
##
##      AIC      BIC   logLik deviance df.resid
##      608      635     -297      594      345
##
## Scaled residuals:
##     Min      1Q  Median      3Q     Max
## -2.2092 -0.5876 -0.0150  0.3459  7.1238
##
## Random effects:
##  Groups   Name        Variance Std.Dev.
##  Time     (Intercept) 0.03580  0.1892
##  Blocks   (Intercept) 0.01508  0.1228
##  Residual             0.30412  0.5515
## Number of obs: 352, groups:  Time, 6; Blocks, 3
##
## Fixed effects:
##             Estimate Std. Error t value
## (Intercept)  0.11635    0.12015   0.968
## Sel.LineF   -0.18898    0.07228  -2.614
## Sel.LineM   -0.16064    0.07166  -2.242
## Sexm         1.73320    0.05879  29.481
##
## Correlation of Fixed Effects:
##           (Intr) Sl.LnF Sl.LnM
## Sel.LineF -0.296
## Sel.LineM -0.299  0.498
## Sexm      -0.245 -0.004 -0.003
Anova(Model2)
## Analysis of Deviance Table (Type II Wald chisquare tests)
##
## Response: CHC_data$Peak_19
##           Chisq Df Pr(>Chisq)
## Sel.Line   8.01  2    0.01822 *
## Sex      869.13  1    < 2e-16 ***
## ---
## Signif. codes:  0 '***' 0.001 '**' 0.01 '*' 0.05 '.' 0.1 ' ' 1
anova(Model1, Model2)
## Data: CHC_data
## Models:
## Model2: CHC_data$Peak_19 ~ Sel.Line + Sex + (1 | Blocks) + (1 | Time)
## Model1: CHC_data$Peak_19 ~ Sel.Line + Sex + Sel.Line * Sex + (1 | Blocks) +
## Model1:     (1 | Time)
##        Df    AIC    BIC  logLik deviance  Chisq Chi Df Pr(>Chisq)
## Model2  7 608.00 635.04 -297.00   594.00
## Model1  9 602.73 637.50 -292.36   584.73 9.2714      2   0.009699 **
## ---
## Signif. codes:  0 '***' 0.001 '**' 0.01 '*' 0.05 '.' 0.1 ' ' 1
```

### Linear mixed effects analysis of Peak 20

```
Model1 = lmer(CHC_data$Peak_20 ~ Sel.Line + Sex + Sel.Line*Sex + (1|Blocks) + (1|Time), data = CHC_data, REML = FALSE)
summary(Model1)
## Linear mixed model fit by maximum likelihood  ['lmerMod']
## Formula: CHC_data$Peak_20 ~ Sel.Line + Sex + Sel.Line * Sex + (1 | Blocks) +
##     (1 | Time)
##    Data: CHC_data
##
##      AIC      BIC   logLik deviance df.resid
##   1469.7   1504.5   -725.9   1451.7      343
##
## Scaled residuals:
##     Min      1Q  Median      3Q     Max
## -2.7443 -0.6026 -0.1262  0.4666  4.6805
##
## Random effects:
##  Groups   Name        Variance Std.Dev.
##  Time     (Intercept) 0.6678   0.8172
##  Blocks   (Intercept) 0.0000   0.0000
##  Residual             3.4680   1.8622
## Number of obs: 352, groups:  Time, 6; Blocks, 3
##
## Fixed effects:
##                Estimate Std. Error t value
## (Intercept)      8.6098     0.4125  20.874
## Sel.LineF        0.1993     0.3461   0.576
## Sel.LineM       -1.0230     0.3430  -2.983
## Sexm            -4.8595     0.3430 -14.169
## Sel.LineF:Sexm  -0.2097     0.4884  -0.429
## Sel.LineM:Sexm   0.7789     0.4840   1.609
##
## Correlation of Fixed Effects:
##             (Intr) Sl.LnF Sl.LnM Sexm   S.LF:S
## Sel.LineF   -0.412
## Sel.LineM   -0.416  0.496
## Sexm        -0.416  0.496  0.500
## Sel.LnF:Sxm  0.292 -0.709 -0.351 -0.703
## Sel.LnM:Sxm  0.295 -0.351 -0.709 -0.709  0.498
## convergence code: 0
## boundary (singular) fit: see ?isSingular
Anova(Model1)
## Analysis of Deviance Table (Type II Wald chisquare tests)
##
## Response: CHC_data$Peak_20
##                Chisq Df Pr(>Chisq)
## Sel.Line      10.547  2   0.005125 **
## Sex          552.067  1  < 2.2e-16 ***
## Sel.Line:Sex   4.604  2   0.100056
## ---
## Signif. codes:  0 '***' 0.001 '**' 0.01 '*' 0.05 '.' 0.1 ' ' 1
Model2 = lmer(CHC_data$Peak_20 ~ Sel.Line + Sex  + (1|Blocks) + (1|Time), data = CHC_data, REML = FALSE)
summary(Model2)
## Linear mixed model fit by maximum likelihood  ['lmerMod']
## Formula: CHC_data$Peak_20 ~ Sel.Line + Sex + (1 | Blocks) + (1 | Time)
##    Data: CHC_data
##
##      AIC      BIC   logLik deviance df.resid
##   1470.3   1497.3   -728.1   1456.3      345
##
## Scaled residuals:
##     Min      1Q  Median      3Q     Max
## -2.8808 -0.5668 -0.1059  0.5135  4.4921
##
## Random effects:
##  Groups   Name        Variance  Std.Dev.
##  Time     (Intercept) 6.714e-01 0.819397
##  Blocks   (Intercept) 7.090e-07 0.000842
##  Residual             3.514e+00 1.874500
## Number of obs: 352, groups:  Time, 6; Blocks, 3
##
## Fixed effects:
##             Estimate Std. Error t value
## (Intercept)  8.51238    0.38945  21.857
## Sel.LineF    0.09261    0.24571   0.377
## Sel.LineM   -0.63111    0.24357  -2.591
## Sexm        -4.66461    0.19983 -23.343
##
## Correlation of Fixed Effects:
##           (Intr) Sl.LnF Sl.LnM
## Sel.LineF -0.310
## Sel.LineM -0.313  0.498
## Sexm      -0.257 -0.004 -0.003
## convergence code: 0
## Model failed to converge with max|grad| = 0.00336749 (tol = 0.002, component 1)
Anova(Model2)
## Analysis of Deviance Table (Type II Wald chisquare tests)
##
## Response: CHC_data$Peak_20
##           Chisq Df Pr(>Chisq)
## Sel.Line  10.41  2   0.005489 **
## Sex      544.87  1  < 2.2e-16 ***
## ---
## Signif. codes:  0 '***' 0.001 '**' 0.01 '*' 0.05 '.' 0.1 ' ' 1
anova(Model1, Model2)
## Data: CHC_data
## Models:
## Model2: CHC_data$Peak_20 ~ Sel.Line + Sex + (1 | Blocks) + (1 | Time)
## Model1: CHC_data$Peak_20 ~ Sel.Line + Sex + Sel.Line * Sex + (1 | Blocks) +
## Model1:     (1 | Time)
##        Df    AIC    BIC  logLik deviance  Chisq Chi Df Pr(>Chisq)
## Model2  7 1470.3 1497.3 -728.15   1456.3
## Model1  9 1469.7 1504.5 -725.86   1451.7 4.5741      2     0.1016
```

### Linear mixed effects analysis of Peak 21

```
Model1 = lmer(CHC_data$Peak_21 ~ Sel.Line + Sex + Sel.Line*Sex + (1|Blocks) + (1|Time), data = CHC_data, REML = FALSE)
summary(Model1)
## Linear mixed model fit by maximum likelihood  ['lmerMod']
## Formula: CHC_data$Peak_21 ~ Sel.Line + Sex + Sel.Line * Sex + (1 | Blocks) +
##     (1 | Time)
##    Data: CHC_data
##
##      AIC      BIC   logLik deviance df.resid
##   2466.1   2500.9  -1224.1   2448.1      343
##
## Scaled residuals:
##     Min      1Q  Median      3Q     Max
## -2.3339 -0.4719 -0.1198  0.3943  4.4729
##
## Random effects:
##  Groups   Name        Variance Std.Dev.
##  Time     (Intercept)  5.429   2.330
##  Blocks   (Intercept)  0.000   0.000
##  Residual             59.474   7.712
## Number of obs: 352, groups:  Time, 6; Blocks, 3
##
## Fixed effects:
##                Estimate Std. Error t value
## (Intercept)      19.482      1.383  14.084
## Sel.LineF         2.738      1.433   1.910
## Sel.LineM        -1.644      1.420  -1.158
## Sexm            -12.401      1.420  -8.731
## Sel.LineF:Sexm   -1.380      2.022  -0.683
## Sel.LineM:Sexm    1.513      2.004   0.755
##
## Correlation of Fixed Effects:
##             (Intr) Sl.LnF Sl.LnM Sexm   S.LF:S
## Sel.LineF   -0.509
## Sel.LineM   -0.513  0.496
## Sexm        -0.513  0.496  0.500
## Sel.LnF:Sxm  0.361 -0.709 -0.351 -0.703
## Sel.LnM:Sxm  0.364 -0.351 -0.709 -0.709  0.498
## convergence code: 0
## boundary (singular) fit: see ?isSingular
Anova(Model1)
## Analysis of Deviance Table (Type II Wald chisquare tests)
##
## Response: CHC_data$Peak_21
##                 Chisq Df Pr(>Chisq)
## Sel.Line       8.8056  2    0.01224 *
## Sex          225.2805  1    < 2e-16 ***
## Sel.Line:Sex   2.0599  2    0.35703
## ---
## Signif. codes:  0 '***' 0.001 '**' 0.01 '*' 0.05 '.' 0.1 ' ' 1
Model2 = lmer(CHC_data$Peak_21 ~ Sel.Line + Sex  + (1|Blocks) + (1|Time), data = CHC_data, REML = FALSE)
summary(Model2)
## Linear mixed model fit by maximum likelihood  ['lmerMod']
## Formula: CHC_data$Peak_21 ~ Sel.Line + Sex + (1 | Blocks) + (1 | Time)
##    Data: CHC_data
##
##      AIC      BIC   logLik deviance df.resid
##   2464.2   2491.2  -1225.1   2450.2      345
##
## Scaled residuals:
##     Min      1Q  Median      3Q     Max
## -2.3937 -0.4533 -0.0939  0.4038  4.4611
##
## Random effects:
##  Groups   Name        Variance Std.Dev.
##  Time     (Intercept)  5.461   2.337
##  Blocks   (Intercept)  0.000   0.000
##  Residual             59.822   7.734
## Number of obs: 352, groups:  Time, 6; Blocks, 3
##
## Fixed effects:
##             Estimate Std. Error t value
## (Intercept)  19.4522     1.2599  15.440
## Sel.LineF     2.0414     1.0138   2.014
## Sel.LineM    -0.8816     1.0050  -0.877
## Sexm        -12.3398     0.8245 -14.966
##
## Correlation of Fixed Effects:
##           (Intr) Sl.LnF Sl.LnM
## Sel.LineF -0.396
## Sel.LineM -0.399  0.498
## Sexm      -0.327 -0.004 -0.003
## convergence code: 0
## boundary (singular) fit: see ?isSingular
Anova(Model2)
## Analysis of Deviance Table (Type II Wald chisquare tests)
##
## Response: CHC_data$Peak_21
##             Chisq Df Pr(>Chisq)
## Sel.Line   8.7543  2    0.01256 *
## Sex      223.9698  1    < 2e-16 ***
## ---
## Signif. codes:  0 '***' 0.001 '**' 0.01 '*' 0.05 '.' 0.1 ' ' 1
anova(Model1, Model2)
## Data: CHC_data
## Models:
## Model2: CHC_data$Peak_21 ~ Sel.Line + Sex + (1 | Blocks) + (1 | Time)
## Model1: CHC_data$Peak_21 ~ Sel.Line + Sex + Sel.Line * Sex + (1 | Blocks) +
## Model1:     (1 | Time)
##        Df    AIC    BIC  logLik deviance  Chisq Chi Df Pr(>Chisq)
## Model2  7 2464.2 2491.2 -1225.1   2450.2
## Model1  9 2466.1 2500.9 -1224.1   2448.1 2.0539      2     0.3581
```

### Linear mixed effects analysis of Peak 22

```
Model1 = lmer(CHC_data$Peak_22 ~ Sel.Line + Sex + Sel.Line*Sex + (1|Blocks) + (1|Time), data = CHC_data, REML = FALSE)
summary(Model1)
## Linear mixed model fit by maximum likelihood  ['lmerMod']
## Formula: CHC_data$Peak_22 ~ Sel.Line + Sex + Sel.Line * Sex + (1 | Blocks) +
##     (1 | Time)
##    Data: CHC_data
##
##      AIC      BIC   logLik deviance df.resid
##     41.0     75.7    -11.5     23.0      343
##
## Scaled residuals:
##     Min      1Q  Median      3Q     Max
## -2.9231 -0.5246 -0.0476  0.4343  4.1582
##
## Random effects:
##  Groups   Name        Variance Std.Dev.
##  Time     (Intercept) 0.00792  0.08899
##  Blocks   (Intercept) 0.00000  0.00000
##  Residual             0.06023  0.24542
## Number of obs: 352, groups:  Time, 6; Blocks, 3
##
## Fixed effects:
##                Estimate Std. Error t value
## (Intercept)     0.78601    0.04839  16.243
## Sel.LineF       0.11395    0.04561   2.498
## Sel.LineM      -0.10104    0.04520  -2.235
## Sexm           -0.65616    0.04520 -14.518
## Sel.LineF:Sexm -0.09512    0.06436  -1.478
## Sel.LineM:Sexm  0.09985    0.06378   1.566
##
## Correlation of Fixed Effects:
##             (Intr) Sl.LnF Sl.LnM Sexm   S.LF:S
## Sel.LineF   -0.463
## Sel.LineM   -0.467  0.496
## Sexm        -0.467  0.496  0.500
## Sel.LnF:Sxm  0.328 -0.709 -0.351 -0.703
## Sel.LnM:Sxm  0.331 -0.351 -0.709 -0.709  0.498
## convergence code: 0
## boundary (singular) fit: see ?isSingular
Anova(Model1)
## Analysis of Deviance Table (Type II Wald chisquare tests)
##
## Response: CHC_data$Peak_22
##                 Chisq Df Pr(>Chisq)
## Sel.Line      13.2640  2   0.001317 **
## Sex          623.8194  1  < 2.2e-16 ***
## Sel.Line:Sex   9.2261  2   0.009922 **
## ---
## Signif. codes:  0 '***' 0.001 '**' 0.01 '*' 0.05 '.' 0.1 ' ' 1
Model2 = lmer(CHC_data$Peak_22 ~ Sel.Line + Sex  + (1|Blocks) + (1|Time), data = CHC_data, REML = FALSE)
summary(Model2)
## Linear mixed model fit by maximum likelihood  ['lmerMod']
## Formula: CHC_data$Peak_22 ~ Sel.Line + Sex + (1 | Blocks) + (1 | Time)
##    Data: CHC_data
##
##      AIC      BIC   logLik deviance df.resid
##     46.1     73.1    -16.0     32.1      345
##
## Scaled residuals:
##     Min      1Q  Median      3Q     Max
## -2.8774 -0.4905 -0.0086  0.4204  4.1039
##
## Random effects:
##  Groups   Name        Variance Std.Dev.
##  Time     (Intercept) 0.007981 0.08934
##  Blocks   (Intercept) 0.000000 0.00000
##  Residual             0.061828 0.24865
## Number of obs: 352, groups:  Time, 6; Blocks, 3
##
## Fixed effects:
##             Estimate Std. Error t value
## (Intercept)  0.78468    0.04506  17.416
## Sel.LineF    0.06594    0.03259   2.023
## Sel.LineM   -0.05071    0.03231  -1.569
## Sexm        -0.65347    0.02651 -24.652
##
## Correlation of Fixed Effects:
##           (Intr) Sl.LnF Sl.LnM
## Sel.LineF -0.356
## Sel.LineM -0.359  0.498
## Sexm      -0.294 -0.004 -0.003
## convergence code: 0
## boundary (singular) fit: see ?isSingular
Anova(Model2)
## Analysis of Deviance Table (Type II Wald chisquare tests)
##
## Response: CHC_data$Peak_22
##            Chisq Df Pr(>Chisq)
## Sel.Line  12.922  2   0.001563 **
## Sex      607.723  1  < 2.2e-16 ***
## ---
## Signif. codes:  0 '***' 0.001 '**' 0.01 '*' 0.05 '.' 0.1 ' ' 1
anova(Model1, Model2)
## Data: CHC_data
## Models:
## Model2: CHC_data$Peak_22 ~ Sel.Line + Sex + (1 | Blocks) + (1 | Time)
## Model1: CHC_data$Peak_22 ~ Sel.Line + Sex + Sel.Line * Sex + (1 | Blocks) +
## Model1:     (1 | Time)
##        Df    AIC    BIC  logLik deviance  Chisq Chi Df Pr(>Chisq)
## Model2  7 46.067 73.112 -16.034   32.067
## Model1  9 40.961 75.733 -11.480   22.961 9.1065      2    0.01053 *
## ---
## Signif. codes:  0 '***' 0.001 '**' 0.01 '*' 0.05 '.' 0.1 ' ' 1
```

### Linear mixed effects analysis of Peak 23

```
Model1 = lmer(CHC_data$Peak_23 ~ Sel.Line + Sex + Sel.Line*Sex + (1|Blocks) + (1|Time), data = CHC_data, REML = FALSE)
summary(Model1)
## Linear mixed model fit by maximum likelihood  ['lmerMod']
## Formula: CHC_data$Peak_23 ~ Sel.Line + Sex + Sel.Line * Sex + (1 | Blocks) +
##     (1 | Time)
##    Data: CHC_data
##
##      AIC      BIC   logLik deviance df.resid
##   1247.5   1282.3   -614.8   1229.5      343
##
## Scaled residuals:
##     Min      1Q  Median      3Q     Max
## -2.7429 -0.5573 -0.0237  0.4707  3.7132
##
## Random effects:
##  Groups   Name        Variance  Std.Dev.
##  Time     (Intercept) 7.165e-01 8.464e-01
##  Blocks   (Intercept) 2.076e-09 4.556e-05
##  Residual             1.824e+00 1.350e+00
## Number of obs: 352, groups:  Time, 6; Blocks, 3
##
## Fixed effects:
##                Estimate Std. Error t value
## (Intercept)      8.6201     0.3877  22.231
## Sel.LineF        0.2520     0.2510   1.004
## Sel.LineM       -0.8977     0.2487  -3.609
## Sexm            -6.1357     0.2487 -24.671
## Sel.LineF:Sexm  -0.2689     0.3542  -0.759
## Sel.LineM:Sexm   0.6455     0.3510   1.839
##
## Correlation of Fixed Effects:
##             (Intr) Sl.LnF Sl.LnM Sexm   S.LF:S
## Sel.LineF   -0.318
## Sel.LineM   -0.321  0.496
## Sexm        -0.321  0.496  0.500
## Sel.LnF:Sxm  0.225 -0.709 -0.351 -0.703
## Sel.LnM:Sxm  0.227 -0.351 -0.709 -0.709  0.498
## convergence code: 0
## boundary (singular) fit: see ?isSingular
Anova(Model1)
## Analysis of Deviance Table (Type II Wald chisquare tests)
##
## Response: CHC_data$Peak_23
##                  Chisq Df Pr(>Chisq)
## Sel.Line       17.5622  2  0.0001536 ***
## Sex          1740.0229  1  < 2.2e-16 ***
## Sel.Line:Sex    7.1125  2  0.0285458 *
## ---
## Signif. codes:  0 '***' 0.001 '**' 0.01 '*' 0.05 '.' 0.1 ' ' 1
Model2 = lmer(CHC_data$Peak_23 ~ Sel.Line + Sex  + (1|Blocks) + (1|Time), data = CHC_data, REML = FALSE)
summary(Model2)
## Linear mixed model fit by maximum likelihood  ['lmerMod']
## Formula: CHC_data$Peak_23 ~ Sel.Line + Sex + (1 | Blocks) + (1 | Time)
##    Data: CHC_data
##
##      AIC      BIC   logLik deviance df.resid
##   1250.6   1277.6   -618.3   1236.6      345
##
## Scaled residuals:
##     Min      1Q  Median      3Q     Max
## -2.8165 -0.4998 -0.0362  0.5219  3.7275
##
## Random effects:
##  Groups   Name        Variance  Std.Dev.
##  Time     (Intercept) 7.212e-01 8.492e-01
##  Blocks   (Intercept) 2.075e-09 4.555e-05
##  Residual             1.861e+00 1.364e+00
## Number of obs: 352, groups:  Time, 6; Blocks, 3
##
## Fixed effects:
##             Estimate Std. Error t value
## (Intercept)   8.5549     0.3759  22.761
## Sel.LineF     0.1157     0.1788   0.647
## Sel.LineM    -0.5728     0.1773  -3.231
## Sexm         -6.0053     0.1454 -41.294
##
## Correlation of Fixed Effects:
##           (Intr) Sl.LnF Sl.LnM
## Sel.LineF -0.234
## Sel.LineM -0.236  0.498
## Sexm      -0.193 -0.004 -0.003
## convergence code: 0
## boundary (singular) fit: see ?isSingular
Anova(Model2)
## Analysis of Deviance Table (Type II Wald chisquare tests)
##
## Response: CHC_data$Peak_23
##             Chisq Df Pr(>Chisq)
## Sel.Line   17.211  2  0.0001831 ***
## Sex      1705.186  1  < 2.2e-16 ***
## ---
## Signif. codes:  0 '***' 0.001 '**' 0.01 '*' 0.05 '.' 0.1 ' ' 1
anova(Model1, Model2)
## Data: CHC_data
## Models:
## Model2: CHC_data$Peak_23 ~ Sel.Line + Sex + (1 | Blocks) + (1 | Time)
## Model1: CHC_data$Peak_23 ~ Sel.Line + Sex + Sel.Line * Sex + (1 | Blocks) +
## Model1:     (1 | Time)
##        Df    AIC    BIC  logLik deviance  Chisq Chi Df Pr(>Chisq)
## Model2  7 1250.6 1277.6 -618.28   1236.6
## Model1  9 1247.5 1282.3 -614.76   1229.5 7.0411      2    0.02958 *
## ---
## Signif. codes:  0 '***' 0.001 '**' 0.01 '*' 0.05 '.' 0.1 ' ' 1
```

### Linear mixed effects analysis of Peak 24

```
Model1 = lmer(CHC_data$Peak_24 ~ Sel.Line + Sex + Sel.Line*Sex + (1|Blocks) + (1|Time), data = CHC_data, REML = FALSE)
summary(Model1)
## Linear mixed model fit by maximum likelihood  ['lmerMod']
## Formula: CHC_data$Peak_24 ~ Sel.Line + Sex + Sel.Line * Sex + (1 | Blocks) +
##     (1 | Time)
##    Data: CHC_data
##
##      AIC      BIC   logLik deviance df.resid
##   -552.6   -517.8    285.3   -570.6      343
##
## Scaled residuals:
##     Min      1Q  Median      3Q     Max
## -3.0337 -0.3388 -0.1013  0.2053  9.2737
##
## Random effects:
##  Groups   Name        Variance Std.Dev.
##  Time     (Intercept) 0.002597 0.05096
##  Blocks   (Intercept) 0.001364 0.03694
##  Residual             0.010992 0.10484
## Number of obs: 352, groups:  Time, 6; Blocks, 3
##
## Fixed effects:
##                 Estimate Std. Error t value
## (Intercept)     0.229613   0.032773   7.006
## Sel.LineF       0.008977   0.019487   0.461
## Sel.LineM      -0.008357   0.019308  -0.433
## Sexm           -0.007593   0.019308  -0.393
## Sel.LineF:Sexm -0.009257   0.027494  -0.337
## Sel.LineM:Sexm  0.012005   0.027246   0.441
##
## Correlation of Fixed Effects:
##             (Intr) Sl.LnF Sl.LnM Sexm   S.LF:S
## Sel.LineF   -0.292
## Sel.LineM   -0.295  0.496
## Sexm        -0.295  0.496  0.500
## Sel.LnF:Sxm  0.207 -0.709 -0.351 -0.703
## Sel.LnM:Sxm  0.209 -0.351 -0.709 -0.709  0.498
Anova(Model1)
## Analysis of Deviance Table (Type II Wald chisquare tests)
##
## Response: CHC_data$Peak_24
##               Chisq Df Pr(>Chisq)
## Sel.Line     0.2387  2     0.8875
## Sex          0.3443  1     0.5573
## Sel.Line:Sex 0.6052  2     0.7389
Model2 = lmer(CHC_data$Peak_24 ~ Sel.Line + Sex  + (1|Blocks) + (1|Time), data = CHC_data, REML = FALSE)
summary(Model2)
## Linear mixed model fit by maximum likelihood  ['lmerMod']
## Formula: CHC_data$Peak_24 ~ Sel.Line + Sex + (1 | Blocks) + (1 | Time)
##    Data: CHC_data
##
##      AIC      BIC   logLik deviance df.resid
##   -556.0   -528.9    285.0   -570.0      345
##
## Scaled residuals:
##     Min      1Q  Median      3Q     Max
## -3.0845 -0.3419 -0.1152  0.2190  9.3158
##
## Random effects:
##  Groups   Name        Variance Std.Dev.
##  Time     (Intercept) 0.002605 0.05104
##  Blocks   (Intercept) 0.001356 0.03682
##  Residual             0.011011 0.10493
## Number of obs: 352, groups:  Time, 6; Blocks, 3
##
## Fixed effects:
##              Estimate Std. Error t value
## (Intercept)  0.229097   0.031793   7.206
## Sel.LineF    0.004301   0.013754   0.313
## Sel.LineM   -0.002309   0.013635  -0.169
## Sexm        -0.006559   0.011186  -0.586
##
## Correlation of Fixed Effects:
##           (Intr) Sl.LnF Sl.LnM
## Sel.LineF -0.213
## Sel.LineM -0.215  0.498
## Sexm      -0.176 -0.004 -0.003
Anova(Model2)
## Analysis of Deviance Table (Type II Wald chisquare tests)
##
## Response: CHC_data$Peak_24
##           Chisq Df Pr(>Chisq)
## Sel.Line 0.2383  2     0.8877
## Sex      0.3437  1     0.5577
anova(Model1, Model2)
## Data: CHC_data
## Models:
## Model2: CHC_data$Peak_24 ~ Sel.Line + Sex + (1 | Blocks) + (1 | Time)
## Model1: CHC_data$Peak_24 ~ Sel.Line + Sex + Sel.Line * Sex + (1 | Blocks) +
## Model1:     (1 | Time)
##        Df     AIC     BIC logLik deviance  Chisq Chi Df Pr(>Chisq)
## Model2  7 -555.96 -528.92 284.98  -569.96
## Model1  9 -552.57 -517.79 285.28  -570.57 0.6047      2     0.7391
```

### Linear mixed effects analysis of Peak 25

```
Model1 = lmer(CHC_data$Peak_25 ~ Sel.Line + Sex + Sel.Line*Sex + (1|Blocks) + (1|Time), data = CHC_data, REML = FALSE)
summary(Model1)
## Linear mixed model fit by maximum likelihood  ['lmerMod']
## Formula: CHC_data$Peak_25 ~ Sel.Line + Sex + Sel.Line * Sex + (1 | Blocks) +
##     (1 | Time)
##    Data: CHC_data
##
##      AIC      BIC   logLik deviance df.resid
##    -32.4      2.3     25.2    -50.4      343
##
## Scaled residuals:
##     Min      1Q  Median      3Q     Max
## -2.8174 -0.4755 -0.1452  0.3341  8.3040
##
## Random effects:
##  Groups   Name        Variance  Std.Dev.
##  Time     (Intercept) 2.859e-02 1.691e-01
##  Blocks   (Intercept) 1.356e-09 3.683e-05
##  Residual             4.772e-02 2.185e-01
## Number of obs: 352, groups:  Time, 6; Blocks, 3
##
## Fixed effects:
##                Estimate Std. Error t value
## (Intercept)     0.81700    0.07466  10.942
## Sel.LineF       0.08775    0.04061   2.161
## Sel.LineM      -0.07266    0.04023  -1.806
## Sexm           -0.47901    0.04023 -11.906
## Sel.LineF:Sexm -0.08465    0.05729  -1.478
## Sel.LineM:Sexm  0.03867    0.05677   0.681
##
## Correlation of Fixed Effects:
##             (Intr) Sl.LnF Sl.LnM Sexm   S.LF:S
## Sel.LineF   -0.267
## Sel.LineM   -0.270  0.496
## Sexm        -0.270  0.496  0.500
## Sel.LnF:Sxm  0.189 -0.709 -0.351 -0.703
## Sel.LnM:Sxm  0.191 -0.351 -0.709 -0.709  0.498
Anova(Model1)
## Analysis of Deviance Table (Type II Wald chisquare tests)
##
## Response: CHC_data$Peak_25
##                Chisq Df Pr(>Chisq)
## Sel.Line      11.851  2   0.002671 **
## Sex          449.185  1  < 2.2e-16 ***
## Sel.Line:Sex   4.852  2   0.088388 .
## ---
## Signif. codes:  0 '***' 0.001 '**' 0.01 '*' 0.05 '.' 0.1 ' ' 1
Model2 = lmer(CHC_data$Peak_25 ~ Sel.Line + Sex  + (1|Blocks) + (1|Time), data = CHC_data, REML = FALSE)
summary(Model2)
## Linear mixed model fit by maximum likelihood  ['lmerMod']
## Formula: CHC_data$Peak_25 ~ Sel.Line + Sex + (1 | Blocks) + (1 | Time)
##    Data: CHC_data
##
##      AIC      BIC   logLik deviance df.resid
##    -31.6     -4.6     22.8    -45.6      345
##
## Scaled residuals:
##     Min      1Q  Median      3Q     Max
## -2.6374 -0.4766 -0.1655  0.3578  8.4103
##
## Random effects:
##  Groups   Name        Variance Std.Dev.
##  Time     (Intercept) 0.02837  0.1684
##  Blocks   (Intercept) 0.00000  0.0000
##  Residual             0.04840  0.2200
## Number of obs: 352, groups:  Time, 6; Blocks, 3
##
## Fixed effects:
##             Estimate Std. Error t value
## (Intercept)  0.82430    0.07263  11.349
## Sel.LineF    0.04509    0.02884   1.563
## Sel.LineM   -0.05310    0.02859  -1.858
## Sexm        -0.49359    0.02345 -21.046
##
## Correlation of Fixed Effects:
##           (Intr) Sl.LnF Sl.LnM
## Sel.LineF -0.195
## Sel.LineM -0.197  0.498
## Sexm      -0.161 -0.004 -0.003
## convergence code: 0
## boundary (singular) fit: see ?isSingular
Anova(Model2)
## Analysis of Deviance Table (Type II Wald chisquare tests)
##
## Response: CHC_data$Peak_25
##            Chisq Df Pr(>Chisq)
## Sel.Line  11.685  2   0.002901 **
## Sex      442.917  1  < 2.2e-16 ***
## ---
## Signif. codes:  0 '***' 0.001 '**' 0.01 '*' 0.05 '.' 0.1 ' ' 1
anova(Model1, Model2)
## Data: CHC_data
## Models:
## Model2: CHC_data$Peak_25 ~ Sel.Line + Sex + (1 | Blocks) + (1 | Time)
## Model1: CHC_data$Peak_25 ~ Sel.Line + Sex + Sel.Line * Sex + (1 | Blocks) +
## Model1:     (1 | Time)
##        Df     AIC     BIC logLik deviance  Chisq Chi Df Pr(>Chisq)
## Model2  7 -31.621 -4.5758 22.811  -45.621
## Model1  9 -32.439  2.3339 25.219  -50.439 4.8176      2    0.08992 .
## ---
## Signif. codes:  0 '***' 0.001 '**' 0.01 '*' 0.05 '.' 0.1 ' ' 1
```

### Linear mixed effects analysis of Peak 26

```
Model1 = lmer(CHC_data$Peak_26 ~ Sel.Line + Sex + Sel.Line*Sex + (1|Blocks) + (1|Time), data = CHC_data, REML = FALSE)
summary(Model1)
## Linear mixed model fit by maximum likelihood  ['lmerMod']
## Formula: CHC_data$Peak_26 ~ Sel.Line + Sex + Sel.Line * Sex + (1 | Blocks) +
##     (1 | Time)
##    Data: CHC_data
##
##      AIC      BIC   logLik deviance df.resid
##   2511.8   2546.6  -1246.9   2493.8      343
##
## Scaled residuals:
##     Min      1Q  Median      3Q     Max
## -1.4016 -0.4780 -0.1733  0.1387  5.9649
##
## Random effects:
##  Groups   Name        Variance Std.Dev.
##  Time     (Intercept)  2.088   1.445
##  Blocks   (Intercept)  1.027   1.013
##  Residual             68.389   8.270
## Number of obs: 352, groups:  Time, 6; Blocks, 3
##
## Fixed effects:
##                Estimate Std. Error t value
## (Intercept)     16.0495     1.3601  11.800
## Sel.LineF       -1.6299     1.5367  -1.061
## Sel.LineM       -1.1991     1.5229  -0.787
## Sexm            -9.6914     1.5229  -6.364
## Sel.LineF:Sexm   2.1797     2.1683   1.005
## Sel.LineM:Sexm   0.8936     2.1490   0.416
##
## Correlation of Fixed Effects:
##             (Intr) Sl.LnF Sl.LnM Sexm   S.LF:S
## Sel.LineF   -0.555
## Sel.LineM   -0.560  0.496
## Sexm        -0.560  0.496  0.500
## Sel.LnF:Sxm  0.394 -0.709 -0.351 -0.703
## Sel.LnM:Sxm  0.397 -0.351 -0.709 -0.709  0.498
Anova(Model1)
## Analysis of Deviance Table (Type II Wald chisquare tests)
##
## Response: CHC_data$Peak_26
##                Chisq Df Pr(>Chisq)
## Sel.Line      0.5184  2     0.7717
## Sex          96.8779  1     <2e-16 ***
## Sel.Line:Sex  1.0200  2     0.6005
## ---
## Signif. codes:  0 '***' 0.001 '**' 0.01 '*' 0.05 '.' 0.1 ' ' 1
Model2 = lmer(CHC_data$Peak_26 ~ Sel.Line + Sex  + (1|Blocks) + (1|Time), data = CHC_data, REML = FALSE)
summary(Model2)
## Linear mixed model fit by maximum likelihood  ['lmerMod']
## Formula: CHC_data$Peak_26 ~ Sel.Line + Sex + (1 | Blocks) + (1 | Time)
##    Data: CHC_data
##
##      AIC      BIC   logLik deviance df.resid
##   2508.9   2535.9  -1247.4   2494.9      345
##
## Scaled residuals:
##     Min      1Q  Median      3Q     Max
## -1.3351 -0.4792 -0.1762  0.1391  6.0169
##
## Random effects:
##  Groups   Name        Variance Std.Dev.
##  Time     (Intercept)  2.050   1.432
##  Blocks   (Intercept)  1.018   1.009
##  Residual             68.603   8.283
## Number of obs: 352, groups:  Time, 6; Blocks, 3
##
## Fixed effects:
##             Estimate Std. Error t value
## (Intercept)  15.5422     1.2073  12.874
## Sel.LineF    -0.5345     1.0856  -0.492
## Sel.LineM    -0.7528     1.0762  -0.700
## Sexm         -8.6774     0.8830  -9.827
##
## Correlation of Fixed Effects:
##           (Intr) Sl.LnF Sl.LnM
## Sel.LineF -0.443
## Sel.LineM -0.446  0.498
## Sexm      -0.366 -0.004 -0.003
Anova(Model2)
## Analysis of Deviance Table (Type II Wald chisquare tests)
##
## Response: CHC_data$Peak_26
##           Chisq Df Pr(>Chisq)
## Sel.Line  0.517  2     0.7722
## Sex      96.578  1     <2e-16 ***
## ---
## Signif. codes:  0 '***' 0.001 '**' 0.01 '*' 0.05 '.' 0.1 ' ' 1
anova(Model1, Model2)
## Data: CHC_data
## Models:
## Model2: CHC_data$Peak_26 ~ Sel.Line + Sex + (1 | Blocks) + (1 | Time)
## Model1: CHC_data$Peak_26 ~ Sel.Line + Sex + Sel.Line * Sex + (1 | Blocks) +
## Model1:     (1 | Time)
##        Df    AIC    BIC  logLik deviance  Chisq Chi Df Pr(>Chisq)
## Model2  7 2508.8 2535.9 -1247.4   2494.8
## Model1  9 2511.8 2546.6 -1246.9   2493.8 1.0181      2     0.6011
```

### Linear mixed effects analysis of Peak 27

```
Model1 = lmer(CHC_data$Peak_27 ~ Sel.Line + Sex + Sel.Line*Sex + (1|Blocks) + (1|Time), data = CHC_data, REML = FALSE)
summary(Model1)
## Linear mixed model fit by maximum likelihood  ['lmerMod']
## Formula: CHC_data$Peak_27 ~ Sel.Line + Sex + Sel.Line * Sex + (1 | Blocks) +
##     (1 | Time)
##    Data: CHC_data
##
##      AIC      BIC   logLik deviance df.resid
##   1467.6   1502.4   -724.8   1449.6      343
##
## Scaled residuals:
##     Min      1Q  Median      3Q     Max
## -2.2493 -0.1748 -0.0619  0.2905  8.4364
##
## Random effects:
##  Groups   Name        Variance Std.Dev.
##  Time     (Intercept) 0.1293   0.3596
##  Blocks   (Intercept) 0.0000   0.0000
##  Residual             3.5286   1.8784
## Number of obs: 352, groups:  Time, 6; Blocks, 3
##
## Fixed effects:
##                Estimate Std. Error t value
## (Intercept)      7.6704     0.2853  26.887
## Sel.LineF        0.5919     0.3491   1.696
## Sel.LineM       -1.1317     0.3459  -3.272
## Sexm            -7.5051     0.3459 -21.696
## Sel.LineF:Sexm  -0.5791     0.4925  -1.176
## Sel.LineM:Sexm   1.1482     0.4881   2.352
##
## Correlation of Fixed Effects:
##             (Intr) Sl.LnF Sl.LnM Sexm   S.LF:S
## Sel.LineF   -0.601
## Sel.LineM   -0.606  0.496
## Sexm        -0.606  0.496  0.500
## Sel.LnF:Sxm  0.426 -0.709 -0.351 -0.703
## Sel.LnM:Sxm  0.430 -0.351 -0.709 -0.709  0.498
## convergence code: 0
## boundary (singular) fit: see ?isSingular
Anova(Model1)
## Analysis of Deviance Table (Type II Wald chisquare tests)
##
## Response: CHC_data$Peak_27
##                 Chisq Df Pr(>Chisq)
## Sel.Line       12.444  2   0.001985 **
## Sex          1331.112  1  < 2.2e-16 ***
## Sel.Line:Sex   12.856  2   0.001615 **
## ---
## Signif. codes:  0 '***' 0.001 '**' 0.01 '*' 0.05 '.' 0.1 ' ' 1
Model2 = lmer(CHC_data$Peak_27 ~ Sel.Line + Sex  + (1|Blocks) + (1|Time), data = CHC_data, REML = FALSE)
summary(Model2)
## Linear mixed model fit by maximum likelihood  ['lmerMod']
## Formula: CHC_data$Peak_27 ~ Sel.Line + Sex + (1 | Blocks) + (1 | Time)
##    Data: CHC_data
##
##      AIC      BIC   logLik deviance df.resid
##   1476.3   1503.3   -731.1   1462.3      345
##
## Scaled residuals:
##     Min      1Q  Median      3Q     Max
## -2.1561 -0.3369 -0.0610  0.2545  8.4914
##
## Random effects:
##  Groups   Name        Variance Std.Dev.
##  Time     (Intercept) 0.1285   0.3585
##  Blocks   (Intercept) 0.0000   0.0000
##  Residual             3.6592   1.9129
## Number of obs: 352, groups:  Time, 6; Blocks, 3
##
## Fixed effects:
##             Estimate Std. Error t value
## (Intercept)   7.5710     0.2507  30.203
## Sel.LineF     0.2987     0.2507   1.191
## Sel.LineM    -0.5536     0.2485  -2.228
## Sexm         -7.3061     0.2039 -35.827
##
## Correlation of Fixed Effects:
##           (Intr) Sl.LnF Sl.LnM
## Sel.LineF -0.492
## Sel.LineM -0.496  0.498
## Sexm      -0.407 -0.004 -0.003
## convergence code: 0
## boundary (singular) fit: see ?isSingular
Anova(Model2)
## Analysis of Deviance Table (Type II Wald chisquare tests)
##
## Response: CHC_data$Peak_27
##             Chisq Df Pr(>Chisq)
## Sel.Line   11.999  2    0.00248 **
## Sex      1283.605  1    < 2e-16 ***
## ---
## Signif. codes:  0 '***' 0.001 '**' 0.01 '*' 0.05 '.' 0.1 ' ' 1
anova(Model1, Model2)
## Data: CHC_data
## Models:
## Model2: CHC_data$Peak_27 ~ Sel.Line + Sex + (1 | Blocks) + (1 | Time)
## Model1: CHC_data$Peak_27 ~ Sel.Line + Sex + Sel.Line * Sex + (1 | Blocks) +
## Model1:     (1 | Time)
##        Df    AIC    BIC  logLik deviance  Chisq Chi Df Pr(>Chisq)
## Model2  7 1476.3 1503.3 -731.14   1462.3
## Model1  9 1467.7 1502.4 -724.82   1449.7 12.625      2   0.001814 **
## ---
## Signif. codes:  0 '***' 0.001 '**' 0.01 '*' 0.05 '.' 0.1 ' ' 1
```

### Linear mixed effects analysis of Peak 28

```
Model1 = lmer(CHC_data$Peak_28 ~ Sel.Line + Sex + Sel.Line*Sex + (1|Blocks) + (1|Time), data = CHC_data, REML = FALSE)
summary(Model1)
## Linear mixed model fit by maximum likelihood  ['lmerMod']
## Formula: CHC_data$Peak_28 ~ Sel.Line + Sex + Sel.Line * Sex + (1 | Blocks) +
##     (1 | Time)
##    Data: CHC_data
##
##      AIC      BIC   logLik deviance df.resid
##   2521.3   2556.1  -1251.7   2503.3      343
##
## Scaled residuals:
##     Min      1Q  Median      3Q     Max
## -3.2751 -0.3628 -0.0268  0.4291  5.5489
##
## Random effects:
##  Groups   Name        Variance Std.Dev.
##  Time     (Intercept)  3.283   1.812
##  Blocks   (Intercept)  2.677   1.636
##  Residual             69.723   8.350
## Number of obs: 352, groups:  Time, 6; Blocks, 3
##
## Fixed effects:
##                Estimate Std. Error t value
## (Intercept)      24.897      1.619  15.376
## Sel.LineF         5.198      1.552   3.350
## Sel.LineM        -2.980      1.538  -1.938
## Sexm            -24.485      1.538 -15.923
## Sel.LineF:Sexm   -5.228      2.190  -2.388
## Sel.LineM:Sexm    2.975      2.170   1.371
##
## Correlation of Fixed Effects:
##             (Intr) Sl.LnF Sl.LnM Sexm   S.LF:S
## Sel.LineF   -0.471
## Sel.LineM   -0.475  0.496
## Sexm        -0.475  0.496  0.500
## Sel.LnF:Sxm  0.334 -0.709 -0.351 -0.703
## Sel.LnM:Sxm  0.337 -0.351 -0.709 -0.709  0.498
Anova(Model1)
## Analysis of Deviance Table (Type II Wald chisquare tests)
##
## Response: CHC_data$Peak_28
##                Chisq Df Pr(>Chisq)
## Sel.Line      13.979  2  0.0009216 ***
## Sex          800.573  1  < 2.2e-16 ***
## Sel.Line:Sex  14.411  2  0.0007424 ***
## ---
## Signif. codes:  0 '***' 0.001 '**' 0.01 '*' 0.05 '.' 0.1 ' ' 1
Model2 = lmer(CHC_data$Peak_28 ~ Sel.Line + Sex  + (1|Blocks) + (1|Time), data = CHC_data, REML = FALSE)
summary(Model2)
## Linear mixed model fit by maximum likelihood  ['lmerMod']
## Formula: CHC_data$Peak_28 ~ Sel.Line + Sex + (1 | Blocks) + (1 | Time)
##    Data: CHC_data
##
##      AIC      BIC   logLik deviance df.resid
##   2531.4   2558.5  -1258.7   2517.4      345
##
## Scaled residuals:
##     Min      1Q  Median      3Q     Max
## -2.9731 -0.3382  0.0167  0.4406  5.7111
##
## Random effects:
##  Groups   Name        Variance Std.Dev.
##  Time     (Intercept)  3.423   1.850
##  Blocks   (Intercept)  2.459   1.568
##  Residual             72.616   8.522
## Number of obs: 352, groups:  Time, 6; Blocks, 3
##
## Fixed effects:
##             Estimate Std. Error t value
## (Intercept)  25.2490     1.4873  16.976
## Sel.LineF     2.5630     1.1169   2.295
## Sel.LineM    -1.4767     1.1072  -1.334
## Sexm        -25.1865     0.9084 -27.725
##
## Correlation of Fixed Effects:
##           (Intr) Sl.LnF Sl.LnM
## Sel.LineF -0.370
## Sel.LineM -0.373  0.498
## Sexm      -0.305 -0.004 -0.003
Anova(Model2)
## Analysis of Deviance Table (Type II Wald chisquare tests)
##
## Response: CHC_data$Peak_28
##            Chisq Df Pr(>Chisq)
## Sel.Line  13.419  2   0.001219 **
## Sex      768.678  1  < 2.2e-16 ***
## ---
## Signif. codes:  0 '***' 0.001 '**' 0.01 '*' 0.05 '.' 0.1 ' ' 1
anova(Model1, Model2)
## Data: CHC_data
## Models:
## Model2: CHC_data$Peak_28 ~ Sel.Line + Sex + (1 | Blocks) + (1 | Time)
## Model1: CHC_data$Peak_28 ~ Sel.Line + Sex + Sel.Line * Sex + (1 | Blocks) +
## Model1:     (1 | Time)
##        Df    AIC    BIC  logLik deviance  Chisq Chi Df Pr(>Chisq)
## Model2  7 2531.4 2558.5 -1258.7   2517.4
## Model1  9 2521.3 2556.1 -1251.7   2503.3 14.118      2  0.0008597 ***
## ---
## Signif. codes:  0 '***' 0.001 '**' 0.01 '*' 0.05 '.' 0.1 ' ' 1
```

### Linear mixed effects analysis of Peak 29

```
Model1 = lmer(CHC_data$Peak_29 ~ Sel.Line + Sex + Sel.Line*Sex + (1|Blocks) + (1|Time), data = CHC_data, REML = FALSE)
summary(Model1)
## Linear mixed model fit by maximum likelihood  ['lmerMod']
## Formula: CHC_data$Peak_29 ~ Sel.Line + Sex + Sel.Line * Sex + (1 | Blocks) +
##     (1 | Time)
##    Data: CHC_data
##
##      AIC      BIC   logLik deviance df.resid
##    246.1    280.9   -114.1    228.1      343
##
## Scaled residuals:
##     Min      1Q  Median      3Q     Max
## -2.7960 -0.1775 -0.0410  0.2548  4.6920
##
## Random effects:
##  Groups   Name        Variance Std.Dev.
##  Time     (Intercept) 0.003036 0.0551
##  Blocks   (Intercept) 0.000000 0.0000
##  Residual             0.110118 0.3318
## Number of obs: 352, groups:  Time, 6; Blocks, 3
##
## Fixed effects:
##                Estimate Std. Error t value
## (Intercept)     0.79807    0.04872  16.382
## Sel.LineF       0.07490    0.06166   1.215
## Sel.LineM      -0.14215    0.06111  -2.326
## Sexm           -0.79527    0.06111 -13.014
## Sel.LineF:Sexm -0.07366    0.08700  -0.847
## Sel.LineM:Sexm  0.14368    0.08623   1.666
##
## Correlation of Fixed Effects:
##             (Intr) Sl.LnF Sl.LnM Sexm   S.LF:S
## Sel.LineF   -0.622
## Sel.LineM   -0.627  0.496
## Sexm        -0.627  0.496  0.500
## Sel.LnF:Sxm  0.441 -0.709 -0.351 -0.703
## Sel.LnM:Sxm  0.445 -0.351 -0.709 -0.709  0.498
## convergence code: 0
## boundary (singular) fit: see ?isSingular
Anova(Model1)
## Analysis of Deviance Table (Type II Wald chisquare tests)
##
## Response: CHC_data$Peak_29
##                 Chisq Df Pr(>Chisq)
## Sel.Line       6.3351  2    0.04211 *
## Sex          474.7003  1    < 2e-16 ***
## Sel.Line:Sex   6.5113  2    0.03855 *
## ---
## Signif. codes:  0 '***' 0.001 '**' 0.01 '*' 0.05 '.' 0.1 ' ' 1
Model2 = lmer(CHC_data$Peak_29 ~ Sel.Line + Sex  + (1|Blocks) + (1|Time), data = CHC_data, REML = FALSE)
summary(Model2)
## Linear mixed model fit by maximum likelihood  ['lmerMod']
## Formula: CHC_data$Peak_29 ~ Sel.Line + Sex + (1 | Blocks) + (1 | Time)
##    Data: CHC_data
##
##      AIC      BIC   logLik deviance df.resid
##    248.6    275.6   -117.3    234.6      345
##
## Scaled residuals:
##     Min      1Q  Median      3Q     Max
## -2.6222 -0.2311 -0.0002  0.2982  4.6859
##
## Random effects:
##  Groups   Name        Variance Std.Dev.
##  Time     (Intercept) 0.003032 0.05506
##  Blocks   (Intercept) 0.000000 0.00000
##  Residual             0.112178 0.33493
## Number of obs: 352, groups:  Time, 6; Blocks, 3
##
## Fixed effects:
##             Estimate Std. Error t value
## (Intercept)  0.78583    0.04213  18.652
## Sel.LineF    0.03762    0.04390   0.857
## Sel.LineM   -0.06981    0.04352  -1.604
## Sexm        -0.77076    0.03571 -21.587
##
## Correlation of Fixed Effects:
##           (Intr) Sl.LnF Sl.LnM
## Sel.LineF -0.513
## Sel.LineM -0.517  0.498
## Sexm      -0.424 -0.004 -0.003
## convergence code: 0
## boundary (singular) fit: see ?isSingular
Anova(Model2)
## Analysis of Deviance Table (Type II Wald chisquare tests)
##
## Response: CHC_data$Peak_29
##             Chisq Df Pr(>Chisq)
## Sel.Line   6.2184  2    0.04464 *
## Sex      465.9854  1    < 2e-16 ***
## ---
## Signif. codes:  0 '***' 0.001 '**' 0.01 '*' 0.05 '.' 0.1 ' ' 1
anova(Model1, Model2)
## Data: CHC_data
## Models:
## Model2: CHC_data$Peak_29 ~ Sel.Line + Sex + (1 | Blocks) + (1 | Time)
## Model1: CHC_data$Peak_29 ~ Sel.Line + Sex + Sel.Line * Sex + (1 | Blocks) +
## Model1:     (1 | Time)
##        Df    AIC    BIC  logLik deviance  Chisq Chi Df Pr(>Chisq)
## Model2  7 248.57 275.62 -117.29   234.57
## Model1  9 246.12 280.90 -114.06   228.12 6.4514      2    0.03973 *
## ---
## Signif. codes:  0 '***' 0.001 '**' 0.01 '*' 0.05 '.' 0.1 ' ' 1
```

### Linear mixed effects analysis of Peak 30

```
Model1 = lmer(CHC_data$Peak_30 ~ Sel.Line + Sex + Sel.Line*Sex + (1|Blocks) + (1|Time), data = CHC_data, REML = FALSE)
summary(Model1)
## Linear mixed model fit by maximum likelihood  ['lmerMod']
## Formula: CHC_data$Peak_30 ~ Sel.Line + Sex + Sel.Line * Sex + (1 | Blocks) +
##     (1 | Time)
##    Data: CHC_data
##
##      AIC      BIC   logLik deviance df.resid
##   1112.4   1147.1   -547.2   1094.4      343
##
## Scaled residuals:
##     Min      1Q  Median      3Q     Max
## -2.4292 -0.5055  0.0296  0.3710  6.6017
##
## Random effects:
##  Groups   Name        Variance Std.Dev.
##  Time     (Intercept) 0.3394   0.5826
##  Blocks   (Intercept) 0.0000   0.0000
##  Residual             1.2497   1.1179
## Number of obs: 352, groups:  Time, 6; Blocks, 3
##
## Fixed effects:
##                Estimate Std. Error t value
## (Intercept)      6.1579     0.2789  22.083
## Sel.LineF        0.1767     0.2078   0.850
## Sel.LineM       -0.7053     0.2059  -3.426
## Sexm            -5.0195     0.2059 -24.381
## Sel.LineF:Sexm  -0.2507     0.2932  -0.855
## Sel.LineM:Sexm   0.6206     0.2905   2.136
##
## Correlation of Fixed Effects:
##             (Intr) Sl.LnF Sl.LnM Sexm   S.LF:S
## Sel.LineF   -0.366
## Sel.LineM   -0.369  0.496
## Sexm        -0.369  0.496  0.500
## Sel.LnF:Sxm  0.260 -0.709 -0.351 -0.703
## Sel.LnM:Sxm  0.262 -0.351 -0.709 -0.709  0.498
## convergence code: 0
## boundary (singular) fit: see ?isSingular
Anova(Model1)
## Analysis of Deviance Table (Type II Wald chisquare tests)
##
## Response: CHC_data$Peak_30
##                  Chisq Df Pr(>Chisq)
## Sel.Line       11.0917  2   0.003904 **
## Sex          1684.7808  1  < 2.2e-16 ***
## Sel.Line:Sex    9.4586  2   0.008833 **
## ---
## Signif. codes:  0 '***' 0.001 '**' 0.01 '*' 0.05 '.' 0.1 ' ' 1
Model2 = lmer(CHC_data$Peak_30 ~ Sel.Line + Sex  + (1|Blocks) + (1|Time), data = CHC_data, REML = FALSE)
summary(Model2)
## Linear mixed model fit by maximum likelihood  ['lmerMod']
## Formula: CHC_data$Peak_30 ~ Sel.Line + Sex + (1 | Blocks) + (1 | Time)
##    Data: CHC_data
##
##      AIC      BIC   logLik deviance df.resid
##   1117.7   1144.7   -551.8   1103.7      345
##
## Scaled residuals:
##     Min      1Q  Median      3Q     Max
## -2.6118 -0.5004 -0.0254  0.4271  6.5706
##
## Random effects:
##  Groups   Name        Variance Std.Dev.
##  Time     (Intercept) 0.3424   0.5852
##  Blocks   (Intercept) 0.0000   0.0000
##  Residual             1.2836   1.1330
## Number of obs: 352, groups:  Time, 6; Blocks, 3
##
## Fixed effects:
##             Estimate Std. Error t value
## (Intercept)  6.09397    0.26758  22.775
## Sel.LineF    0.04956    0.14851   0.334
## Sel.LineM   -0.39290    0.14722  -2.669
## Sexm        -4.89162    0.12078 -40.500
##
## Correlation of Fixed Effects:
##           (Intr) Sl.LnF Sl.LnM
## Sel.LineF -0.273
## Sel.LineM -0.276  0.498
## Sexm      -0.226 -0.004 -0.003
## convergence code: 0
## boundary (singular) fit: see ?isSingular
Anova(Model2)
## Analysis of Deviance Table (Type II Wald chisquare tests)
##
## Response: CHC_data$Peak_30
##             Chisq Df Pr(>Chisq)
## Sel.Line   10.799  2   0.004519 **
## Sex      1640.230  1  < 2.2e-16 ***
## ---
## Signif. codes:  0 '***' 0.001 '**' 0.01 '*' 0.05 '.' 0.1 ' ' 1
anova(Model1, Model2)
## Data: CHC_data
## Models:
## Model2: CHC_data$Peak_30 ~ Sel.Line + Sex + (1 | Blocks) + (1 | Time)
## Model1: CHC_data$Peak_30 ~ Sel.Line + Sex + Sel.Line * Sex + (1 | Blocks) +
## Model1:     (1 | Time)
##        Df    AIC    BIC  logLik deviance  Chisq Chi Df Pr(>Chisq)
## Model2  7 1117.7 1144.7 -551.85   1103.7
## Model1  9 1112.4 1147.1 -547.18   1094.4 9.3329      2   0.009405 **
## ---
## Signif. codes:  0 '***' 0.001 '**' 0.01 '*' 0.05 '.' 0.1 ' ' 1
```

### Linear mixed effects analysis of Peak 31

```
Model1 = lmer(CHC_data$Peak_31 ~ Sel.Line + Sex + Sel.Line*Sex + (1|Blocks) + (1|Time), data = CHC_data, REML = FALSE)
summary(Model1)
## Linear mixed model fit by maximum likelihood  ['lmerMod']
## Formula: CHC_data$Peak_31 ~ Sel.Line + Sex + Sel.Line * Sex + (1 | Blocks) +
##     (1 | Time)
##    Data: CHC_data
##
##      AIC      BIC   logLik deviance df.resid
##    352.7    387.5   -167.4    334.7      343
##
## Scaled residuals:
##     Min      1Q  Median      3Q     Max
## -1.5326 -0.5635 -0.2069  0.2308  8.5304
##
## Random effects:
##  Groups   Name        Variance Std.Dev.
##  Time     (Intercept) 0.028653 0.16927
##  Blocks   (Intercept) 0.001178 0.03433
##  Residual             0.145041 0.38084
## Number of obs: 352, groups:  Time, 6; Blocks, 3
##
## Fixed effects:
##                Estimate Std. Error t value
## (Intercept)     0.44359    0.08734   5.079
## Sel.LineF      -0.04052    0.07079  -0.573
## Sel.LineM      -0.04626    0.07014  -0.659
## Sexm           -0.11198    0.07014  -1.597
## Sel.LineF:Sexm  0.10864    0.09987   1.088
## Sel.LineM:Sexm  0.03631    0.09897   0.367
##
## Correlation of Fixed Effects:
##             (Intr) Sl.LnF Sl.LnM Sexm   S.LF:S
## Sel.LineF   -0.398
## Sel.LineM   -0.402  0.496
## Sexm        -0.402  0.496  0.500
## Sel.LnF:Sxm  0.282 -0.709 -0.351 -0.703
## Sel.LnM:Sxm  0.285 -0.351 -0.709 -0.709  0.498
Anova(Model1)
## Analysis of Deviance Table (Type II Wald chisquare tests)
##
## Response: CHC_data$Peak_31
##               Chisq Df Pr(>Chisq)
## Sel.Line     0.7485  2     0.6878
## Sex          2.5017  1     0.1137
## Sel.Line:Sex 1.2239  2     0.5423
Model2 = lmer(CHC_data$Peak_31 ~ Sel.Line + Sex  + (1|Blocks) + (1|Time), data = CHC_data, REML = FALSE)
summary(Model2)
## Linear mixed model fit by maximum likelihood  ['lmerMod']
## Formula: CHC_data$Peak_31 ~ Sel.Line + Sex + (1 | Blocks) + (1 | Time)
##    Data: CHC_data
##
##      AIC      BIC   logLik deviance df.resid
##      350      377     -168      336      345
##
## Scaled residuals:
##     Min      1Q  Median      3Q     Max
## -1.5313 -0.5378 -0.2088  0.2300  8.5972
##
## Random effects:
##  Groups   Name        Variance  Std.Dev.
##  Time     (Intercept) 2.972e-02 0.172398
##  Blocks   (Intercept) 3.955e-05 0.006289
##  Residual             1.456e-01 0.381522
## Number of obs: 352, groups:  Time, 6; Blocks, 3
##
## Fixed effects:
##             Estimate Std. Error t value
## (Intercept)  0.41970    0.08133   5.161
## Sel.LineF    0.01408    0.05001   0.282
## Sel.LineM   -0.02813    0.04957  -0.568
## Sexm        -0.06421    0.04067  -1.579
##
## Correlation of Fixed Effects:
##           (Intr) Sl.LnF Sl.LnM
## Sel.LineF -0.303
## Sel.LineM -0.305  0.498
## Sexm      -0.250 -0.004 -0.003
## convergence code: 0
## unable to evaluate scaled gradient
## Model failed to converge: degenerate  Hessian with 1 negative eigenvalues
Anova(Model2)
## Analysis of Deviance Table (Type II Wald chisquare tests)
##
## Response: CHC_data$Peak_31
##           Chisq Df Pr(>Chisq)
## Sel.Line 0.7453  2     0.6889
## Sex      2.4926  1     0.1144
anova(Model1, Model2)
## Data: CHC_data
## Models:
## Model2: CHC_data$Peak_31 ~ Sel.Line + Sex + (1 | Blocks) + (1 | Time)
## Model1: CHC_data$Peak_31 ~ Sel.Line + Sex + Sel.Line * Sex + (1 | Blocks) +
## Model1:     (1 | Time)
##        Df    AIC   BIC  logLik deviance  Chisq Chi Df Pr(>Chisq)
## Model2  7 349.95 377.0 -167.98   335.95
## Model1  9 352.73 387.5 -167.36   334.73 1.2269      2     0.5415
```

### Linear mixed effects analysis of Peak 32

```
Model1 = lmer(CHC_data$Peak_32 ~ Sel.Line + Sex + Sel.Line*Sex + (1|Blocks) + (1|Time), data = CHC_data, REML = FALSE)
summary(Model1)
## Linear mixed model fit by maximum likelihood  ['lmerMod']
## Formula: CHC_data$Peak_32 ~ Sel.Line + Sex + Sel.Line * Sex + (1 | Blocks) +
##     (1 | Time)
##    Data: CHC_data
##
##      AIC      BIC   logLik deviance df.resid
##   2122.2   2157.0  -1052.1   2104.2      343
##
## Scaled residuals:
##     Min      1Q  Median      3Q     Max
## -1.4726 -0.5594 -0.2182  0.2025  5.6067
##
## Random effects:
##  Groups   Name        Variance Std.Dev.
##  Time     (Intercept)  2.0829  1.443
##  Blocks   (Intercept)  0.3782  0.615
##  Residual             22.3307  4.726
## Number of obs: 352, groups:  Time, 6; Blocks, 3
##
## Fixed effects:
##                Estimate Std. Error t value
## (Intercept)     8.89065    0.92303   9.632
## Sel.LineF      -0.88734    0.87827  -1.010
## Sel.LineM      -0.01729    0.87026  -0.020
## Sexm           -4.13284    0.87026  -4.749
## Sel.LineF:Sexm  1.28764    1.23919   1.039
## Sel.LineM:Sexm  0.30464    1.22804   0.248
##
## Correlation of Fixed Effects:
##             (Intr) Sl.LnF Sl.LnM Sexm   S.LF:S
## Sel.LineF   -0.468
## Sel.LineM   -0.472  0.496
## Sexm        -0.472  0.496  0.500
## Sel.LnF:Sxm  0.331 -0.709 -0.351 -0.703
## Sel.LnM:Sxm  0.334 -0.351 -0.709 -0.709  0.498
Anova(Model1)
## Analysis of Deviance Table (Type II Wald chisquare tests)
##
## Response: CHC_data$Peak_32
##                Chisq Df Pr(>Chisq)
## Sel.Line      0.3750  2     0.8290
## Sex          51.3294  1   7.81e-13 ***
## Sel.Line:Sex  1.1761  2     0.5554
## ---
## Signif. codes:  0 '***' 0.001 '**' 0.01 '*' 0.05 '.' 0.1 ' ' 1
Model2 = lmer(CHC_data$Peak_32 ~ Sel.Line + Sex  + (1|Blocks) + (1|Time), data = CHC_data, REML = FALSE)
summary(Model2)
## Linear mixed model fit by maximum likelihood  ['lmerMod']
## Formula: CHC_data$Peak_32 ~ Sel.Line + Sex + (1 | Blocks) + (1 | Time)
##    Data: CHC_data
##
##      AIC      BIC   logLik deviance df.resid
##   2119.4   2146.5  -1052.7   2105.4      345
##
## Scaled residuals:
##     Min      1Q  Median      3Q     Max
## -1.4118 -0.5614 -0.2446  0.1951  5.6523
##
## Random effects:
##  Groups   Name        Variance Std.Dev.
##  Time     (Intercept)  2.0605  1.4355
##  Blocks   (Intercept)  0.3762  0.6133
##  Residual             22.4098  4.7339
## Number of obs: 352, groups:  Time, 6; Blocks, 3
##
## Fixed effects:
##             Estimate Std. Error t value
## (Intercept)   8.6287     0.8500  10.152
## Sel.LineF    -0.2399     0.6205  -0.387
## Sel.LineM     0.1342     0.6151   0.218
## Sexm         -3.6093     0.5047  -7.152
##
## Correlation of Fixed Effects:
##           (Intr) Sl.LnF Sl.LnM
## Sel.LineF -0.359
## Sel.LineM -0.362  0.498
## Sexm      -0.297 -0.004 -0.003
Anova(Model2)
## Analysis of Deviance Table (Type II Wald chisquare tests)
##
## Response: CHC_data$Peak_32
##            Chisq Df Pr(>Chisq)
## Sel.Line  0.3736  2     0.8296
## Sex      51.1488  1  8.562e-13 ***
## ---
## Signif. codes:  0 '***' 0.001 '**' 0.01 '*' 0.05 '.' 0.1 ' ' 1
anova(Model1, Model2)
## Data: CHC_data
## Models:
## Model2: CHC_data$Peak_32 ~ Sel.Line + Sex + (1 | Blocks) + (1 | Time)
## Model1: CHC_data$Peak_32 ~ Sel.Line + Sex + Sel.Line * Sex + (1 | Blocks) +
## Model1:     (1 | Time)
##        Df    AIC    BIC  logLik deviance  Chisq Chi Df Pr(>Chisq)
## Model2  7 2119.4 2146.5 -1052.7   2105.4
## Model1  9 2122.2 2157.0 -1052.1   2104.2 1.1737      2     0.5561
```

### Linear mixed effects analysis of Peak 33

```
Model1 = lmer(CHC_data$Peak_33 ~ Sel.Line + Sex + Sel.Line*Sex + (1|Blocks) + (1|Time), data = CHC_data, REML = FALSE)
summary(Model1)
## Linear mixed model fit by maximum likelihood  ['lmerMod']
## Formula: CHC_data$Peak_33 ~ Sel.Line + Sex + Sel.Line * Sex + (1 | Blocks) +
##     (1 | Time)
##    Data: CHC_data
##
##      AIC      BIC   logLik deviance df.resid
##   1263.4   1298.2   -622.7   1245.4      343
##
## Scaled residuals:
##     Min      1Q  Median      3Q     Max
## -0.6817 -0.1054 -0.0365  0.0293 17.2365
##
## Random effects:
##  Groups   Name        Variance  Std.Dev.
##  Time     (Intercept) 0.0000453 0.006731
##  Blocks   (Intercept) 0.0000000 0.000000
##  Residual             2.0140927 1.419187
## Number of obs: 352, groups:  Time, 6; Blocks, 3
##
## Fixed effects:
##                 Estimate Std. Error t value
## (Intercept)     0.568808   0.184783   3.078
## Sel.LineF       0.638919   0.263576   2.424
## Sel.LineM      -0.006944   0.261294  -0.027
## Sexm           -0.420325   0.261294  -1.609
## Sel.LineF:Sexm -0.665506   0.371935  -1.789
## Sel.LineM:Sexm  0.002128   0.368754   0.006
##
## Correlation of Fixed Effects:
##             (Intr) Sl.LnF Sl.LnM Sexm   S.LF:S
## Sel.LineF   -0.701
## Sel.LineM   -0.707  0.496
## Sexm        -0.707  0.496  0.500
## Sel.LnF:Sxm  0.497 -0.709 -0.351 -0.703
## Sel.LnM:Sxm  0.501 -0.351 -0.709 -0.709  0.498
## convergence code: 0
## boundary (singular) fit: see ?isSingular
Anova(Model1)
## Analysis of Deviance Table (Type II Wald chisquare tests)
##
## Response: CHC_data$Peak_33
##                Chisq Df Pr(>Chisq)
## Sel.Line      3.6170  2     0.1639
## Sex          17.7294  1  2.547e-05 ***
## Sel.Line:Sex  4.2701  2     0.1182
## ---
## Signif. codes:  0 '***' 0.001 '**' 0.01 '*' 0.05 '.' 0.1 ' ' 1
Model2 = lmer(CHC_data$Peak_33 ~ Sel.Line + Sex  + (1|Blocks) + (1|Time), data = CHC_data, REML = FALSE)
summary(Model2)
## Linear mixed model fit by maximum likelihood  ['lmerMod']
## Formula: CHC_data$Peak_33 ~ Sel.Line + Sex + (1 | Blocks) + (1 | Time)
##    Data: CHC_data
##
##      AIC      BIC   logLik deviance df.resid
##   1263.6   1290.7   -624.8   1249.6      345
##
## Scaled residuals:
##     Min      1Q  Median      3Q     Max
## -0.5192 -0.1836 -0.0768  0.0524 17.2916
##
## Random effects:
##  Groups   Name        Variance Std.Dev.
##  Time     (Intercept) 0.000    0.000
##  Blocks   (Intercept) 0.000    0.000
##  Residual             2.039    1.428
## Number of obs: 352, groups:  Time, 6; Blocks, 3
##
## Fixed effects:
##              Estimate Std. Error t value
## (Intercept)  0.677157   0.151881   4.458
## Sel.LineF    0.304213   0.187091   1.626
## Sel.LineM   -0.004959   0.185492  -0.027
## Sexm        -0.637026   0.152206  -4.185
##
## Correlation of Fixed Effects:
##           (Intr) Sl.LnF Sl.LnM
## Sel.LineF -0.606
## Sel.LineM -0.611  0.498
## Sexm      -0.501 -0.004 -0.003
## convergence code: 0
## boundary (singular) fit: see ?isSingular
Anova(Model2)
## Analysis of Deviance Table (Type II Wald chisquare tests)
##
## Response: CHC_data$Peak_33
##            Chisq Df Pr(>Chisq)
## Sel.Line  3.5735  2     0.1675
## Sex      17.5166  1  2.848e-05 ***
## ---
## Signif. codes:  0 '***' 0.001 '**' 0.01 '*' 0.05 '.' 0.1 ' ' 1
anova(Model1, Model2)
## Data: CHC_data
## Models:
## Model2: CHC_data$Peak_33 ~ Sel.Line + Sex + (1 | Blocks) + (1 | Time)
## Model1: CHC_data$Peak_33 ~ Sel.Line + Sex + Sel.Line * Sex + (1 | Blocks) +
## Model1:     (1 | Time)
##        Df    AIC    BIC  logLik deviance  Chisq Chi Df Pr(>Chisq)
## Model2  7 1263.6 1290.7 -624.82   1249.6
## Model1  9 1263.4 1298.2 -622.70   1245.4 4.2441      2     0.1198
```

### Linear mixed effects analysis of Peak 34

```
Model1 = lmer(CHC_data$Peak_34 ~ Sel.Line + Sex + Sel.Line*Sex + (1|Blocks) + (1|Time), data = CHC_data, REML = FALSE)
summary(Model1)
## Linear mixed model fit by maximum likelihood  ['lmerMod']
## Formula: CHC_data$Peak_34 ~ Sel.Line + Sex + Sel.Line * Sex + (1 | Blocks) +
##     (1 | Time)
##    Data: CHC_data
##
##      AIC      BIC   logLik deviance df.resid
##    690.4    725.1   -336.2    672.4      343
##
## Scaled residuals:
##     Min      1Q  Median      3Q     Max
## -2.5745 -0.3351 -0.0038  0.2523 10.7451
##
## Random effects:
##  Groups   Name        Variance Std.Dev.
##  Time     (Intercept) 0.044286 0.21044
##  Blocks   (Intercept) 0.001872 0.04327
##  Residual             0.381587 0.61773
## Number of obs: 352, groups:  Time, 6; Blocks, 3
##
## Fixed effects:
##                Estimate Std. Error t value
## (Intercept)     1.49304    0.12032  12.409
## Sel.LineF       0.27075    0.11481   2.358
## Sel.LineM      -0.08034    0.11376  -0.706
## Sexm           -1.31700    0.11376 -11.577
## Sel.LineF:Sexm -0.19390    0.16199  -1.197
## Sel.LineM:Sexm  0.14979    0.16053   0.933
##
## Correlation of Fixed Effects:
##             (Intr) Sl.LnF Sl.LnM Sexm   S.LF:S
## Sel.LineF   -0.469
## Sel.LineM   -0.473  0.496
## Sexm        -0.473  0.496  0.500
## Sel.LnF:Sxm  0.332 -0.709 -0.351 -0.703
## Sel.LnM:Sxm  0.335 -0.351 -0.709 -0.709  0.498
Anova(Model1)
## Analysis of Deviance Table (Type II Wald chisquare tests)
##
## Response: CHC_data$Peak_34
##                 Chisq Df Pr(>Chisq)
## Sel.Line       6.2399  2    0.04416 *
## Sex          407.7042  1    < 2e-16 ***
## Sel.Line:Sex   4.5411  2    0.10326
## ---
## Signif. codes:  0 '***' 0.001 '**' 0.01 '*' 0.05 '.' 0.1 ' ' 1
Model2 = lmer(CHC_data$Peak_34 ~ Sel.Line + Sex  + (1|Blocks) + (1|Time), data = CHC_data, REML = FALSE)
summary(Model2)
## Linear mixed model fit by maximum likelihood  ['lmerMod']
## Formula: CHC_data$Peak_34 ~ Sel.Line + Sex + (1 | Blocks) + (1 | Time)
##    Data: CHC_data
##
##      AIC      BIC   logLik deviance df.resid
##    690.9    717.9   -338.4    676.9      345
##
## Scaled residuals:
##     Min      1Q  Median      3Q     Max
## -2.4082 -0.3743 -0.0076  0.2885 10.5490
##
## Random effects:
##  Groups   Name        Variance Std.Dev.
##  Time     (Intercept) 0.044956 0.21203
##  Blocks   (Intercept) 0.001148 0.03388
##  Residual             0.386588 0.62176
## Number of obs: 352, groups:  Time, 6; Blocks, 3
##
## Fixed effects:
##              Estimate Std. Error t value
## (Intercept)  1.499419   0.110685  13.547
## Sel.LineF    0.172945   0.081498   2.122
## Sel.LineM   -0.004759   0.080789  -0.059
## Sexm        -1.329692   0.066284 -20.061
##
## Correlation of Fixed Effects:
##           (Intr) Sl.LnF Sl.LnM
## Sel.LineF -0.362
## Sel.LineM -0.366  0.498
## Sexm      -0.299 -0.004 -0.003
Anova(Model2)
## Analysis of Deviance Table (Type II Wald chisquare tests)
##
## Response: CHC_data$Peak_34
##             Chisq Df Pr(>Chisq)
## Sel.Line   6.1581  2      0.046 *
## Sex      402.4302  1     <2e-16 ***
## ---
## Signif. codes:  0 '***' 0.001 '**' 0.01 '*' 0.05 '.' 0.1 ' ' 1
anova(Model1, Model2)
## Data: CHC_data
## Models:
## Model2: CHC_data$Peak_34 ~ Sel.Line + Sex + (1 | Blocks) + (1 | Time)
## Model1: CHC_data$Peak_34 ~ Sel.Line + Sex + Sel.Line * Sex + (1 | Blocks) +
## Model1:     (1 | Time)
##        Df    AIC    BIC  logLik deviance  Chisq Chi Df Pr(>Chisq)
## Model2  7 690.87 717.91 -338.43   676.87
## Model1  9 690.35 725.13 -336.18   672.35 4.5111      2     0.1048
```

### Linear mixed effects analysis of Peak 35

```
Model1 = lmer(CHC_data$Peak_35 ~ Sel.Line + Sex + Sel.Line*Sex + (1|Blocks) + (1|Time), data = CHC_data, REML = FALSE)
summary(Model1)
## Linear mixed model fit by maximum likelihood  ['lmerMod']
## Formula: CHC_data$Peak_35 ~ Sel.Line + Sex + Sel.Line * Sex + (1 | Blocks) +
##     (1 | Time)
##    Data: CHC_data
##
##      AIC      BIC   logLik deviance df.resid
##    -42.5     -7.7     30.2    -60.5      343
##
## Scaled residuals:
##     Min      1Q  Median      3Q     Max
## -2.0271 -0.4753 -0.0439  0.2838  7.8121
##
## Random effects:
##  Groups   Name        Variance Std.Dev.
##  Time     (Intercept) 0.01045  0.1022
##  Blocks   (Intercept) 0.00000  0.0000
##  Residual             0.04714  0.2171
## Number of obs: 352, groups:  Time, 6; Blocks, 3
##
## Fixed effects:
##                Estimate Std. Error t value
## (Intercept)     0.93382    0.05042  18.523
## Sel.LineF       0.06206    0.04035   1.538
## Sel.LineM      -0.07994    0.03998  -1.999
## Sexm           -0.56597    0.03998 -14.155
## Sel.LineF:Sexm -0.06957    0.05694  -1.222
## Sel.LineM:Sexm  0.09209    0.05642   1.632
##
## Correlation of Fixed Effects:
##             (Intr) Sl.LnF Sl.LnM Sexm   S.LF:S
## Sel.LineF   -0.393
## Sel.LineM   -0.397  0.496
## Sexm        -0.397  0.496  0.500
## Sel.LnF:Sxm  0.279 -0.709 -0.351 -0.703
## Sel.LnM:Sxm  0.281 -0.351 -0.709 -0.709  0.498
## convergence code: 0
## boundary (singular) fit: see ?isSingular
Anova(Model1)
## Analysis of Deviance Table (Type II Wald chisquare tests)
##
## Response: CHC_data$Peak_35
##                 Chisq Df Pr(>Chisq)
## Sel.Line       4.5596  2    0.10231
## Sex          580.3199  1    < 2e-16 ***
## Sel.Line:Sex   8.1675  2    0.01684 *
## ---
## Signif. codes:  0 '***' 0.001 '**' 0.01 '*' 0.05 '.' 0.1 ' ' 1
Model2 = lmer(CHC_data$Peak_35 ~ Sel.Line + Sex  + (1|Blocks) + (1|Time), data = CHC_data, REML = FALSE)
summary(Model2)
## Linear mixed model fit by maximum likelihood  ['lmerMod']
## Formula: CHC_data$Peak_35 ~ Sel.Line + Sex + (1 | Blocks) + (1 | Time)
##    Data: CHC_data
##
##      AIC      BIC   logLik deviance df.resid
##    -38.4    -11.4     26.2    -52.4      345
##
## Scaled residuals:
##     Min      1Q  Median      3Q     Max
## -2.1904 -0.5309 -0.0391  0.3035  7.7411
##
## Random effects:
##  Groups   Name        Variance Std.Dev.
##  Time     (Intercept) 0.01051  0.1025
##  Blocks   (Intercept) 0.00000  0.0000
##  Residual             0.04824  0.2196
## Number of obs: 352, groups:  Time, 6; Blocks, 3
##
## Fixed effects:
##             Estimate Std. Error t value
## (Intercept)  0.92963    0.04793  19.395
## Sel.LineF    0.02692    0.02879   0.935
## Sel.LineM   -0.03355    0.02854  -1.175
## Sexm        -0.55756    0.02342 -23.812
##
## Correlation of Fixed Effects:
##           (Intr) Sl.LnF Sl.LnM
## Sel.LineF -0.296
## Sel.LineM -0.298  0.498
## Sexm      -0.244 -0.004 -0.003
## convergence code: 0
## boundary (singular) fit: see ?isSingular
Anova(Model2)
## Analysis of Deviance Table (Type II Wald chisquare tests)
##
## Response: CHC_data$Peak_35
##             Chisq Df Pr(>Chisq)
## Sel.Line   4.4551  2     0.1078
## Sex      567.0012  1     <2e-16 ***
## ---
## Signif. codes:  0 '***' 0.001 '**' 0.01 '*' 0.05 '.' 0.1 ' ' 1
anova(Model1, Model2)
## Data: CHC_data
## Models:
## Model2: CHC_data$Peak_35 ~ Sel.Line + Sex + (1 | Blocks) + (1 | Time)
## Model1: CHC_data$Peak_35 ~ Sel.Line + Sex + Sel.Line * Sex + (1 | Blocks) +
## Model1:     (1 | Time)
##        Df     AIC      BIC logLik deviance  Chisq Chi Df Pr(>Chisq)
## Model2  7 -38.417 -11.3718 26.209  -52.417
## Model1  9 -42.491  -7.7179 30.245  -60.491 8.0733      2    0.01766 *
## ---
## Signif. codes:  0 '***' 0.001 '**' 0.01 '*' 0.05 '.' 0.1 ' ' 1
```

### Linear mixed effects analysis of Peak 36

```
Model1 = lmer(CHC_data$Peak_36 ~ Sel.Line + Sex + Sel.Line*Sex + (1|Blocks) + (1|Time), data = CHC_data, REML = FALSE)
summary(Model1)
## Linear mixed model fit by maximum likelihood  ['lmerMod']
## Formula: CHC_data$Peak_36 ~ Sel.Line + Sex + Sel.Line * Sex + (1 | Blocks) +
##     (1 | Time)
##    Data: CHC_data
##
##      AIC      BIC   logLik deviance df.resid
##   1783.6   1818.4   -882.8   1765.6      343
##
## Scaled residuals:
##     Min      1Q  Median      3Q     Max
## -4.7526 -0.1736  0.0133  0.2656  6.1752
##
## Random effects:
##  Groups   Name        Variance Std.Dev.
##  Time     (Intercept) 25.206   5.021
##  Blocks   (Intercept)  0.000   0.000
##  Residual              8.078   2.842
## Number of obs: 352, groups:  Time, 6; Blocks, 3
##
## Fixed effects:
##                Estimate Std. Error t value
## (Intercept)      4.8963     2.0828   2.351
## Sel.LineF       -0.7881     0.5283  -1.492
## Sel.LineM       -0.8623     0.5234  -1.647
## Sexm             0.3586     0.5234   0.685
## Sel.LineF:Sexm   0.1276     0.7454   0.171
## Sel.LineM:Sexm   0.5650     0.7386   0.765
##
## Correlation of Fixed Effects:
##             (Intr) Sl.LnF Sl.LnM Sexm   S.LF:S
## Sel.LineF   -0.125
## Sel.LineM   -0.126  0.496
## Sexm        -0.126  0.496  0.500
## Sel.LnF:Sxm  0.088 -0.709 -0.351 -0.703
## Sel.LnM:Sxm  0.089 -0.351 -0.709 -0.709  0.498
## convergence code: 0
## boundary (singular) fit: see ?isSingular
Anova(Model1)
## Analysis of Deviance Table (Type II Wald chisquare tests)
##
## Response: CHC_data$Peak_36
##               Chisq Df Pr(>Chisq)
## Sel.Line     4.2583  2    0.11894
## Sex          3.8085  1    0.05099 .
## Sel.Line:Sex 0.6436  2    0.72484
## ---
## Signif. codes:  0 '***' 0.001 '**' 0.01 '*' 0.05 '.' 0.1 ' ' 1
Model2 = lmer(CHC_data$Peak_36 ~ Sel.Line + Sex  + (1|Blocks) + (1|Time), data = CHC_data, REML = FALSE)
summary(Model2)
## Linear mixed model fit by maximum likelihood  ['lmerMod']
## Formula: CHC_data$Peak_36 ~ Sel.Line + Sex + (1 | Blocks) + (1 | Time)
##    Data: CHC_data
##
##      AIC      BIC   logLik deviance df.resid
##   1780.2   1807.3   -883.1   1766.2      345
##
## Scaled residuals:
##     Min      1Q  Median      3Q     Max
## -4.7072 -0.1782  0.0015  0.2562  6.1509
##
## Random effects:
##  Groups   Name        Variance Std.Dev.
##  Time     (Intercept) 25.210   5.021
##  Blocks   (Intercept)  0.000   0.000
##  Residual              8.093   2.845
## Number of obs: 352, groups:  Time, 6; Blocks, 3
##
## Fixed effects:
##             Estimate Std. Error t value
## (Intercept)   4.7799     2.0720   2.307
## Sel.LineF    -0.7247     0.3729  -1.943
## Sel.LineM    -0.5784     0.3696  -1.565
## Sexm          0.5913     0.3033   1.950
##
## Correlation of Fixed Effects:
##           (Intr) Sl.LnF Sl.LnM
## Sel.LineF -0.089
## Sel.LineM -0.089  0.498
## Sexm      -0.073 -0.004 -0.003
## convergence code: 0
## boundary (singular) fit: see ?isSingular
Anova(Model2)
## Analysis of Deviance Table (Type II Wald chisquare tests)
##
## Response: CHC_data$Peak_36
##           Chisq Df Pr(>Chisq)
## Sel.Line 4.2505  2    0.11941
## Sex      3.8014  1    0.05121 .
## ---
## Signif. codes:  0 '***' 0.001 '**' 0.01 '*' 0.05 '.' 0.1 ' ' 1
anova(Model1, Model2)
## Data: CHC_data
## Models:
## Model2: CHC_data$Peak_36 ~ Sel.Line + Sex + (1 | Blocks) + (1 | Time)
## Model1: CHC_data$Peak_36 ~ Sel.Line + Sex + Sel.Line * Sex + (1 | Blocks) +
## Model1:     (1 | Time)
##        Df    AIC    BIC  logLik deviance Chisq Chi Df Pr(>Chisq)
## Model2  7 1780.2 1807.3 -883.12   1766.2
## Model1  9 1783.6 1818.4 -882.79   1765.6 0.643      2     0.7251
```

### Linear mixed effects analysis of Peak 37

```
Model1 = lmer(CHC_data$Peak_37 ~ Sel.Line + Sex + Sel.Line*Sex + (1|Blocks) + (1|Time), data = CHC_data, REML = FALSE)
summary(Model1)
## Linear mixed model fit by maximum likelihood  ['lmerMod']
## Formula: CHC_data$Peak_37 ~ Sel.Line + Sex + Sel.Line * Sex + (1 | Blocks) +
##     (1 | Time)
##    Data: CHC_data
##
##      AIC      BIC   logLik deviance df.resid
##   -509.3   -474.5    263.6   -527.3      343
##
## Scaled residuals:
##     Min      1Q  Median      3Q     Max
## -3.8505 -0.5844 -0.0348  0.5622  4.5808
##
## Random effects:
##  Groups   Name        Variance  Std.Dev.
##  Time     (Intercept) 1.444e-02 0.1201670
##  Blocks   (Intercept) 3.136e-08 0.0001771
##  Residual             1.217e-02 0.1103364
## Number of obs: 352, groups:  Time, 6; Blocks, 3
##
## Fixed effects:
##                 Estimate Std. Error t value
## (Intercept)     0.627804   0.051119  12.281
## Sel.LineF       0.022446   0.020509   1.094
## Sel.LineM      -0.028350   0.020320  -1.395
## Sexm            0.110526   0.020320   5.439
## Sel.LineF:Sexm  0.006283   0.028936   0.217
## Sel.LineM:Sexm  0.032853   0.028674   1.146
##
## Correlation of Fixed Effects:
##             (Intr) Sl.LnF Sl.LnM Sexm   S.LF:S
## Sel.LineF   -0.197
## Sel.LineM   -0.199  0.496
## Sexm        -0.199  0.496  0.500
## Sel.LnF:Sxm  0.140 -0.709 -0.351 -0.703
## Sel.LnM:Sxm  0.141 -0.351 -0.709 -0.709  0.498
## convergence code: 0
## Model failed to converge with max|grad| = 0.00312769 (tol = 0.002, component 1)
Anova(Model1)
## Analysis of Deviance Table (Type II Wald chisquare tests)
##
## Response: CHC_data$Peak_37
##                 Chisq Df Pr(>Chisq)
## Sel.Line       6.9920  2    0.03032 *
## Sex          110.5702  1    < 2e-16 ***
## Sel.Line:Sex   1.4786  2    0.47744
## ---
## Signif. codes:  0 '***' 0.001 '**' 0.01 '*' 0.05 '.' 0.1 ' ' 1
Model2 = lmer(CHC_data$Peak_37 ~ Sel.Line + Sex  + (1|Blocks) + (1|Time), data = CHC_data, REML = FALSE)
summary(Model2)
## Linear mixed model fit by maximum likelihood  ['lmerMod']
## Formula: CHC_data$Peak_37 ~ Sel.Line + Sex + (1 | Blocks) + (1 | Time)
##    Data: CHC_data
##
##      AIC      BIC   logLik deviance df.resid
##   -511.8   -484.8    262.9   -525.8      345
##
## Scaled residuals:
##     Min      1Q  Median      3Q     Max
## -3.8750 -0.5785  0.0003  0.5435  4.5405
##
## Random effects:
##  Groups   Name        Variance  Std.Dev.
##  Time     (Intercept) 1.444e-02 1.202e-01
##  Blocks   (Intercept) 2.516e-10 1.586e-05
##  Residual             1.223e-02 1.106e-01
## Number of obs: 352, groups:  Time, 6; Blocks, 3
##
## Fixed effects:
##             Estimate Std. Error t value
## (Intercept)  0.62122    0.05045  12.314
## Sel.LineF    0.02556    0.01449   1.763
## Sel.LineM   -0.01184    0.01437  -0.824
## Sexm         0.12369    0.01179  10.493
##
## Correlation of Fixed Effects:
##           (Intr) Sl.LnF Sl.LnM
## Sel.LineF -0.141
## Sel.LineM -0.143  0.498
## Sexm      -0.117 -0.004 -0.003
Anova(Model2)
## Analysis of Deviance Table (Type II Wald chisquare tests)
##
## Response: CHC_data$Peak_37
##             Chisq Df Pr(>Chisq)
## Sel.Line   6.9622  2    0.03077 *
## Sex      110.0995  1    < 2e-16 ***
## ---
## Signif. codes:  0 '***' 0.001 '**' 0.01 '*' 0.05 '.' 0.1 ' ' 1
anova(Model1, Model2)
## Data: CHC_data
## Models:
## Model2: CHC_data$Peak_37 ~ Sel.Line + Sex + (1 | Blocks) + (1 | Time)
## Model1: CHC_data$Peak_37 ~ Sel.Line + Sex + Sel.Line * Sex + (1 | Blocks) +
## Model1:     (1 | Time)
##        Df     AIC     BIC logLik deviance  Chisq Chi Df Pr(>Chisq)
## Model2  7 -511.82 -484.78 262.91  -525.82
## Model1  9 -509.30 -474.53 263.65  -527.30 1.4755      2     0.4782
```

### Linear mixed effects analysis of Peak 38

```
Model1 = lmer(CHC_data$Peak_38 ~ Sel.Line + Sex + Sel.Line*Sex + (1|Blocks) + (1|Time), data = CHC_data, REML = FALSE)
summary(Model1)
## Linear mixed model fit by maximum likelihood  ['lmerMod']
## Formula: CHC_data$Peak_38 ~ Sel.Line + Sex + Sel.Line * Sex + (1 | Blocks) +
##     (1 | Time)
##    Data: CHC_data
##
##      AIC      BIC   logLik deviance df.resid
##    331.7    366.5   -156.8    313.7      343
##
## Scaled residuals:
##     Min      1Q  Median      3Q     Max
## -1.8098 -0.4715 -0.0459  0.3015 12.6587
##
## Random effects:
##  Groups   Name        Variance Std.Dev.
##  Time     (Intercept) 0.0179   0.1338
##  Blocks   (Intercept) 0.0000   0.0000
##  Residual             0.1376   0.3709
## Number of obs: 352, groups:  Time, 6; Blocks, 3
##
## Fixed effects:
##                Estimate Std. Error t value
## (Intercept)     1.32585    0.07292  18.183
## Sel.LineF      -0.09364    0.06894  -1.358
## Sel.LineM      -0.15000    0.06831  -2.196
## Sexm           -0.46926    0.06831  -6.869
## Sel.LineF:Sexm  0.07222    0.09727   0.742
## Sel.LineM:Sexm  0.20341    0.09640   2.110
##
## Correlation of Fixed Effects:
##             (Intr) Sl.LnF Sl.LnM Sexm   S.LF:S
## Sel.LineF   -0.465
## Sel.LineM   -0.469  0.496
## Sexm        -0.469  0.496  0.500
## Sel.LnF:Sxm  0.329 -0.709 -0.351 -0.703
## Sel.LnM:Sxm  0.332 -0.351 -0.709 -0.709  0.498
## convergence code: 0
## boundary (singular) fit: see ?isSingular
Anova(Model1)
## Analysis of Deviance Table (Type II Wald chisquare tests)
##
## Response: CHC_data$Peak_38
##                Chisq Df Pr(>Chisq)
## Sel.Line      1.6183  2     0.4452
## Sex          90.8442  1     <2e-16 ***
## Sel.Line:Sex  4.5788  2     0.1013
## ---
## Signif. codes:  0 '***' 0.001 '**' 0.01 '*' 0.05 '.' 0.1 ' ' 1
Model2 = lmer(CHC_data$Peak_38 ~ Sel.Line + Sex  + (1|Blocks) + (1|Time), data = CHC_data, REML = FALSE)
summary(Model2)
## Linear mixed model fit by maximum likelihood  ['lmerMod']
## Formula: CHC_data$Peak_38 ~ Sel.Line + Sex + (1 | Blocks) + (1 | Time)
##    Data: CHC_data
##
##      AIC      BIC   logLik deviance df.resid
##    332.2    359.3   -159.1    318.2      345
##
## Scaled residuals:
##     Min      1Q  Median      3Q     Max
## -1.9501 -0.4902 -0.0588  0.2829 12.4259
##
## Random effects:
##  Groups   Name        Variance  Std.Dev.
##  Time     (Intercept) 1.798e-02 1.341e-01
##  Blocks   (Intercept) 1.130e-09 3.361e-05
##  Residual             1.394e-01 3.734e-01
## Number of obs: 352, groups:  Time, 6; Blocks, 3
##
## Fixed effects:
##             Estimate Std. Error t value
## (Intercept)  1.27967    0.06764  18.919
## Sel.LineF   -0.05760    0.04894  -1.177
## Sel.LineM   -0.04781    0.04851  -0.986
## Sexm        -0.37690    0.03980  -9.469
##
## Correlation of Fixed Effects:
##           (Intr) Sl.LnF Sl.LnM
## Sel.LineF -0.356
## Sel.LineM -0.359  0.498
## Sexm      -0.294 -0.004 -0.003
## convergence code: 0
## boundary (singular) fit: see ?isSingular
Anova(Model2)
## Analysis of Deviance Table (Type II Wald chisquare tests)
##
## Response: CHC_data$Peak_38
##            Chisq Df Pr(>Chisq)
## Sel.Line  1.5976  2     0.4499
## Sex      89.6668  1     <2e-16 ***
## ---
## Signif. codes:  0 '***' 0.001 '**' 0.01 '*' 0.05 '.' 0.1 ' ' 1
anova(Model1, Model2)
## Data: CHC_data
## Models:
## Model2: CHC_data$Peak_38 ~ Sel.Line + Sex + (1 | Blocks) + (1 | Time)
## Model1: CHC_data$Peak_38 ~ Sel.Line + Sex + Sel.Line * Sex + (1 | Blocks) +
## Model1:     (1 | Time)
##        Df    AIC    BIC  logLik deviance  Chisq Chi Df Pr(>Chisq)
## Model2  7 332.24 359.28 -159.12   318.24
## Model1  9 331.69 366.46 -156.84   313.69 4.5491      2     0.1028
```
