## Supplementary material for "Cuticular hydrocarbon divergence in *Drosophila melanogaster* populations evolving under differential operational sex ratios": SI2

Journal Name: Evolutionary Biology

Dutta^1*^, Tejinder Singh Chechi^1^, Ankit Yadav^2^, Nagaraj Guru Prasad^1^

^1^Department of Biological Sciences, Indian Institute of Science Education and Research, Mohali, India

^2^Department of Earth and Environmental Sciences, Indian Institute of Science Education and Research, Mohali, India

**Boxplots showing sex-wise differences in *D. melanogaster* MCF cuticular hydrocarbon abundance. Those cuticular hydrocarbons have been considered that are affected by selection lines but not by time of extraction as predictor variables in the linear mixed effect model**

The Table 3 in the main article summarises the analysis that has been detailed here. There are three different plots from three different analysis for each cuticular hydrocarbon peak that has been presented here together under the title bearing peak identity. The first analysis tests sexual dimorphism in the abundance of a particular cuticular hydrocarbon peak for each selection regime. The second and third analysis tests differences in the abundance of a particular cuticular hydrocarbon peak in females and males of each selection regime respectively. All between group comparisons tests differences between means of distribution of CHC abundance (ng/ml) using Wilcoxon test.

Peak 2

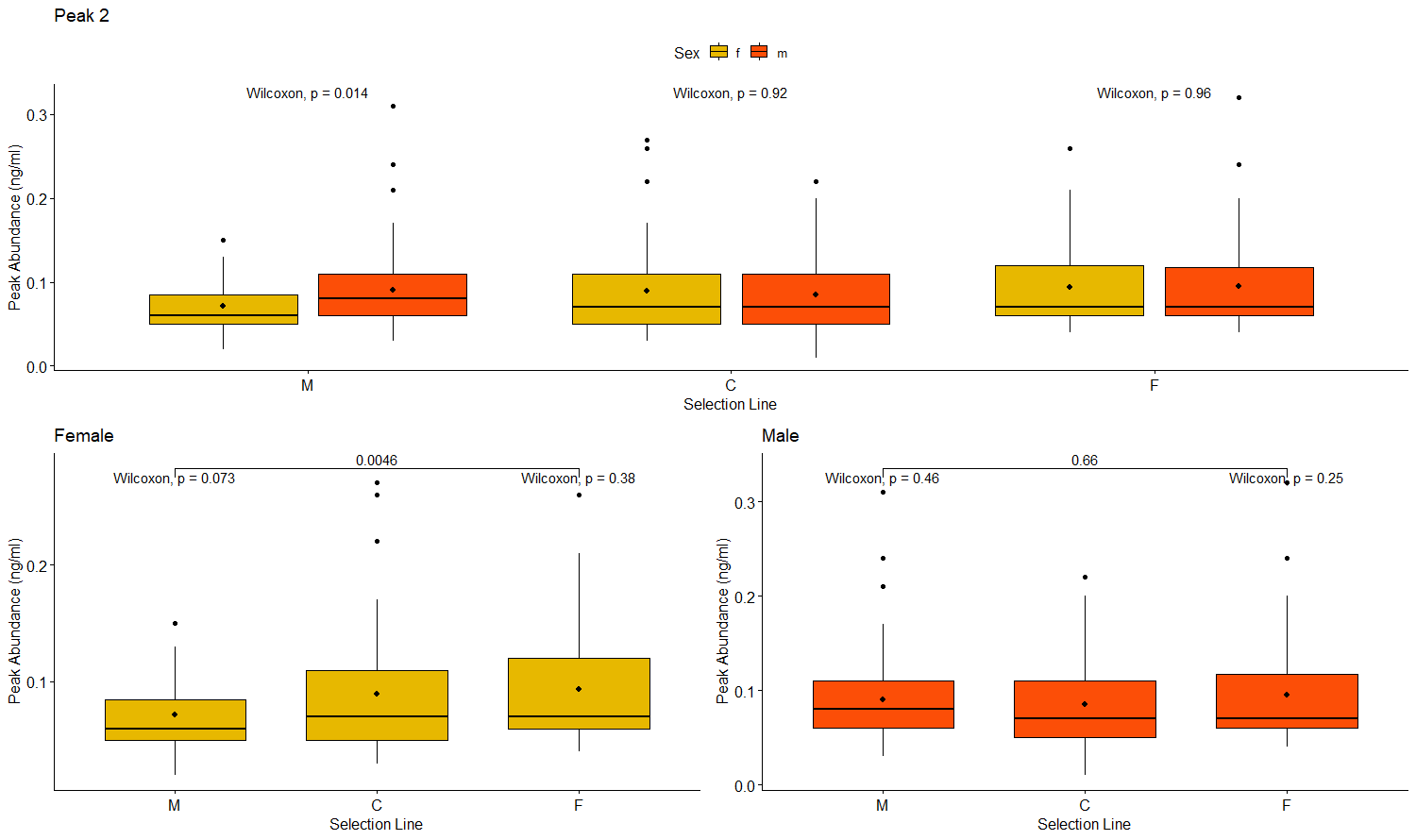

Peak 9

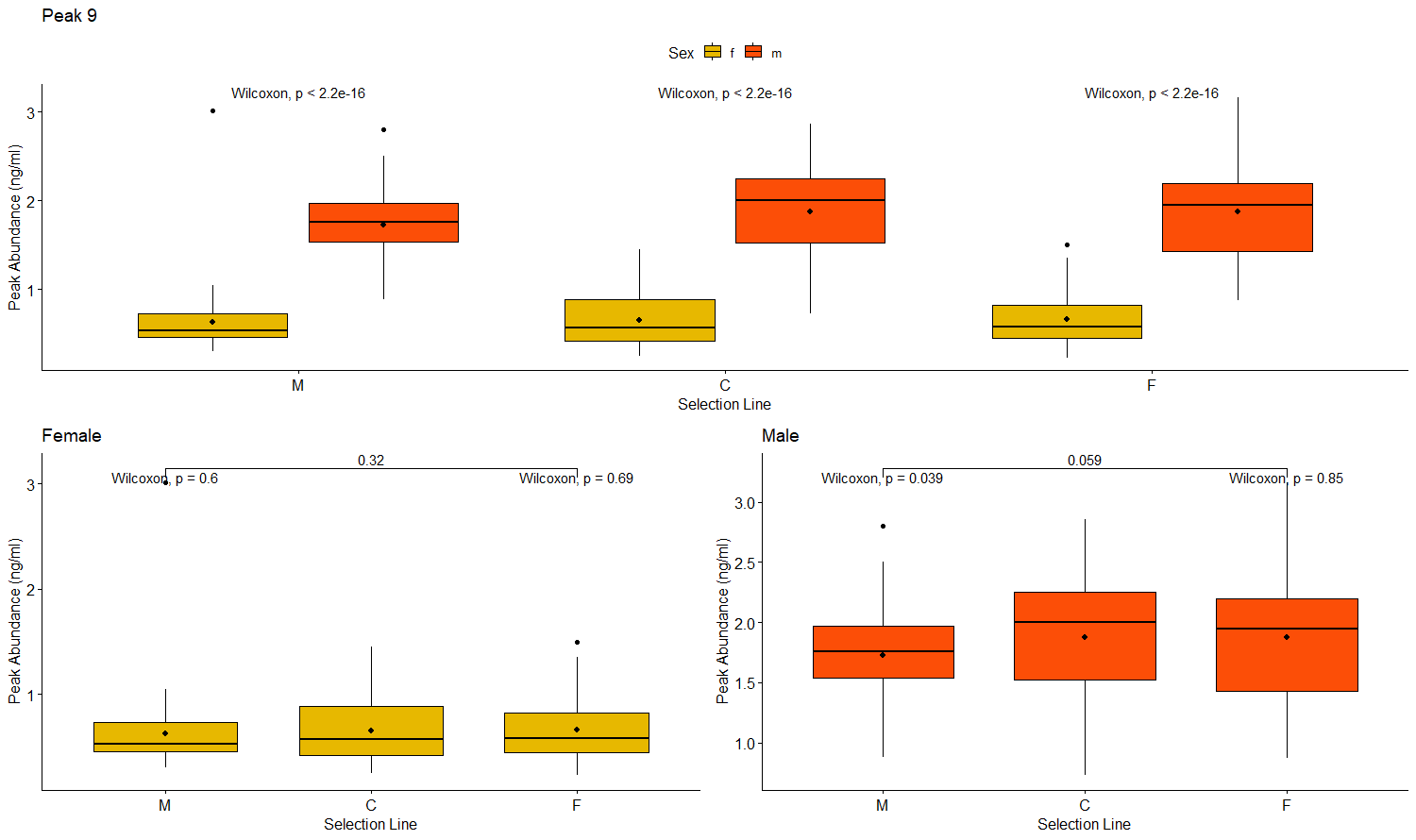

Peak 12 (7-tricosene (in male) and 5,9/7,11-tricosadiene (in female))

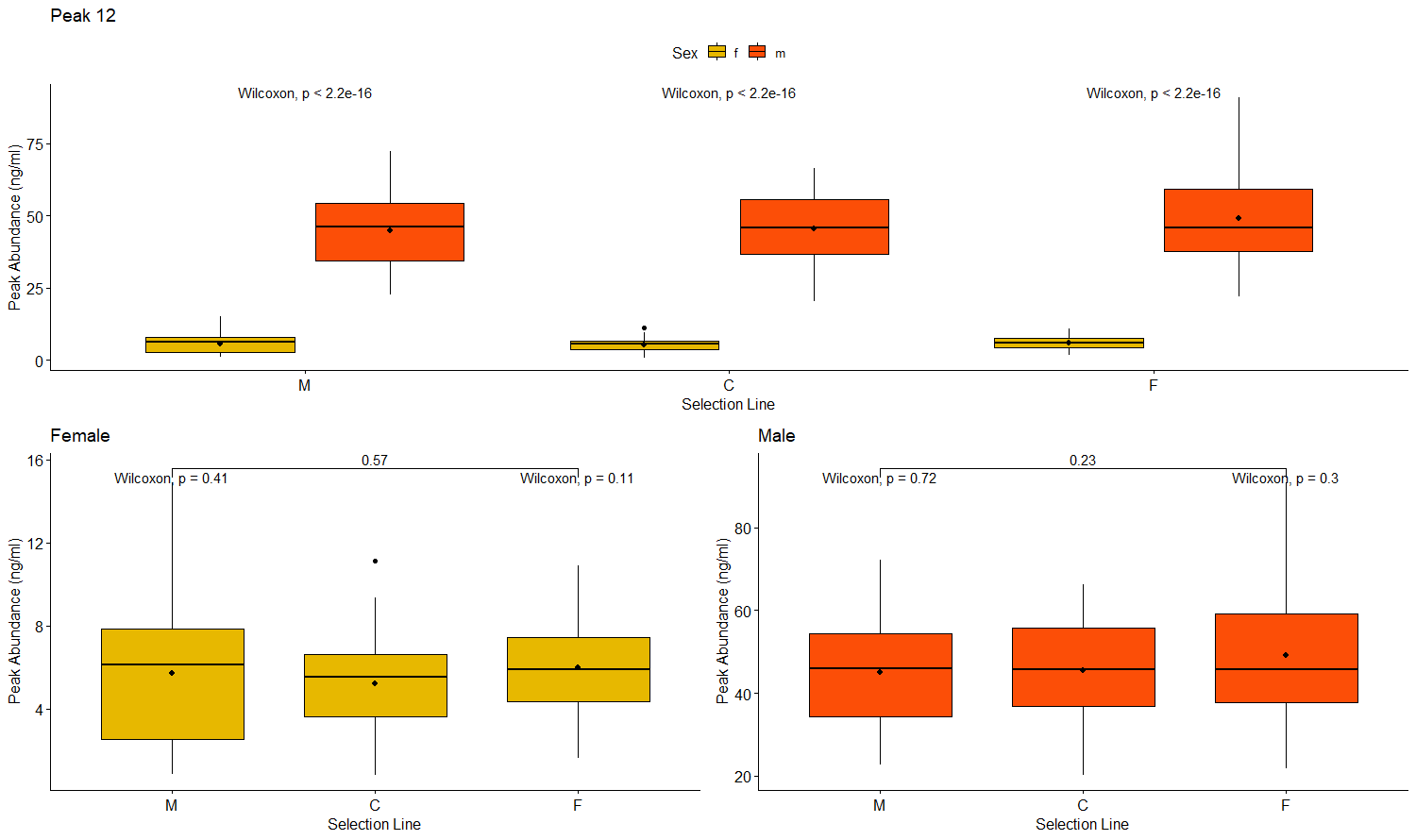

Peak 14

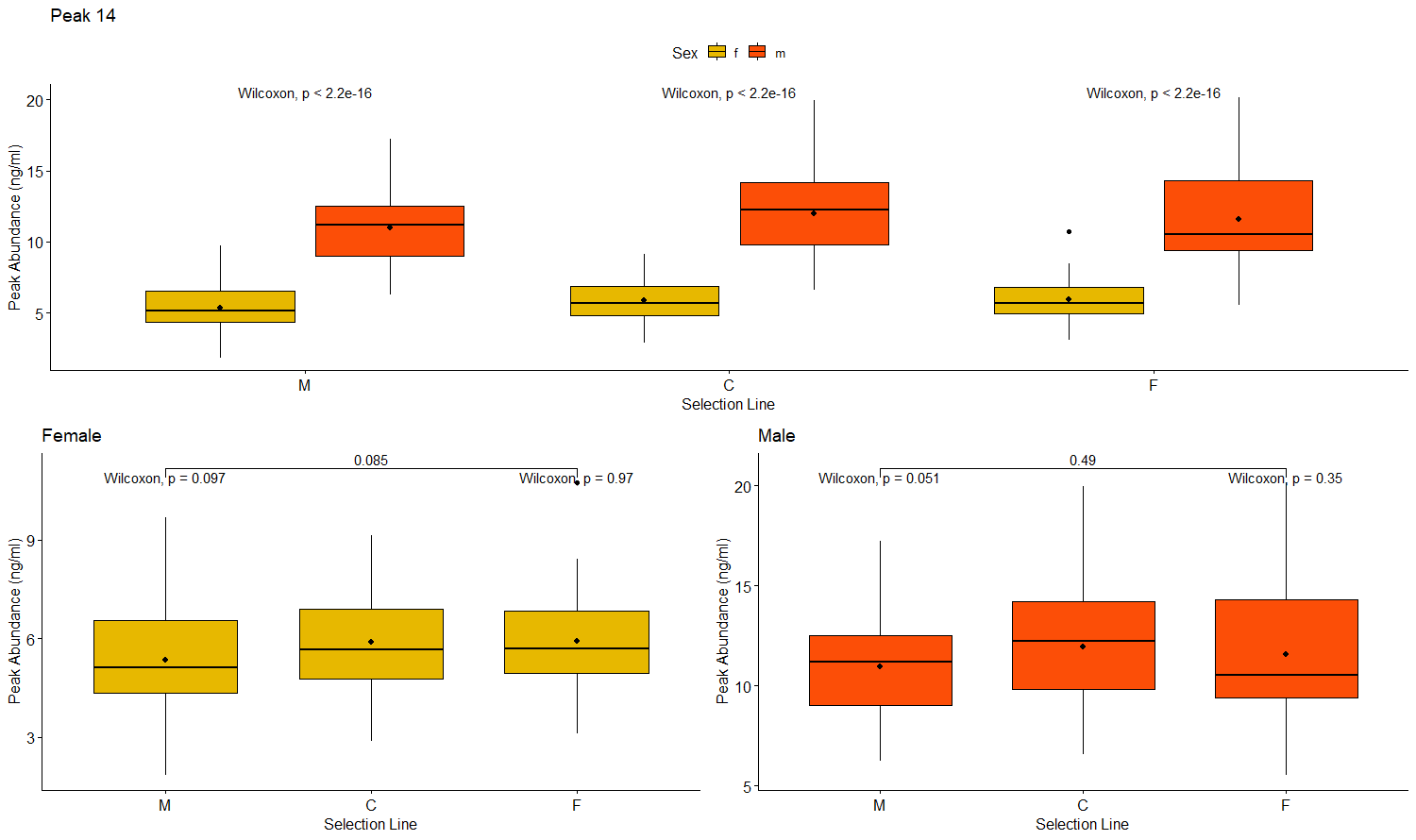

Peak 15

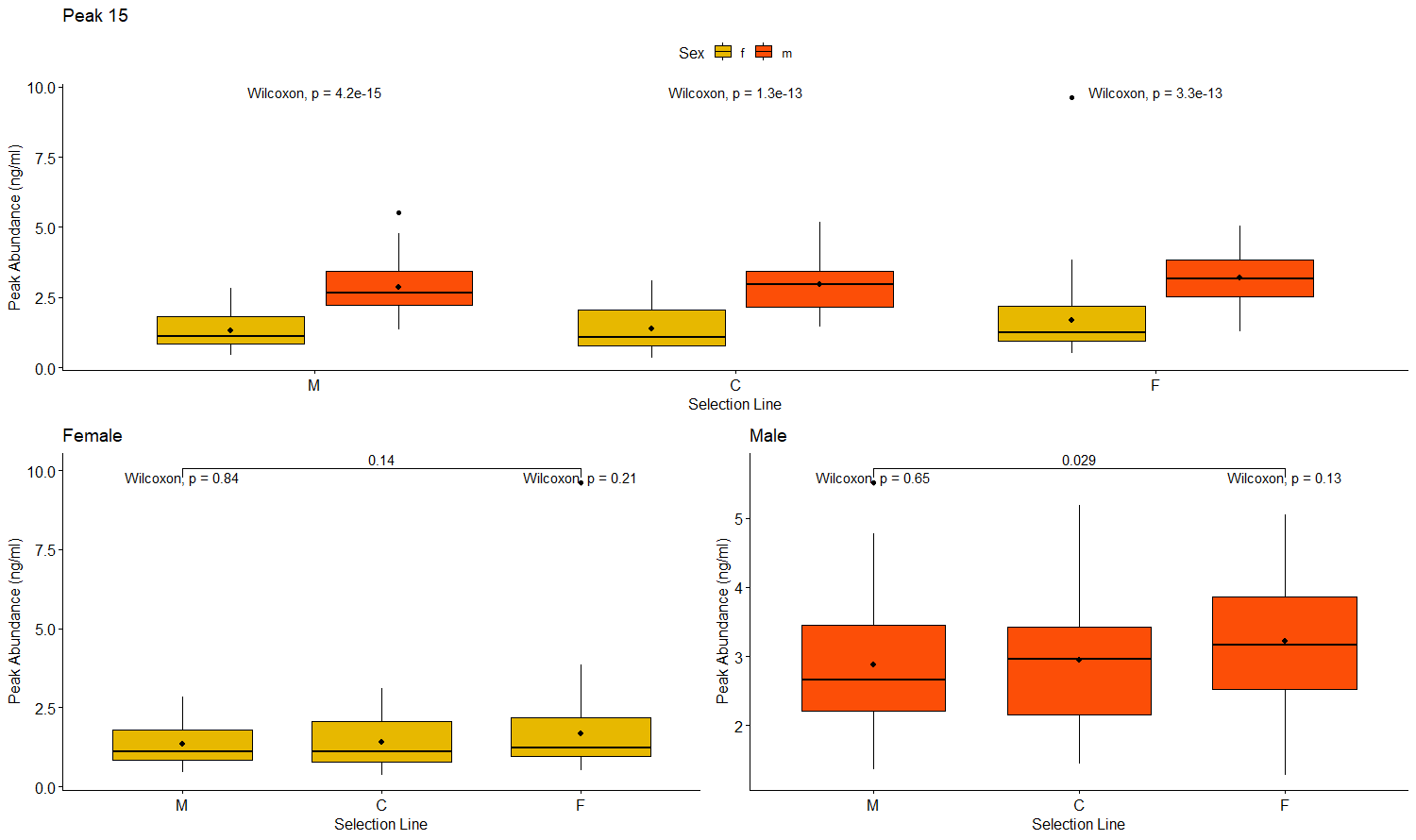

Peak 19

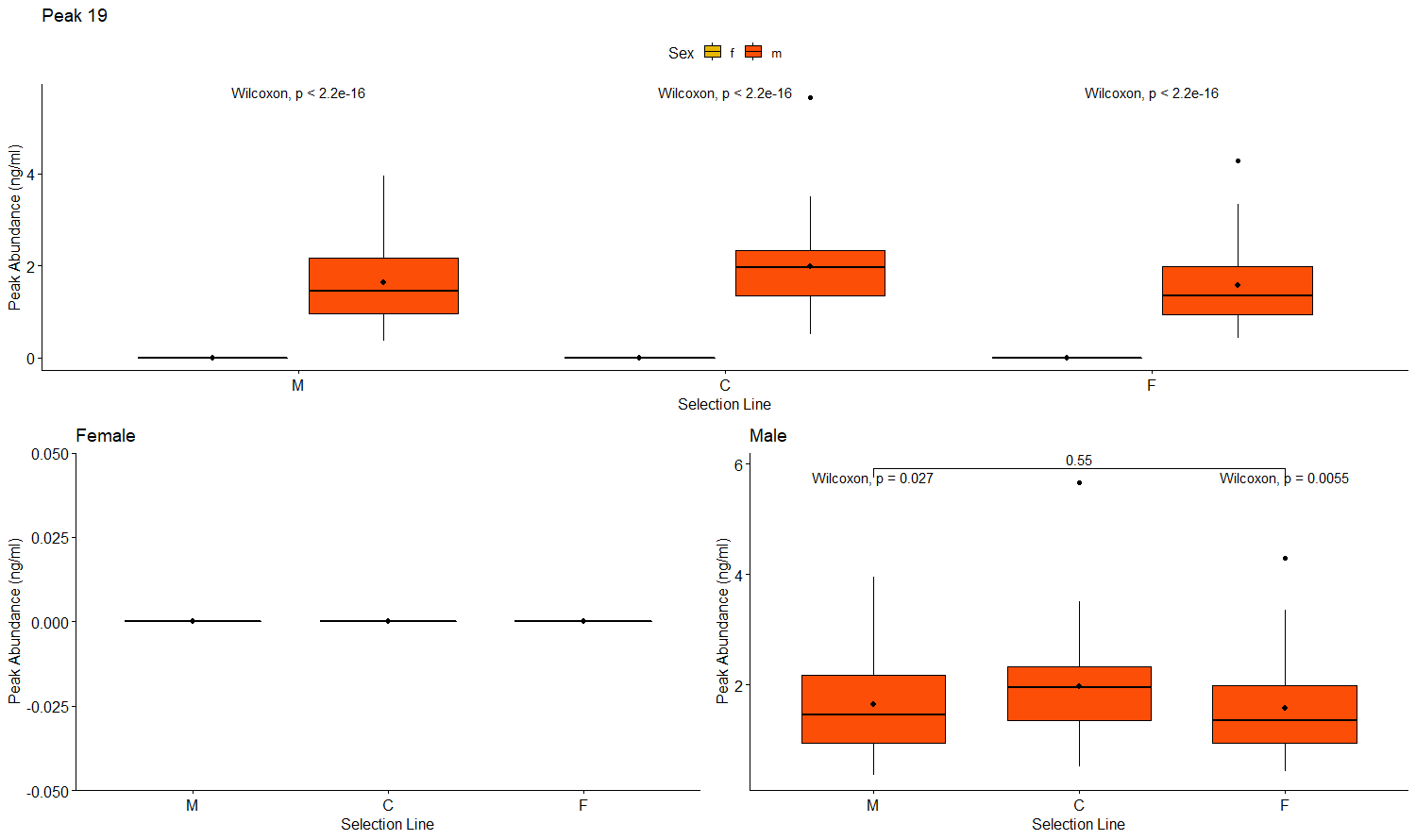

Peak 20 (9-pentacosene)

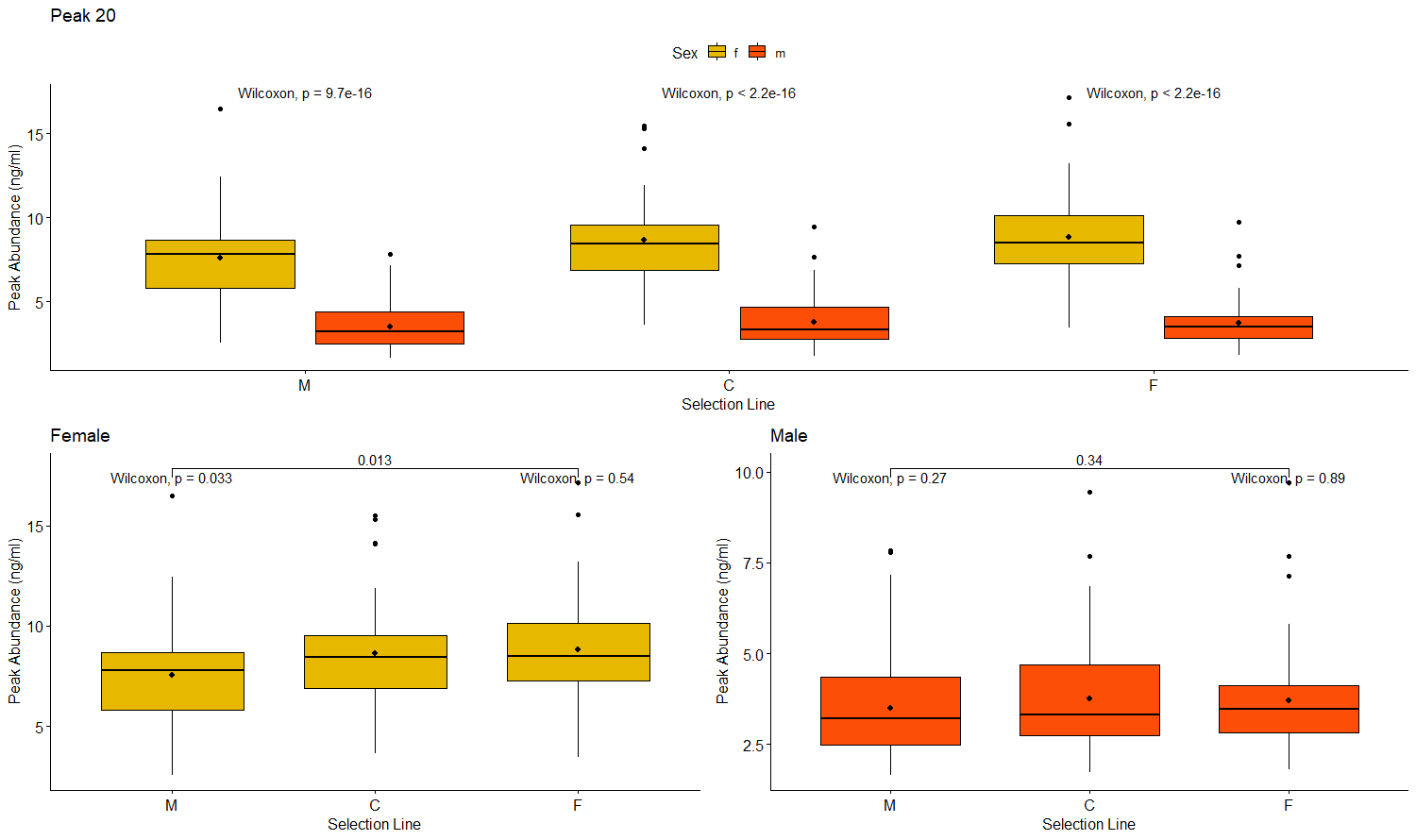

Peak 21 (7-pentacosene (in male) and 5,9/7,11-pentacosadiene (in female))

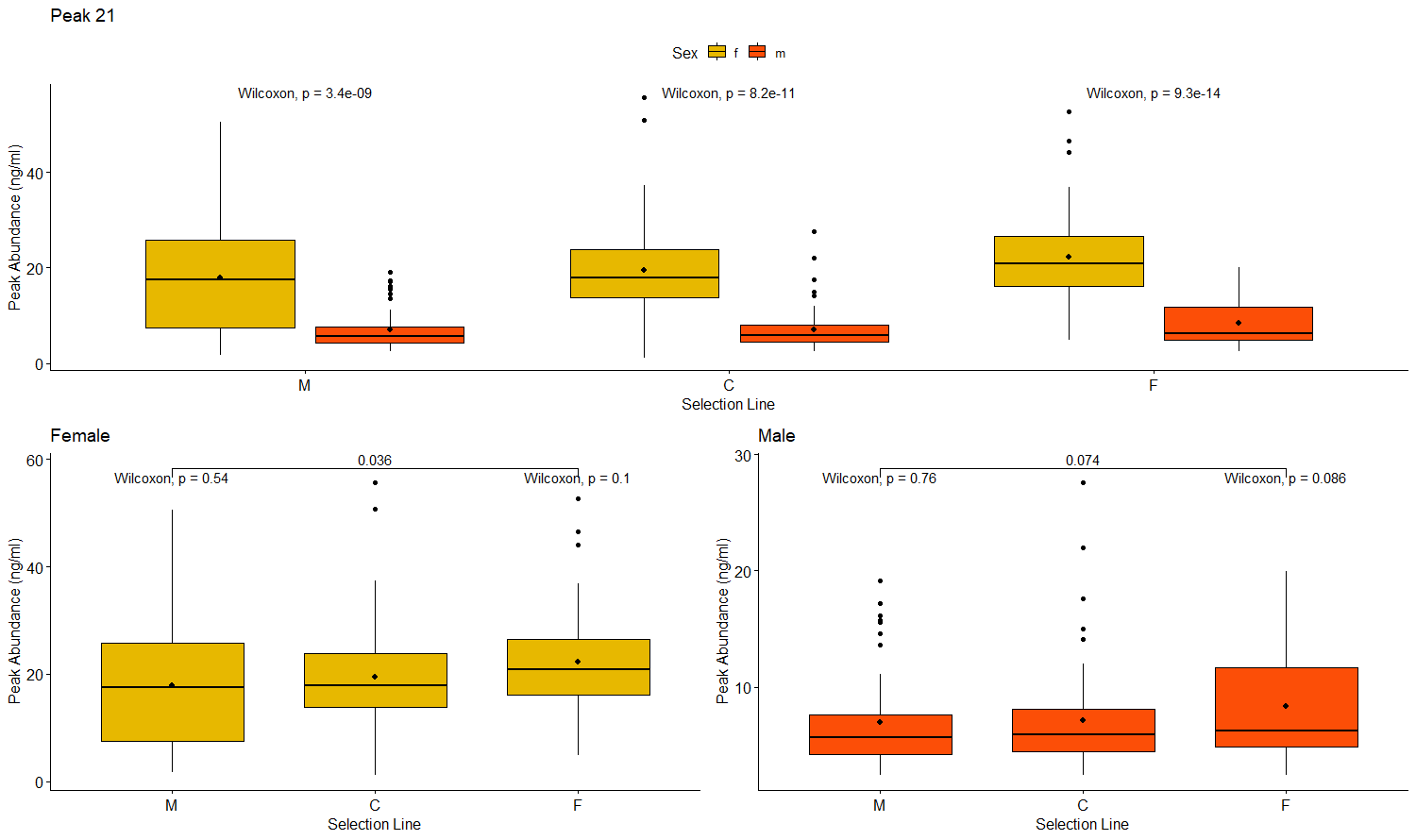

Peak 22

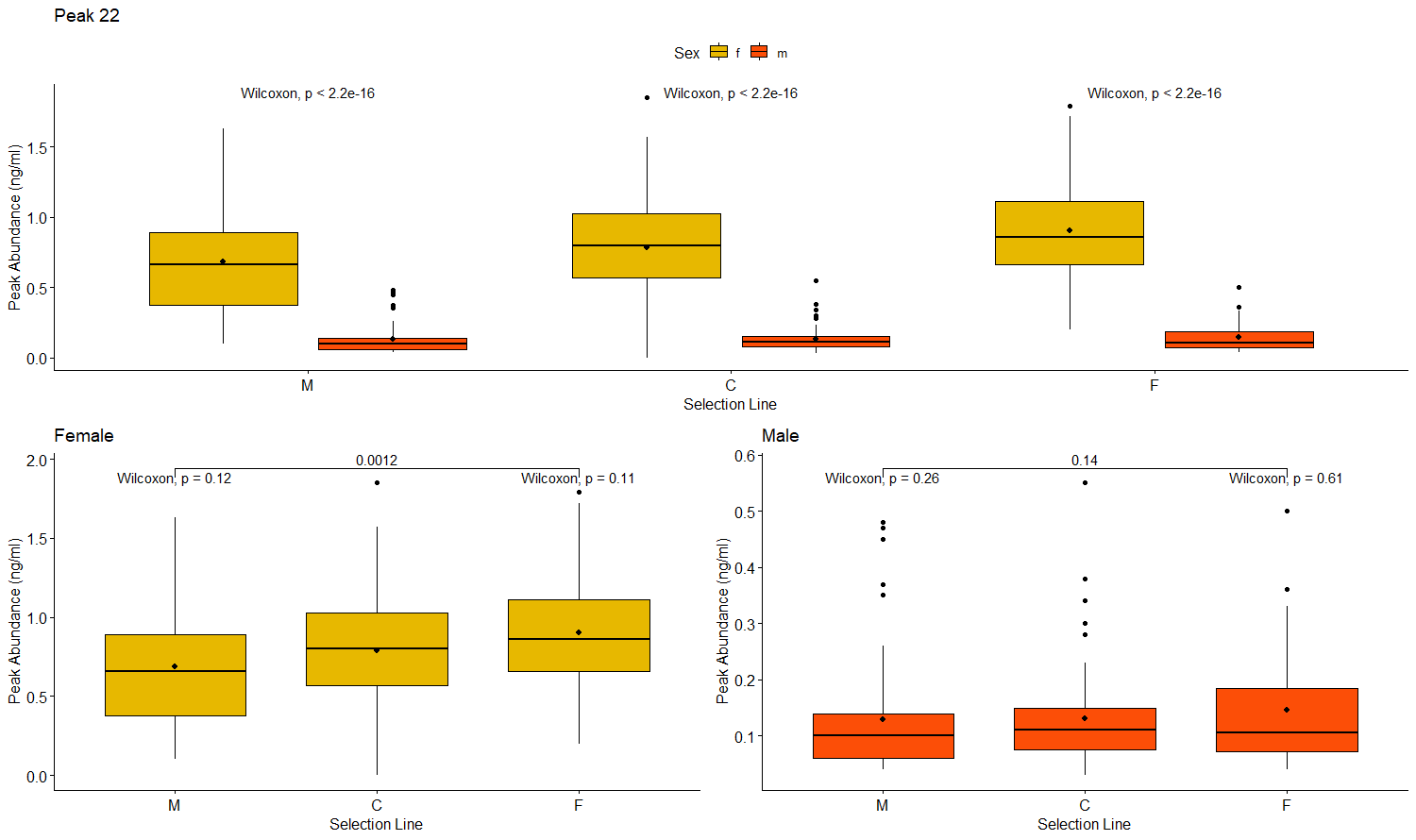

Peak 23

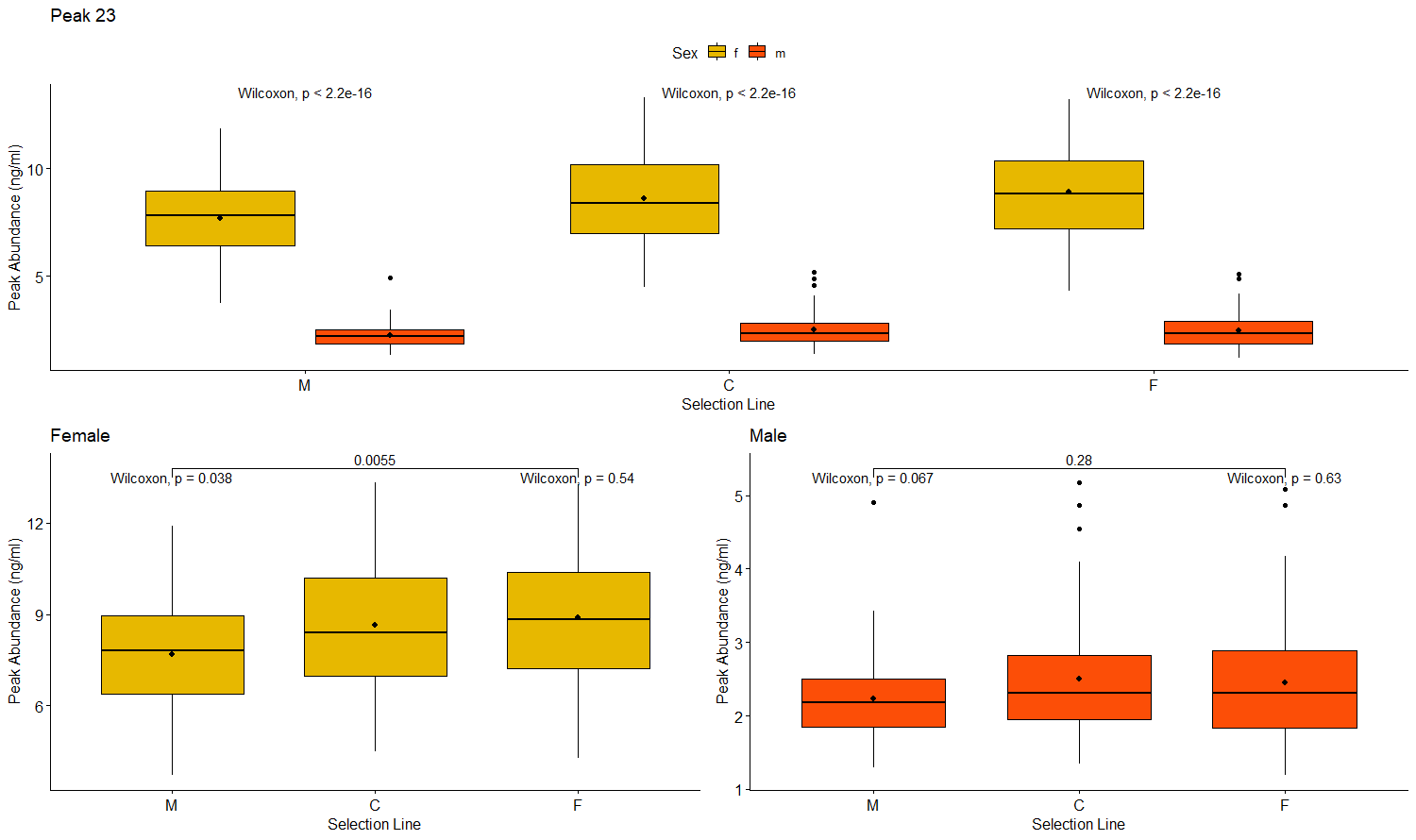

Peak 25

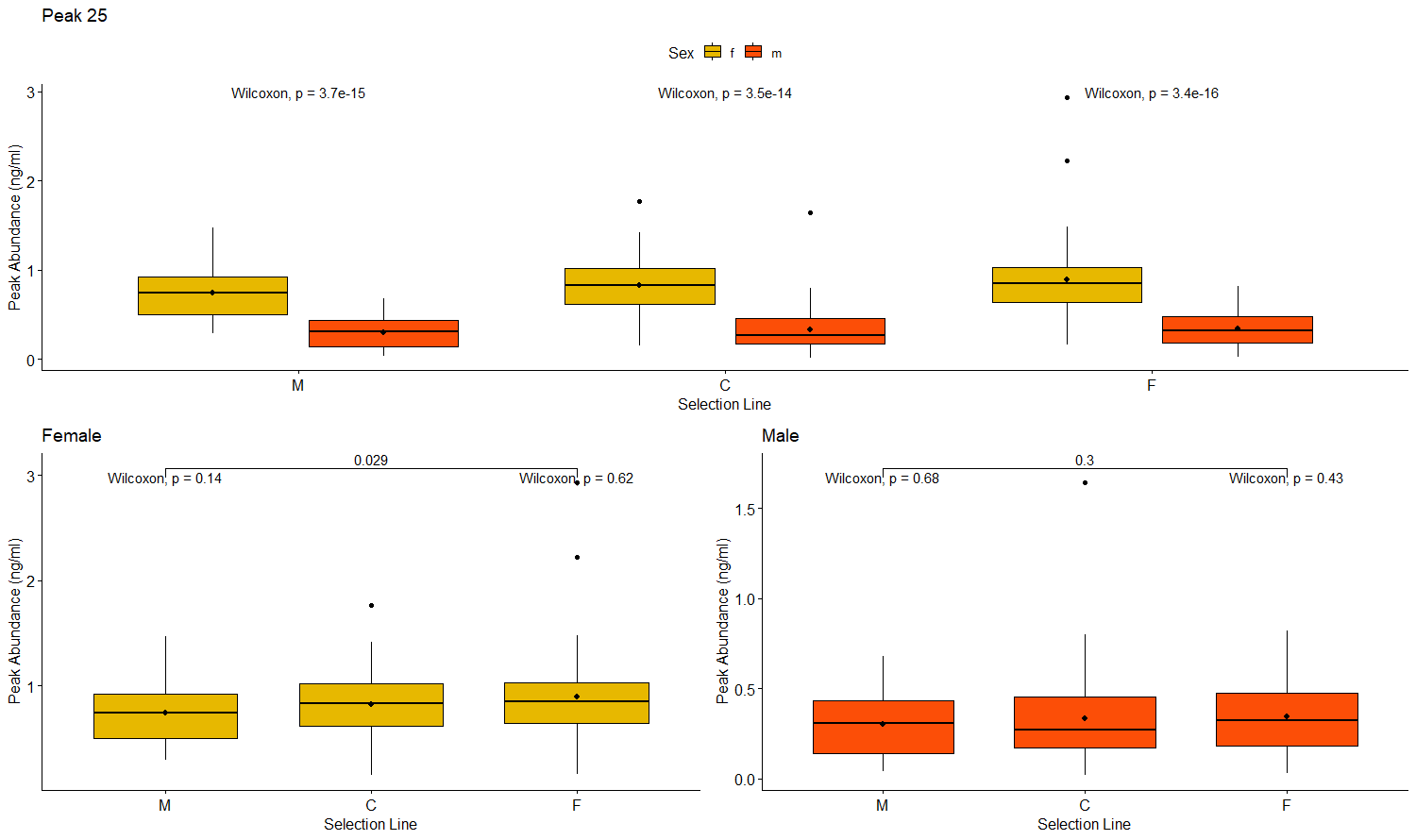

Peak 27

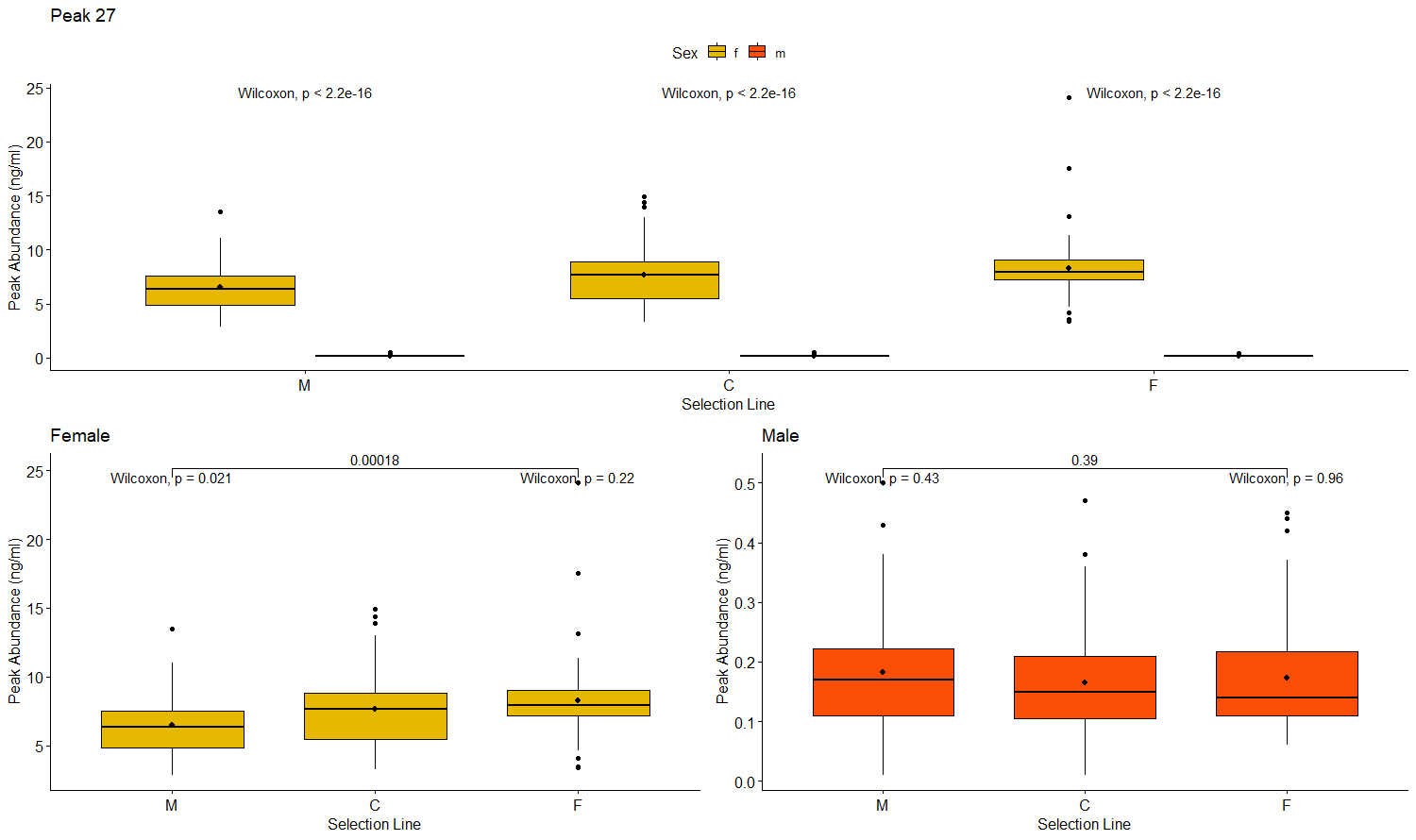

Peak 28 (7-heptacosene (in male) and 5,9/7,11-heptacosadiene (in female))

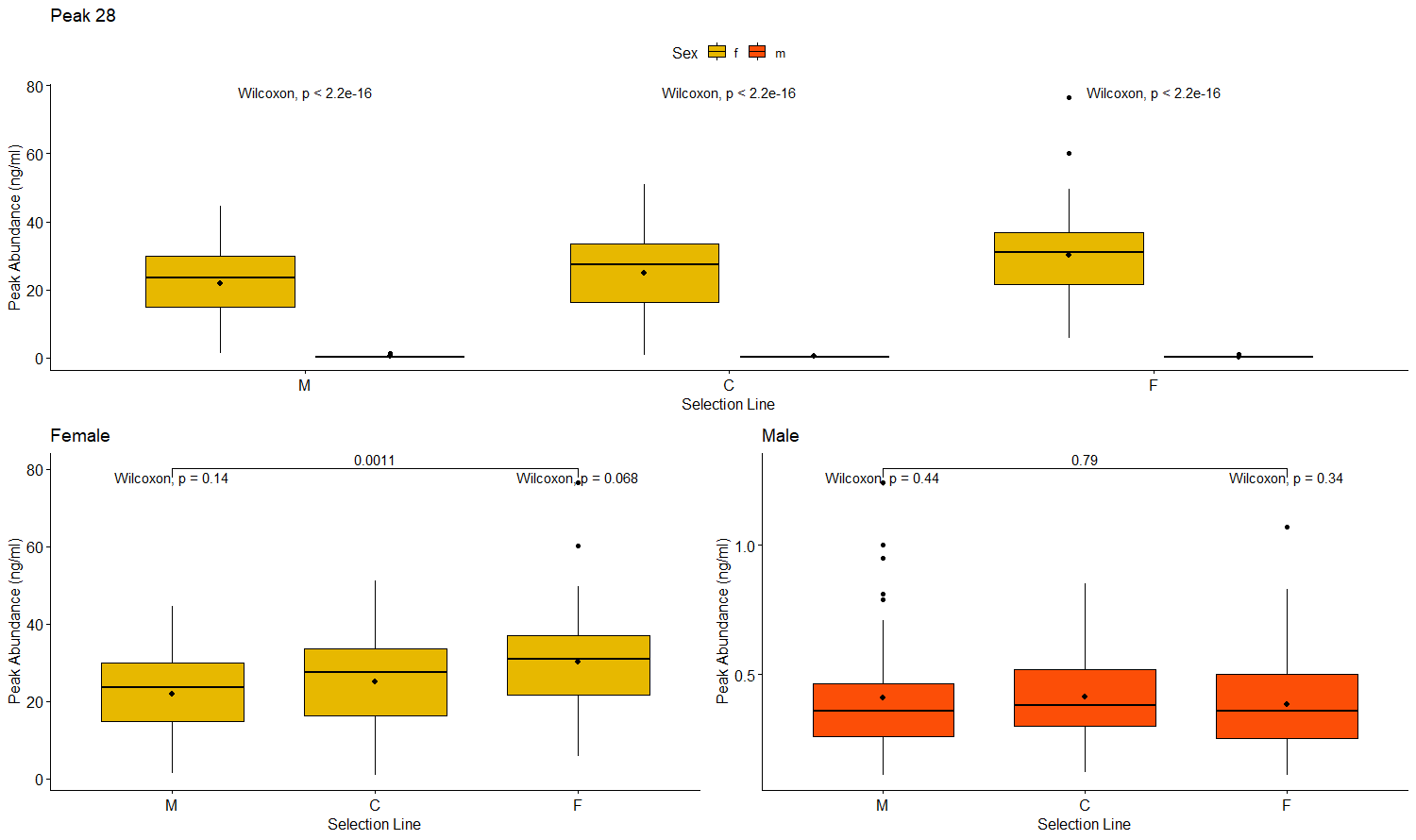

Peak 29

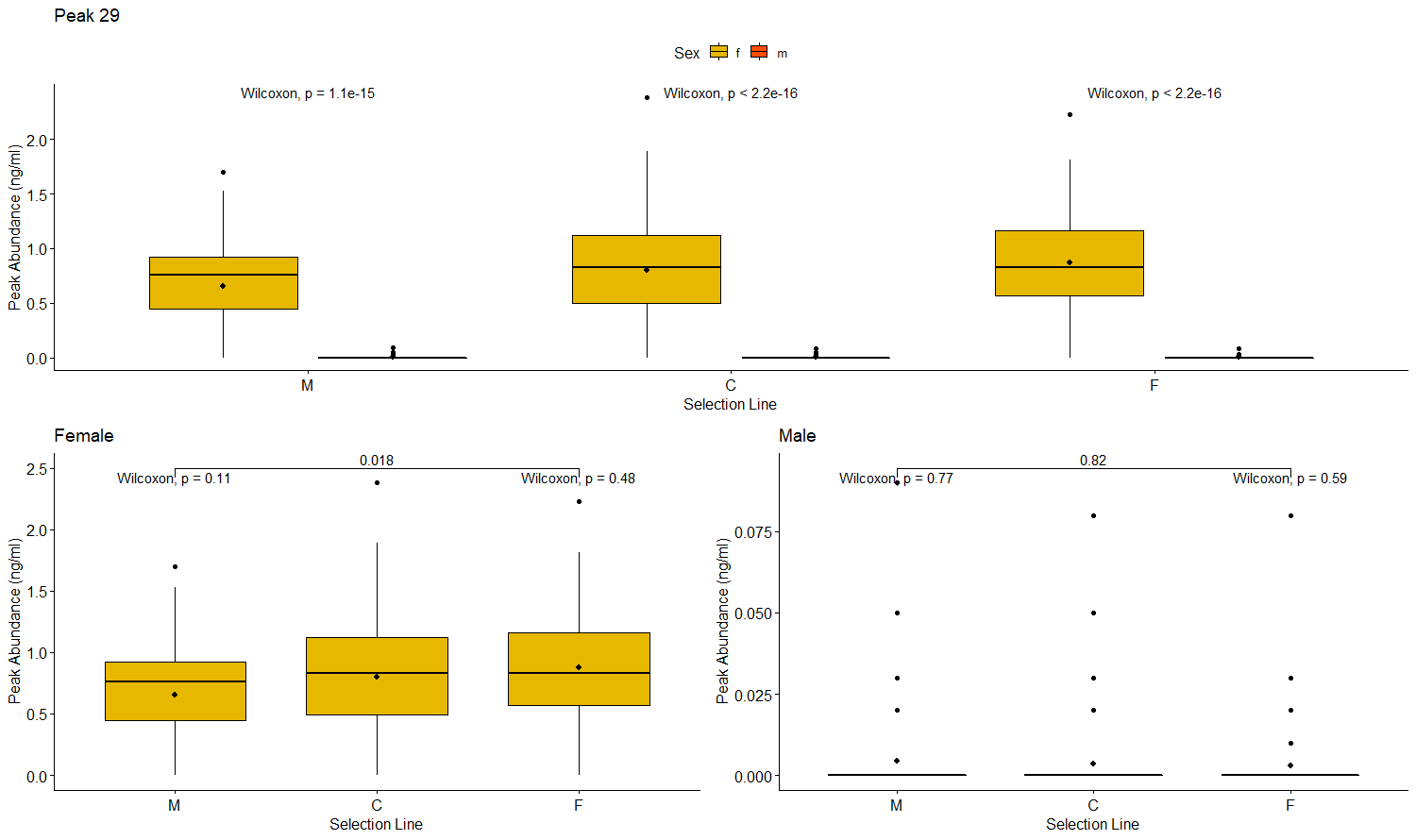

Peak 30

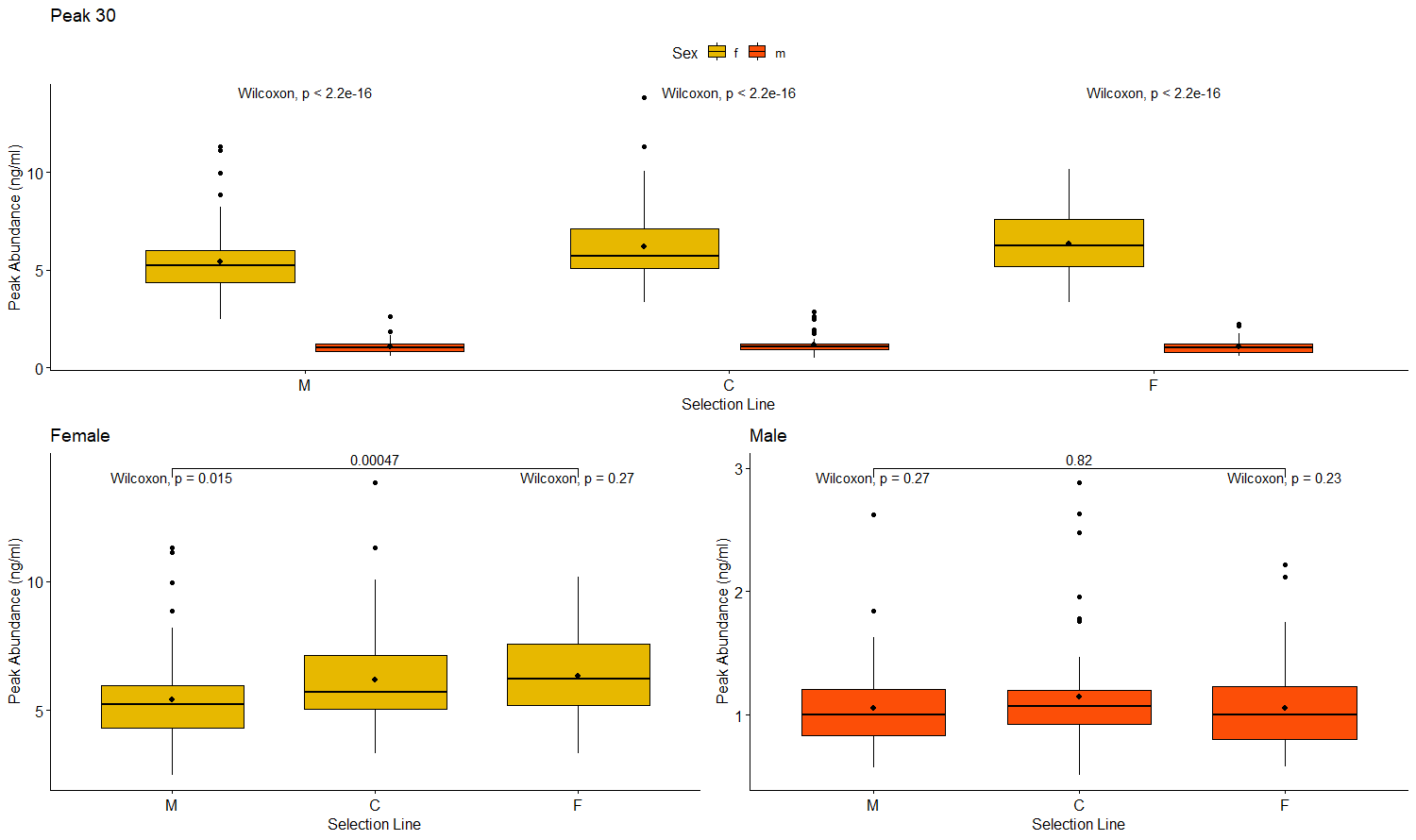

Peak 34

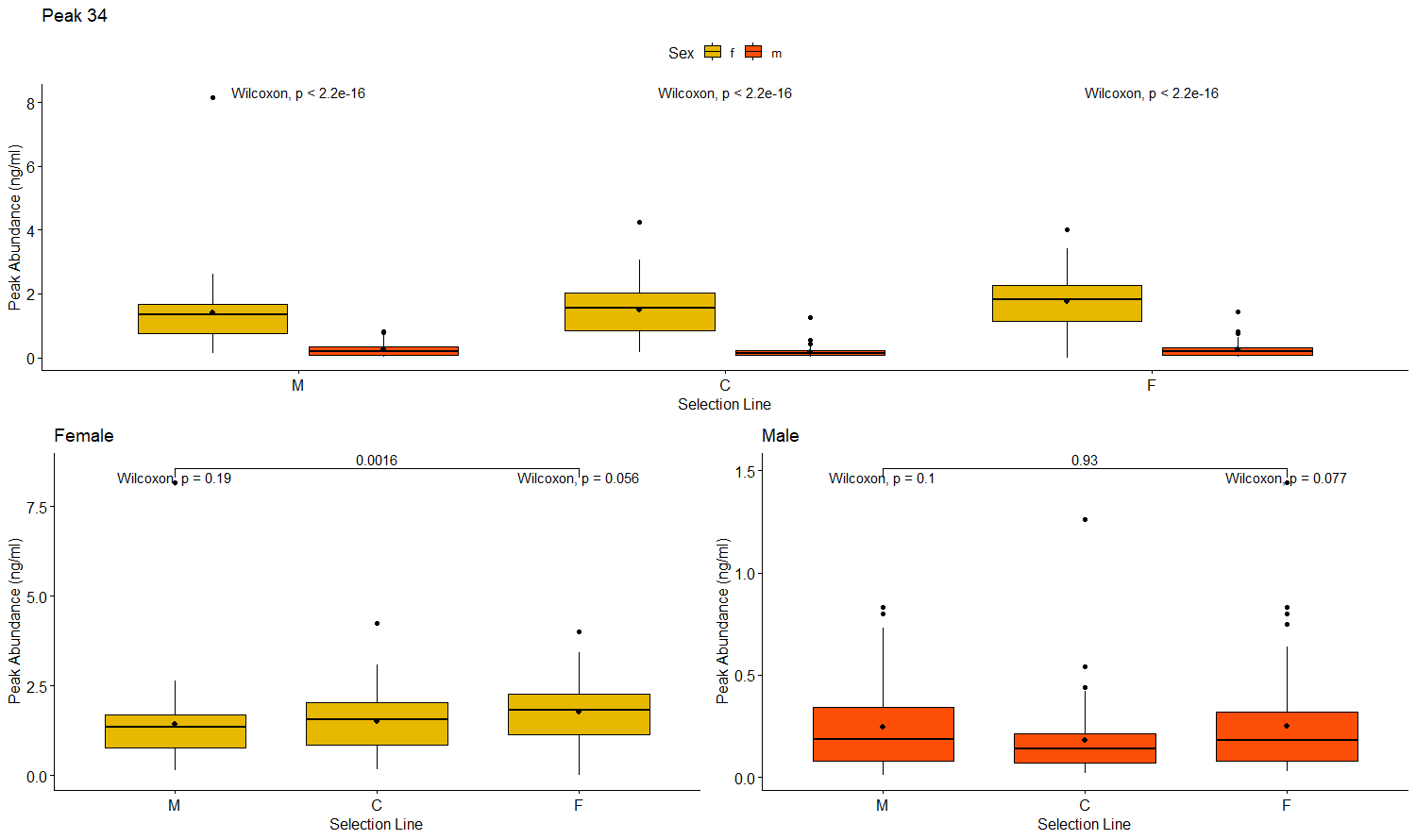
