## Supplementary material for "Cuticular hydrocarbon divergence in *Drosophila melanogaster* populations evolving under differential operational sex ratios": SI3

Journal Name: Evolutionary Biology

Dutta^1*^, Tejinder Singh Chechi^1^, Ankit Yadav^2^, Nagaraj Guru Prasad^1^

^1^Department of Biological Sciences, Indian Institute of Science Education and Research, Mohali, India

^2^Department of Earth and Environmental Sciences, Indian Institute of Science Education and Research, Mohali, India

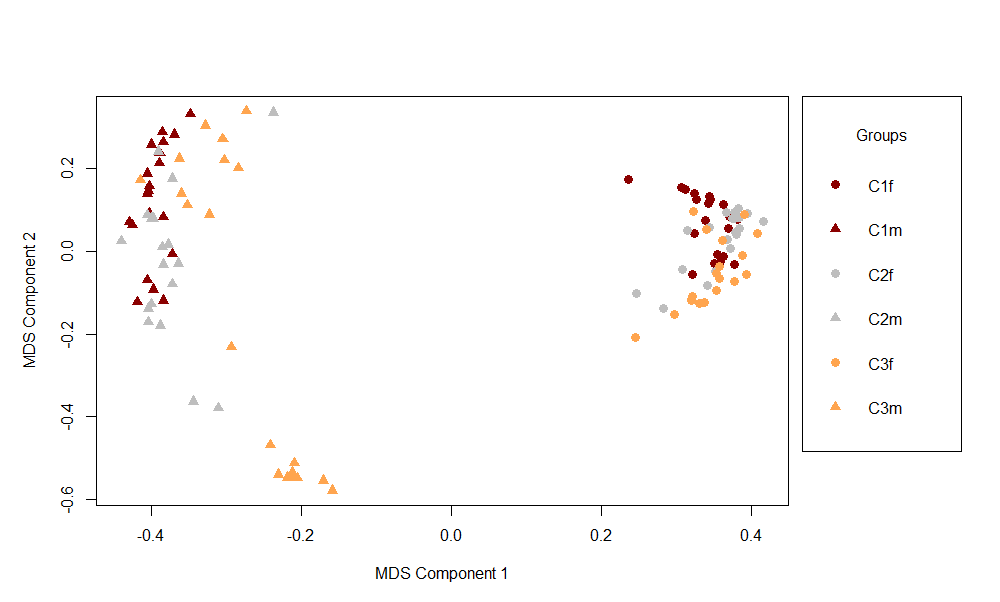
SI Fig. 1 Multi-dimensional scaling (MDS) of the proximity matrix of C individuals based on 16 *D. melanogaster* cuticular hydrocarbon peaks that were affected by selection but not time of extraction in linear mixed effect analysis (C_1/2/3_ are C blocks, m= males and f= females)

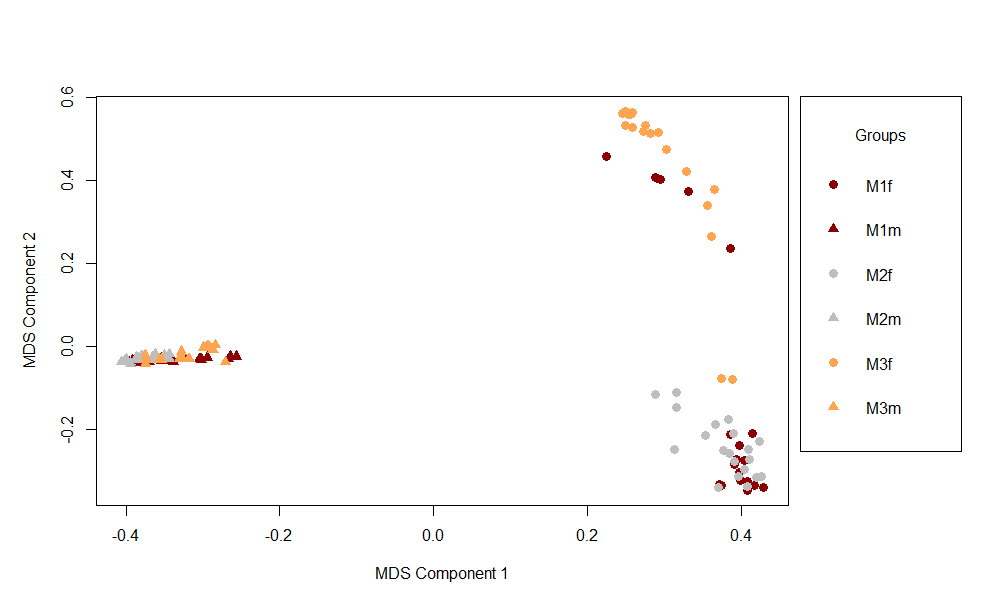
SI Fig. 2 Multi-dimensional scaling (MDS) of the proximity matrix of M individuals based on 16 *D. melanogaster* cuticular hydrocarbon peaks that were affected by selection but not time of extraction in linear mixed effect analysis (M_1/2/3_ are M blocks, m= males and f= females)

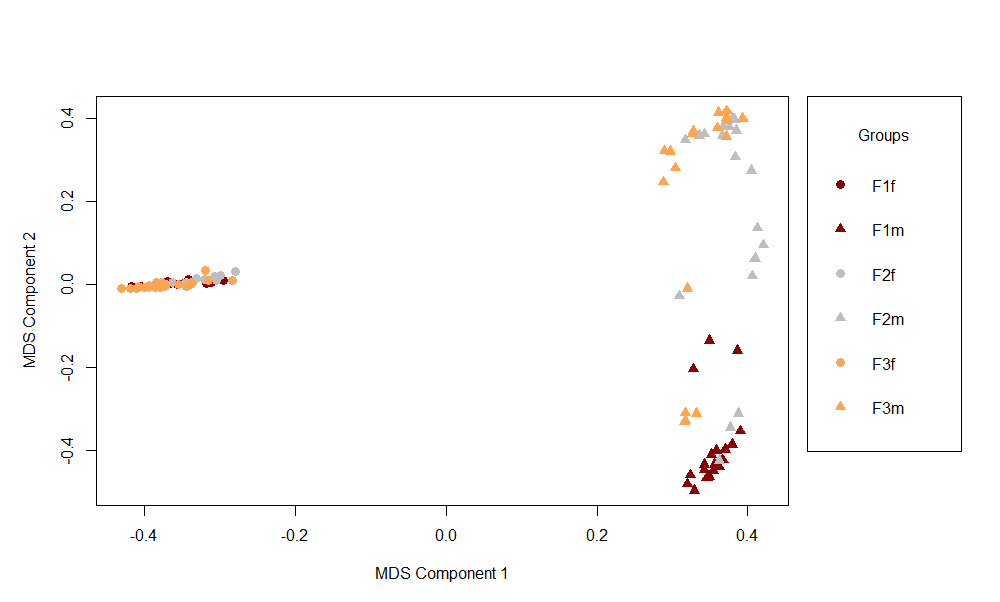
SI Fig. 3 Multi-dimensional scaling (MDS) of the proximity matrix of F individuals based on 16 *D. melanogaster* cuticular hydrocarbon peaks that were affected by selection but not time of extraction in linear mixed effect analysis (F_1/2/3_ are F blocks, m= males and f= females)

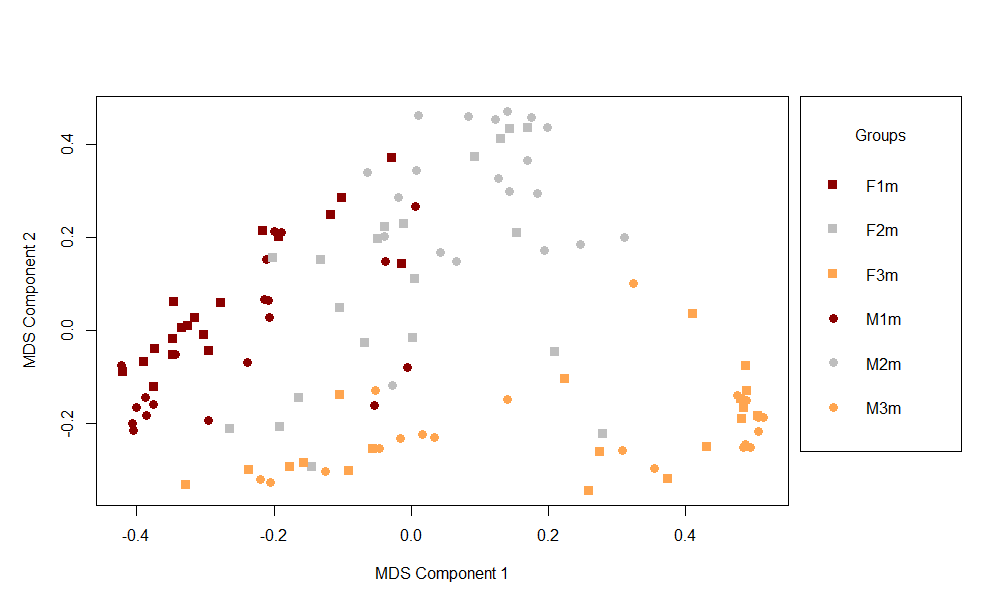
SI Fig. 4 MDS of proximity matrix derived from Random Forest analysis of males from three M and F replicates based on 16 *D. melanogaster* cuticular hydrocarbon peaks that were affected by selection but not time of extraction in linear mixed effect analysis

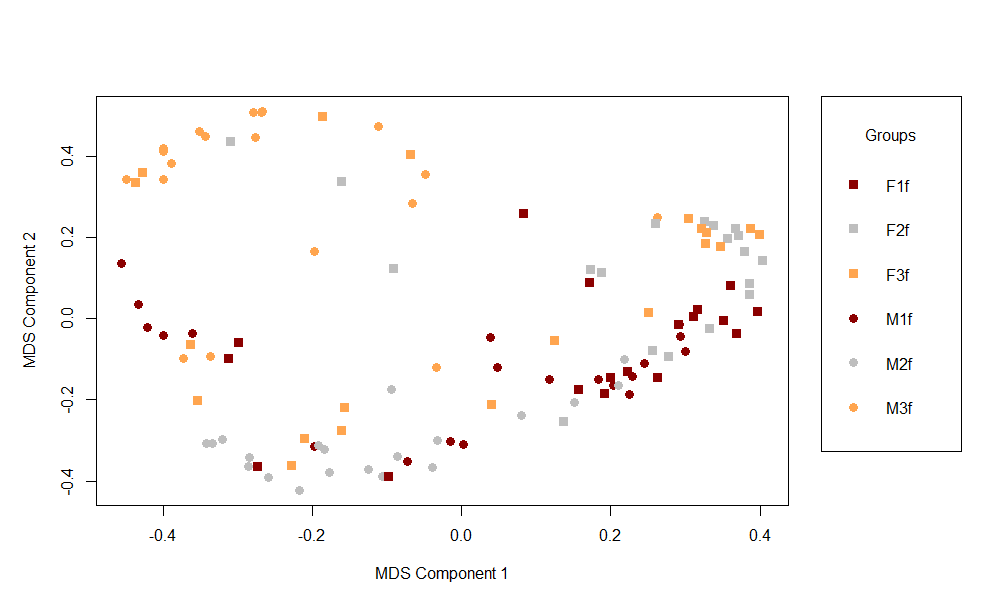
SI Fig. 5 MDS of proximity matrix derived from Random Forest analysis of females from three M and F replicates based on 16 *D. melanogaster* cuticular hydrocarbon peaks that were affected by selection but not time of extraction in linear mixed effect analysis
