## Supplementary material for "Cuticular hydrocarbon divergence in *Drosophila melanogaster* populations evolving under differential operational sex ratios": SI4

Journal Name: Evolutionary Biology

Dutta^1*^, Tejinder Singh Chechi^1^, Ankit Yadav^2^, Nagaraj Guru Prasad^1^

^1^Department of Biological Sciences, Indian Institute of Science Education and Research, Mohali, India

^2^Department of Earth and Environmental Sciences, Indian Institute of Science Education and Research, Mohali, India

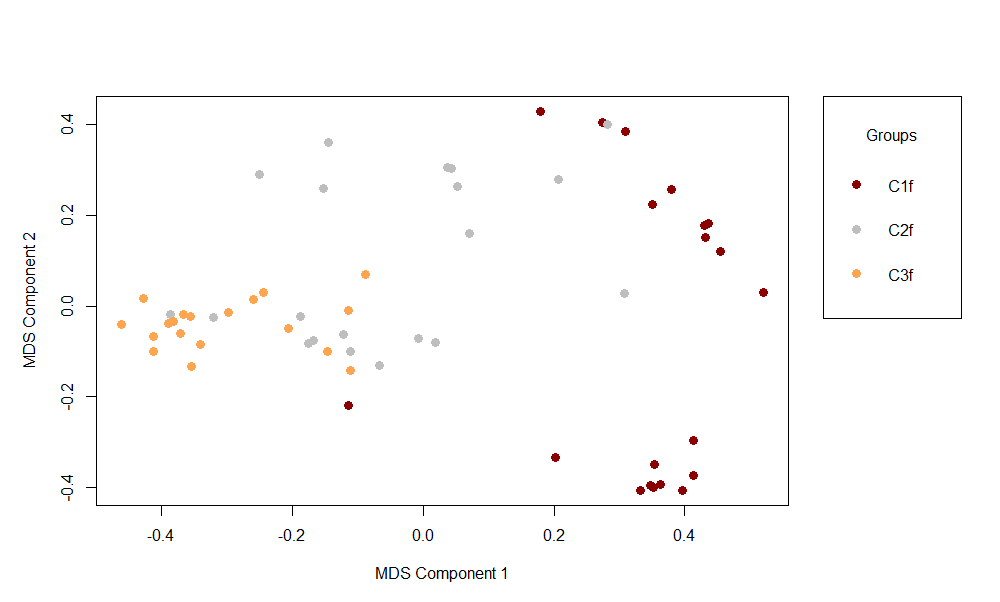
SI Fig. 6 Multi-dimensional scaling plot of proximity matrix derived from Random Forest analysis of females from the three C replicates based on 31 *D. melanogaster* cuticular hydrocarbon peaks

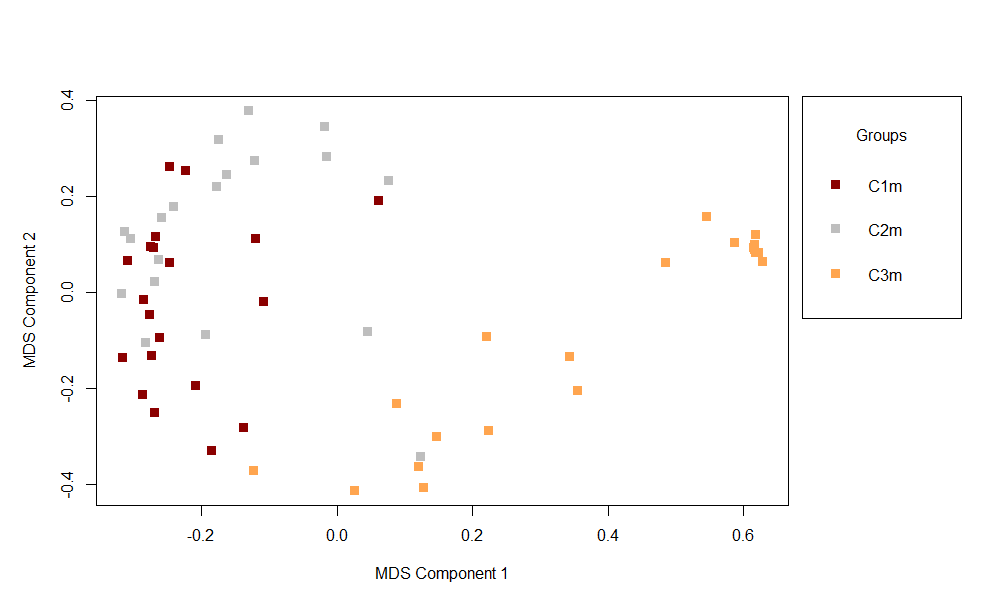
SI Fig. 7 Multi-dimensional scaling plot of proximity matrix derived from Random Forest analysis of males from the three C replicates based on 31 *D. melanogaster* cuticular hydrocarbon peaks
